# The mitochondrial unfolded protein response in human microglia disrupts neuronal-glial communication and promotes senescence

**DOI:** 10.1101/2024.09.03.610925

**Authors:** J. Maria Jose Perez, Alicia Lam, Christin Weissleder, Federico Bertoli, Hariam Raji, Mariella Bosch, Ivan Nemazanyy, Stefanie Kalb, Mohammed Kehili, Insa Hirschberg, Dario Brunetti, Indra Heckenbach, Morten Scheibye-Knudsen, Michela Deleidi

## Abstract

Mitochondria have evolved a specialized mitochondrial unfolded protein response (UPR^mt^) to maintain proteostasis and promote recovery under stress. Studies in simple organisms have shown that UPR^mt^ activation in glial cells supports proteostasis through beneficial noncell-autonomous communication with neurons. However, the role of mitochondrial stress responses in the human brain remains unclear. To address this gap, we investigated the cell type-specific effects of mitochondrial proteotoxic stress using human induced pluripotent stem cell-derived neuronal and glial cultures, as well as brain organoids. We show that mitochondrial proteotoxic stress induces metabolic rewiring in human microglia, marked by depletion of S-adenosylmethionine and lipid remodeling, ultimately leading to a senescent phenotype. Using human neuronal-glial tricultures and microglia-containing brain organoids, we identified the specific contributions of microglia to brain senescence and mitochondrial stress-driven neurodegenerative processes. UPR^mt^ activation disrupts microglial communication with neighboring cells, triggering inflammatory signaling and impairing proteostasis. Together, these findings reveal how impaired mitochondrial proteostasis alters intercellular networks and identify a critical role for the UPR^mt^ in neurodegenerative disease pathogenesis.

## Introduction

Defects in proteostasis and protein aggregation are hallmarks of aging and neurodegenerative diseases. Mitochondria have evolved a specialized unfolded protein response (UPR^mt^) to maintain proteostasis and facilitate recovery under stress by inducing genes encoding mitochondrial chaperones and proteases ^1, 2^. Studies in *Caenorhabditis elegans* (*C. elegans*) have demonstrated that UPR^mt^ activation can extend lifespan through metabolic adaptations and mitochondrial recovery programs ^3, 4^. While these findings highlight protective roles for the UPR^mt^ in simple organisms, their relevance to the mammalian brain remains unclear. Neurons are long-lived, post-mitotic cells that rely on tightly regulated mitochondrial quality control to maintain cellular homeostasis, raising key questions about how mitochondrial proteostasis is controlled in the aging brain and whether UPR^mt^ activation is beneficial or detrimental within neuronal and glial networks.

Age-dependent changes in UPR^mt^ regulation suggest a declining capacity to sense mitochondrial stress. For instance, activity of the mitochondrial protease LONP1 decreases with age, and PITRM1 expression is reduced in the aging brain and in Alzheimer’s disease (AD) patients ^5, 6^. Although UPR^mt^ activation can alleviate neurodegenerative phenotypes ^7, 8, 9^, evidence also indicates that it may contribute to age-related brain disorders ^10, 11^. While initially protective against mitochondrial protein misfolding, chronic UPR^mt^ activation may impair neuronal function and provoke maladaptive immune responses ^12, 13^. Specifically, the UPR^mt^ induces genes linked to mitochondrial repair and innate immunity ^12, 13^, promoting chronic inflammation, a hallmark of neurodegeneration.

In simple organisms, UPR^mt^ signaling can occur in a noncell-autonomous manner, with neurons inducing UPR^mt^ activation in distant tissues. Recent studies in *C. elegans* further show that astrocyte-like glial cells can sense mitochondrial stress and initiate protective signaling that supports neuronal proteostasis ^14^. These findings suggest that mitochondrial stress responses propagate across tissues through integrated intercellular signaling networks ^15–17^. However, how mitochondrial proteotoxic stress is regulated in a cell type-specific manner in the mammalian brain, and how it shapes intercellular communication, remain largely unexplored.

Here, we define the cell type-specific roles of the UPR^mt^ in the human brain. Using human induced pluripotent stem cell (iPSC)-derived monocultures, tricultures, and disease-associated microglia-brain (MgBr) assembloids, we investigate how distinct cell types initiate and transmit mitochondrial proteotoxic stress signals and how these processes impact intercellular communication.

## Results

### Pharmacological inhibition of LONP1 differentially induces UPR^mt^ activation and protein aggregation in human iPSC-derived neurons and glial cells

We differentiated wild-type human iPSCs into cortical neurons, astrocytes, and microglia (Supplementary Figure 1A-H) and treated them with the LONP1 inhibitor C-28 methyl ester of 2-cyano-3,12-dioxoolean-1,9-dien-28-oic acid (CDDO-Me)^18^ to induce mitochondrial protein misfolding and activate the UPR^mt^ (Figure 1A; Supplementary Figure 2A-C). Microglia and astrocytes exhibited early UPR^mt^ activation at 6 hours post-treatment, whereas peak activation in neurons occurred at 24 hours (Figure 1A). LONP1 inhibition caused mitochondrial fragmentation and decreased membrane potential across all cell types (Supplementary Figure 2D). Next, we examined the cell type-specific effects on proteostasis using the PROTEOSTAT assay. Whereas astrocytes and neurons showed no detectable protein accumulation, microglia exhibited a selective increase in mitochondria-associated misfolded proteins after CDDO-Me treatment (Figure 1B). Co-staining of intact cells with PROTEOSTAT and the mitochondrial markers TOMM20 and HSP60 confirmed the preferential association of misfolded proteins with mitochondria in microglia (Supplementary Figure 2E).

**Figure 1.**
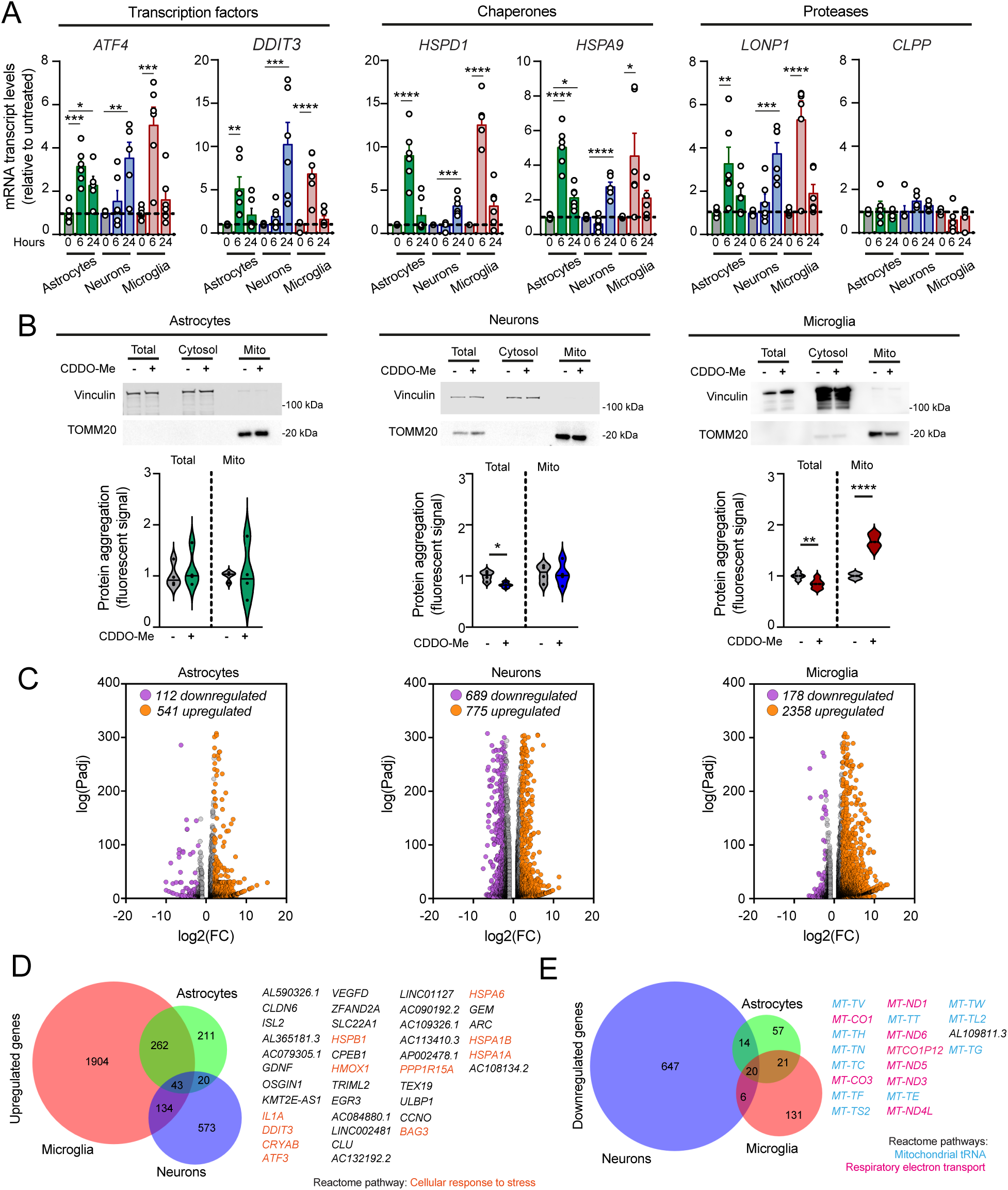
Pharmacological inhibition of LONP1 in human iPSC-derived neurons and glia induces cell type-specific protein misfolding and activation of the UPR^mt^. All experiments were performed using control iPSC line C1. **a,** mRNA expression of UPR^mt^ genes in human iPSC-derived astrocytes, neurons and microglia treated with 1 µM CDDO-Me for 6 h or 24 h. Data are normalized to untreated controls (gray) within each cell type. Mean ± SEM; one-way ANOVA with Bonferroni *post hoc* correction. *ATF4*: \**P* = 0.0229, \*\**P* = 0.004, \*\*\**P* = 0.0003 (astrocytes), and 0.0002 (microglia), *DDIT3*: \*\**P* = 0.0097, \*\*\**P* = 0.0009, and \*\*\*\**P* < 0.0001; *HSPD1*: \*\*\**P* = 0.0003, and \*\*\*\**P* < 0.0001; *HSPA9*: \**P* = 0.0434 (astrocytes), 0.0117 (microglia), and \*\*\*\**P* < 0.0001; *LONP1*: \*\**P* = 0.0083, \*\*\**P* = 0.0002, and \*\*\*\**P* < 0.0001; n = 6 independent experiments. **b,** Upper: representative immunoblots of total, cytosolic (vinculin) and mitochondrial (TOMM20) fractions from neurons, astrocytes and microglia before and after 1 µM CDDO-Me treatment (6 h). Lower: PROTEOSTAT fluorescence intensity in total and mitochondrial fractions. Violin plots show individual values and medians, normalized to untreated controls within each condition. Unpaired two-tailed t test; \**P* = 0.0162, \*\**P* = 0.0060, \*\*\*\**P* < 0.0001. n = 4 (astrocytes, neurons), n = 6 (microglia) independent experiments. **c,** Volcano plots of differentially expressed genes (DEGs) in astrocytes, neurons and microglia under control conditions and after CDDO-Me treatment. Differential expression was determined using DESeq2 with Benjamini–Hochberg correction. Genes with fold change ≥ 2 and FDR ≤ 0.05 are highlighted. **d, e,** Venn diagrams showing shared upregulated (d) and downregulated (e) DEGs across cell types. Transcript isoforms were consolidated to single gene entries for counting. Top enriched Reactome pathways among shared DEGs are indicated.

### LONP1 inhibition elicits divergent stress responses in human iPSC-derived neurons and glia

Bulk RNA-sequencing at peak UPR^mt^ activation (Figure 1C; Supplementary Data 1) revealed robust induction of UPR^mt^-associated genes ^19^ and a broad HSF1-mediated heat shock response across cell types (Figure 1D; Supplementary Figure 3A,B), consistent with qRT-PCR data (Figure 1A) and previous reports in mouse embryonic fibroblasts ^20^. In contrast, downregulated genes were predominantly mitochondrial-encoded (Figure 1E), reflecting reduced mtDNA content and disrupted mitochondrial RNA metabolism in all cell types (Supplementary Figure 3C-G).

Neurons and astrocytes mounted a compensatory proteostatic response marked by increased chaperone expression, activation of the endoplasmic reticulum UPR, enhanced autophagy, and translational repression (Supplementary Figure 3H). In contrast, microglia activated the UPR^mt^ while upregulating components of the translational machinery (Supplementary Figure 3H), a signature previously linked to age-associated integrated stress response (ISR) adaptations ^21, 22, 23, 24^. LONP1 inhibition also induced cell type-specific apoptotic gene programs, with increased *BAX* and *CASP8* and reduced *BCL2* expression in neurons, but elevated *BCL2* in glia (Supplementary Figure 3H, I). Despite these transcriptional changes, no increase in cell death was detected in any cell type (Supplementary Figure 3J).

Pathway-level analyses further revealed distinct stress responses across cell types (Supplementary Figure 4A, B, Supplementary Data 2). Astrocytes preferentially engaged stress-adaptive and innate immune signaling pathways (Supplementary Figure 4A, B). Functionally, this molecular shift did not compromise cellular stability; astrocytes maintained redox homeostasis, as indicated by stable 2, 7-dichlorofluorescein (DCF) levels (Supplementary Figure 4C), and showed no significant change in GFAP protein expression (Supplementary Figure 4D). Neurons exhibited a pronounced oxidative stress response, characterized by the activation of *NFE2L2*-mediated transcriptomic programs (Supplementary Figure 4A, B) and a functional increase in reactive oxygen species (ROS) (Supplementary Figure 4E). While markers of tau and α-synuclein pathology remained unchanged (Supplementary Figure 4F), this stress state resulted in physiological impairment, evidenced by a significant reduction in neuronal excitability and calcium signaling (Supplementary Figure 4G, H). In contrast, microglia displayed a distinct transcriptional profile characterized by senescence-associated remodeling, including epigenetic alterations, changes in nuclear architecture, and activation of senescence-related inflammatory pathways (Supplementary Figure 4A, B). Together, these data indicate that whereas astrocytes adapt and neurons undergo functional impairment without overt degeneration, microglia transition toward a dysfunctional, senescence-like state in response to mitochondrial proteotoxic stress.

### LONP1 inhibition induces a senescent phenotype in human iPSC-derived microglia

To examine the relationship between mitochondrial proteotoxic stress and microglial senescence, we performed gene set enrichment analysis using eleven published senescence-associated gene signatures ^25, 26, 27, 28^. CDDO-Me-treated iPSC-derived microglia and neurons showed significant positive enrichment of gene sets upregulated in senescence and negative enrichment of those typically downregulated (Figure 2A, Supplementary Figure 5A). qRT-PCR and immunostaining revealed that only CDDO-Me-treated microglia upregulated *CDKN2A* (p16) and *CDKN1A* (p21), together with increased expression and secretion of senescence-associated secretory phenotype (SASP) cytokines, including IL-1β, IL-6, IL-8, and TNF-α (Figure 2B-D; Supplementary Figure 5B-D). These findings were independently validated in two additional iPSC lines (Supplementary Figure 5E-I).

**Figure 2.**
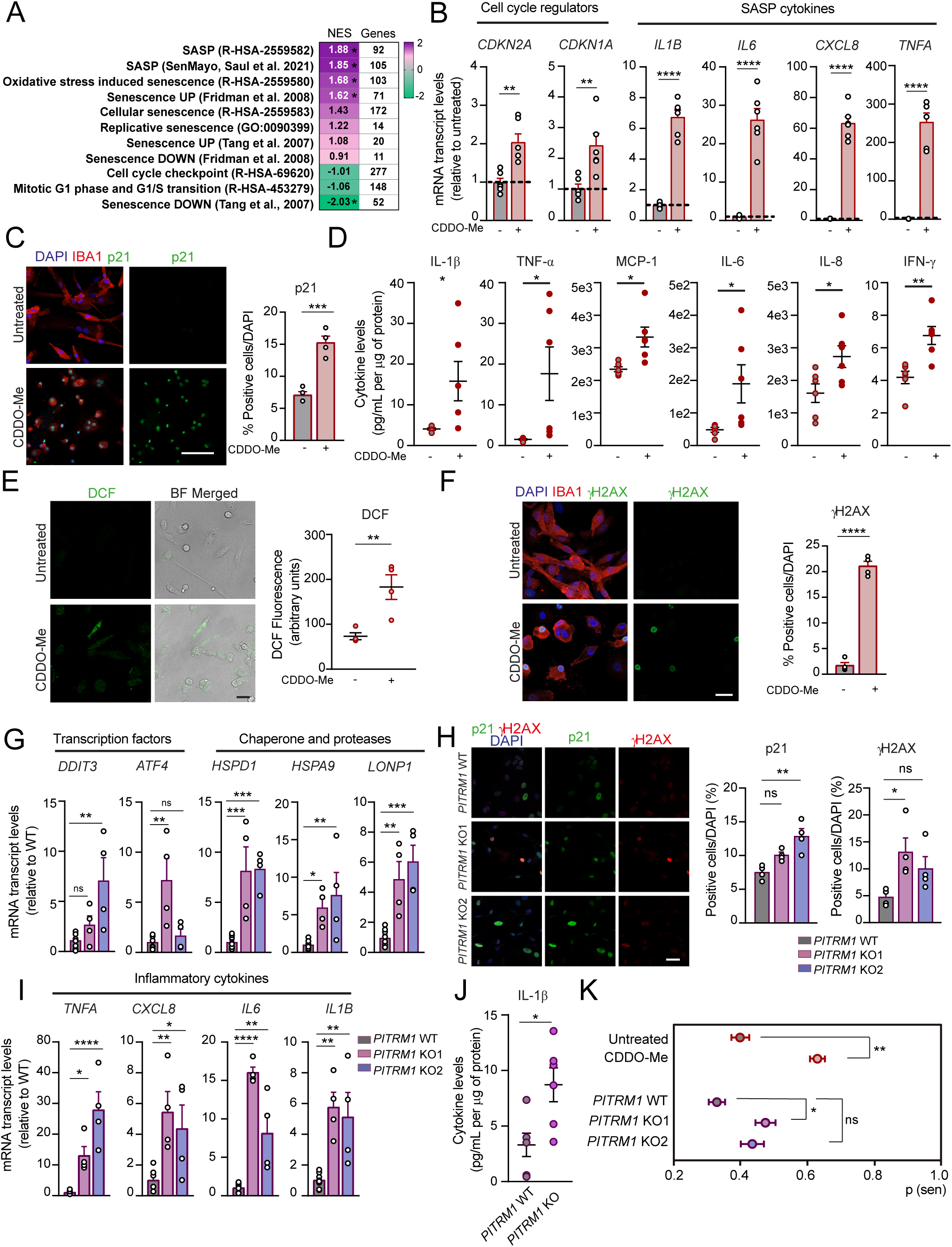
Mitochondrial proteotoxic stress induces a senescent phenotype in human iPSC-derived microglia. Microglial senescence was examined in pharmacological (CDDO-Me treatment in line C1; a-f) and genetic (*PITRM1* knockout; g-k) models associated with UPR^mt^ activation. **a,** Heat map showing enrichment of senescence-associated gene sets in RNA-sequencing data from microglia treated with CDDO-Me. Normalized enrichment scores (NES) were calculated by GSEA relative to untreated controls. Upregulation is shown in purple and downregulation in green. *FDR ≤ 0.05. **b,** mRNA expression of *CDKN2A*, *CDKN1A*, and SASP cytokines (*IL1B*, *IL6*, *CXCL8*, *TNFA*) following CDDO-Me treatment. Data are normalized to untreated controls (gray). Mean ± SEM; unpaired two-tailed *t* test. *CDKN2A* \*\**P* = 0.0036; *CDKN1A* \*\**P* = 0.0016; *IL1B, IL6*, *CXCL8*, *TNFA*, \*\*\*\**P* < 0.0001; n = 6 independent experiments. **c,** Representative confocal images of p21 (green) in human iPSC-derived microglia (IBA1, red; DAPI, blue). Scale bar, 50 µm. Right: quantification of the percentages of p21^+^/DAPI^+^. Mean ± SEM; unpaired two-tailed *t* test; \*\*\**P* = 0.0004; n = 4 independent experiments. **d,** Cytokine levels in supernatants from untreated or CDDO-Me-treated (6 h) microglia. Mean ± SEM; unpaired two-tailed *t* test. \**P* = 0.0356 (IL-1β), 0.0336 (TNF-α), 0.0105 (MCP-1), 0.0381 (IL-6), 0.0283 (IL-8), and \*\**P* = 0.0033; n = 6 independent experiments. **e,** Representative confocal images of DCF-based ROS measurements in control and CDDO-Me-treated microglia. Scale bar, 10 µm. Right: quantification of DCF fluorescence intensity. Mean ± SEM; unpaired two-tailed *t* test; ** *P* = 0.0085. n = 4 independent experiments. **f,** Representative γH2AX immunostaining in control and CDDO-Me-treated microglia (γH2AX, green; IBA1, red; DAPI, blue). Scale bar, 10 µm. Right: quantification of γH2AX⁺/DAPI⁺ cells. Mean ± SEM; unpaired two-tailed *t* test; **** *P* < 0.0001; n = 4 independent experiments. **g,** mRNA expression of UPR^mt^ genes in *PITRM1*-WT and two independent *PITRM1*-KO clones (KO1, KO2). Data are normalized to WT controls (gray). Mean ± SEM; one-way ANOVA with Bonferroni *post hoc* correction. *DDIT3*, \*\**P* = 0.0032; *ATF4*, \*\**P* = 0.0013; *HSPD1*, \*\*\**P* = 0.0010 (KO1), 0.0008 (KO2); *HSPA9,* \**P* = 0.0484, \*\**P* = 0.0092; *LONP1,* \*\**P* = 0.0030, \*\*\**P* = 0.0003; ns = not significant; n = 4 independent biological replicates per KO clone and its matched WT. **h,** Representative p21 (green) and γH2AX (red) staining in *PITRM1*-WT and *PITRM1*-KO microglia (DAPI, blue). Scale bar, 10 µm. Right: quantification of the percentage of p21⁺/DAPI⁺ and γH2AX⁺/DAPI⁺. Mean ± SEM; one-way ANOVA with Bonferroni *post hoc* correction. \**P* = 0.0338, \*\**P* = 0.0014; ns = not significant. n = 4 independent experiments. **i,** SASP gene expression (*TNFA*, *CXCL8, IL6*, *IL1B*) in *PITRM1*-WT and KO microglia (KO1 and KO2). Mean ± SEM; normalized to WT controls (gray); one-way ANOVA with Bonferroni *post hoc* correction. *TNFA*, \**P* = 0.0190, \*\*\*\**P* < 0.0001; *CXCL8,* \**P* = 0.0373; \*\**P* = 0.0072; *IL6*, \*\**P* = 0.0010, \*\*\*\**P* < 0.0001; *IL1B*, \*\**P* = 0.0020 (KO1), 0.0059 (KO2); n = 4 independent biological replicates per KO clone and its matched WT. **j,** IL-1β levels in supernatants of *PITRM1*-WT and *PITRM1*-KO2 microglia. Mean ± SEM; unpaired two-tailed *t* test; \**P* = 0.0152; n = 6 independent experiments. **k,** Predicted senescence scores in untreated, CDDO-Me-treated, *PITRM1*-WT, and *PITRM1*-KO human iPSC-derived microglia. Mean ± SEM; unpaired two-tailed *t* test (CDDO-Me) and one-way ANOVA with Bonferroni post *hoc* correction (*PITRM1*); \**P* = 0.0258, \*\**P* = 0.0078; ns = not significant; n = 4 independent experiments.

Flow cytometric analysis following Hoechst 33342 staining showed that astrocytes underwent a modest redistribution from G1 to S phase, consistent with a stress-induced reactive state (Supplementary Figure 6A-C). In contrast, iPSC-derived microglia and neurons exhibited a largely nonproliferative profile with minimal G2 representation that was unchanged by treatment with CDDO-Me or the senescence inducer etoposide (Supplementary Figure 6D-E), indicating that iPSC-derived microglia are largely terminally differentiated ^29^ and mitochondrial stress induces senescence without cell cycle re-entry. Additional hallmarks confirmed the senescent phenotype. SA-β-gal activity, already detectable under basal conditions^30^, was increased following CDDO-Me treatment (Supplementary Figure 6F). This was accompanied by elevated ROS production, γH2AX-positive nuclei (Figure 2E, F), increased NAD⁺ consumption, and enhanced PARP1 activity (Supplementary Figure 6G, H), consistent with DNA damage-associated senescence ^31, 32, 33, 34^.

To exclude off-target effects, *LONP1* was silenced using shRNA (Supplementary Figure 7A). Knockdown recapitulated p21 induction, γH2AX accumulation, and increased IL-8 secretion (Supplementary Figure 7B,C). Similarly, pharmacological UPR^mt^ activation with gamitrinib-TPP (GTPP) ^35^ or *ATF5* overexpression induced senescence markers and SASP gene expression (Supplementary Figure 7D-G). Together, these data demonstrate that mitochondrial proteotoxic stress and UPR^mt^ activation are sufficient to drive a selective senescence-like program in human microglia.

### Mitochondrial proteotoxic stress induces a transcriptional state associated with early pathological microglial activation

Although senescence markers normalized following CDDO-Me withdrawal (Supplementary Figure 8A, B), senescence-associated stress can imprint long-lasting molecular changes ^29^. To test whether mitochondrial stress alters immune reactivity, we challenged CDDO-Me-pretreated iPSC-derived microglia with LPS. Pretreated cells released more IL-1β but less TNF-α and IL-6 (Supplementary Figure 8C), consistent with age-associated immune remodeling and persistent inflammatory reprogramming ^36, 37^.

To assess disease relevance, we compared the transcriptional profile of CDDO-Me-treated microglia with human microglial clusters from the Human Microglia Atlas (HuMicA) ^38^. Enrichment analysis revealed significant overlap with early pathological states linked to inflammation, amyloid pathology, and cellular stress in aging and neurodegeneration (Supplementary Figure 8D, E; Supplementary Data 1). The strongest similarities were observed for programs marked by immediate early genes, pro-inflammatory signaling, and heat shock responses, reported in AD and amyloid-exposed microglia^39^. The CDDO-Me signature also overlapped with phagocytic and phospho-tau-associated microglial states, while diverging from homeostatic and proliferative programs ^39–41^ (Supplementary Figure 8D). Visualization of the top intersecting genes showed reduced microglial identity alongside activation of inflammatory, UPR, and heat shock pathways (Supplementary Figure 8E). Partial overlap with disease-inflammatory macrophages ^42^ further supports a maladaptive stress trajectory. Together, these data indicate that acute mitochondrial proteotoxic stress primes microglia toward an early pathological activation state characteristic of human aging and neurodegeneration.

### Disease-associated defects in mitochondrial proteostasis induce a senescence-like phenotype in human iPSC-derived microglia

To validate the disease relevance of our findings, we examined models deficient in *PITRM1*, a mitochondrial metallopeptidase that degrades mitochondrial targeting sequences after their cleavage from proteins imported into mitochondria ^43^. Its impairment activates the UPR^mt^ in yeast, human iPSC-derived neurons, and brain organoids ^8, 44^, while loss-of-function mutations cause a slowly progressive autosomal recessive neurodegenerative disorder with cognitive decline ^45, 46^. PITRM1 also contributes to amyloid-β (Aβ) clearance, and its dysfunction results in Aβ accumulation ^8, 45, 47, 48^. Accordingly, *Pitrm1*⁺/⁻ mice develop progressive neurological symptoms and brain Aβ_1_42_ accumulation resembling AD pathology ^45^. Notably, *Pitrm1*⁺/⁻ mice displayed an age-dependent increase in SA-β-gal activity in the cortex and hippocampus. At 15 months, this was accompanied by elevated number of p21-positive microglia and TNF-α levels (Supplementary Figure 9A-D), consistent with microglial senescence.

To assess cell-autonomous effects in human cells, we generated *PITRM1* wild-type (WT) and knockout (KO) iPSC-derived microglia. *PITRM1* deficiency did not impair differentiation, as shown by comparable numbers of IBA1- and PU.1-positive cells (Supplementary Figure 9E), but robustly activated the UPR^mt^ (Figure 2G). *PITRM1*-KO microglia upregulated *CDKN2A* (p16) and *CDKN1A* (p21) at transcript and protein levels (Figure 2H; Supplementary Figure 9F), displayed increased SASP cytokine expression (*IL1B*, *IL6*, *CXCL8*, *TNFA*) and IL-1β secretion, and showed elevated oxidative stress, γH2AX-positive nuclei, and NAD⁺ consumption (Figure 2I, J; Supplementary Figure 9G-I). In contrast, *PITRM1*-deficient astrocytes and neurons did not exhibit increased p21 expression or DNA damage (Supplementary Figure 9J). Together, these findings demonstrate that genetic defects in mitochondrial proteostasis associated with human neurodegenerative disease selectively drive a senescence-like program in microglia.

### Morphology-based deep learning identifies UPR^mt^-driven senescence in human iPSC-derived microglia

To address the challenge of detecting senescence in non-dividing cells ^49, 50^, we leveraged characteristic nuclear features of senescent cells-including increased nuclear area, reduced convexity, and altered aspect ratio-and applied a machine learning-based classifier to identify senescent microglia. This approach has been validated to distinguish senescence from quiescence ^51^. Both CDDO-Me-treated and *PITRM1*-KO microglia exhibited significantly higher predicted senescence scores compared with controls (Figure 2K). Notably, CDDO-Me-treated microglia showed higher senescence scores, larger nuclear area, and reduced convexity and aspect ratio relative to *PITRM1*-KO cells, consistent with a more uniform and pronounced senescence phenotype induced by acute pharmacological stress.

### UPR^mt^ activation dysregulates lipid metabolism in human iPSC-derived microglia

To explore the link between mitochondrial stress responses, lipid metabolism, and senescence ^14, 52, 53^, we performed targeted mass spectrometry-based metabolic profiling in control and CDDO-Me-treated iPSC-derived microglia (Supplementary Data 3). Principal component analysis (PCA) and hierarchical clustering revealed marked metabolic shifts upon UPR^mt^ activation (Figure 3A, Supplementary Figure 10A).

**Figure 3.**
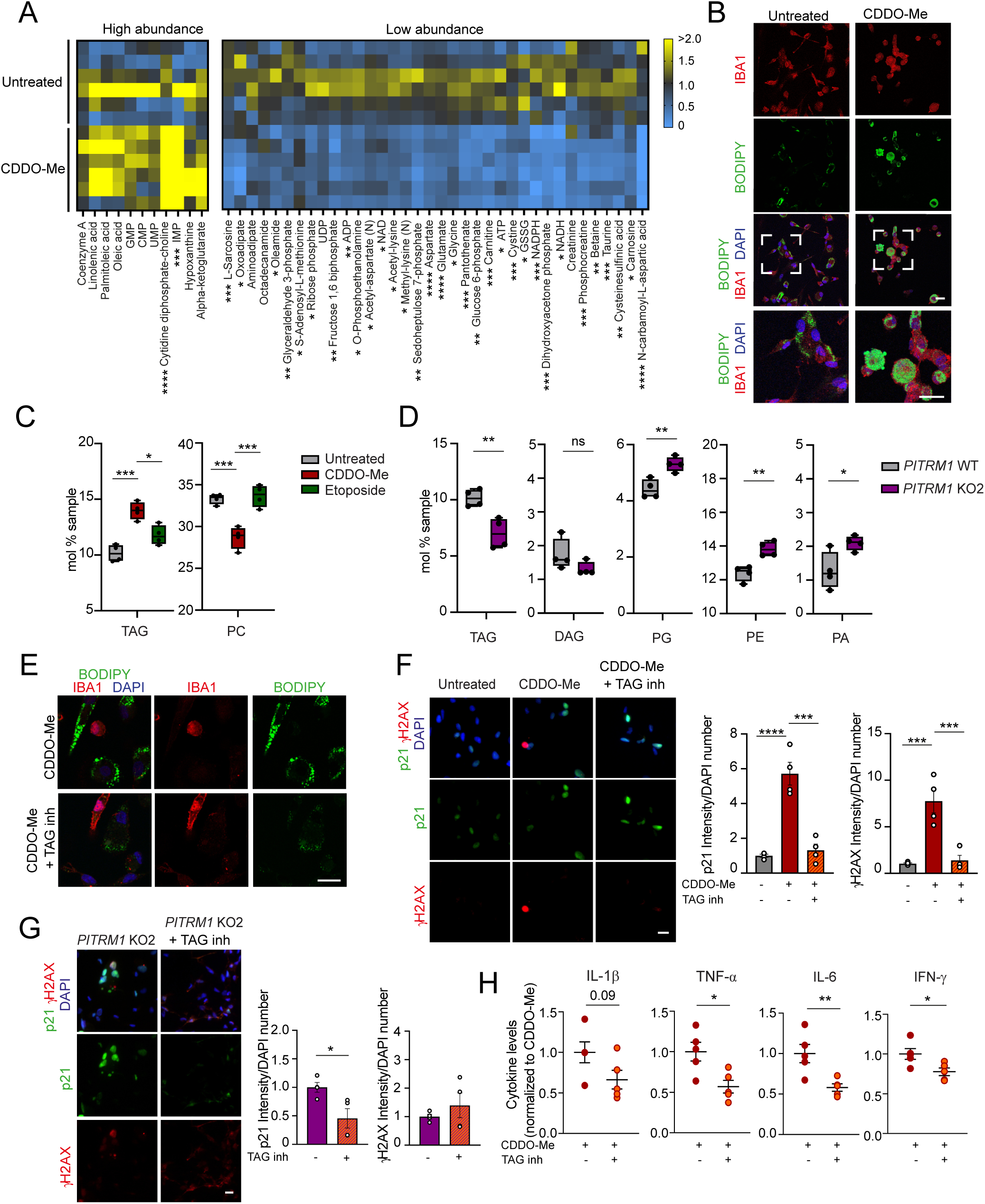
UPR^mt^ dysrupts lipid metabolism in human iPSC-derived microglia. All experiments were performed using iPSC line C2 unless otherwise indicated. **a,** Heat map of the top most significantly altered metabolites in control and CDDO-Me-treated microglia. Metabolites are ranked according to relative abundance in CDDO-Me-treated cells compared to controls (higher abundance shown on the left; lower abundance on the right). Yellow indicates increased and blue decreased metabolite levels. Data are normalized to control; unpaired two-tailed *t* test;\**P* = 0.0366 (oxoadipate), 0.0351 (oleamide), 0.0212 (S-adenosyl-L-methionine), 0.0290 (ribose phosphate), 0.0269 (O-phosphoethanolamine), 0.0186 (acetyl-aspartate), 0.02 (NAD), 0.0161 (acetyl-lysine), 0.0206 (methyl-lysine), 0.0102 (glycine), 0.0176 (ATP), 0.0270 (GSSG), 0.0107 (NADH), 0.0175 (carnosine); \*\**P* = 0.0046 (glyceraldehyde-3-phosphate), 0.0030 (fructose-1,6-biphosphate), 0.0098 (ADP), 0.0016 (sedoheptulose-7-phosphate), 0.0010 (glucose-6-phosphate), 0.0017 (betaine), 0.0030 (cysteinesulfinic acid); \*\*\**P* = 0.0003 (IMP), 0.0007 (L-sarcosine), 0.0009 (pantothenate), 0.005 (carnitine), 0.0002 (cystine), 0.0006 (NADPH), 0.0003 (dihydroxyacetone-phosphate), 0.0002 (phosphocreatine), 0.0009 (taurine); \*\*\*\**P* <0.0001; n = 6 independent experiments. **b,** Representative confocal images of IBA1 (red), BODIPY (green), and DAPI (blue) staining in microglia. Scale bar, 10 µm. Images are representative of at least two independent experiments. **c,** LC–MS quantification of triacylglycerol (TAG) and phosphatidylcholine (PC) (mol%) in untreated, CDDO-Me-treated, and etoposide-treated microglia. Center line, median; box limits, 25th–75th percentiles; whiskers, minimum to maximum. One-way ANOVA with Bonferroni *post hoc* correction. TAG: \**P* = 0.0118, \*\*\**P* = 0.0003; PC, \*\*\**P* = 0.0008 (untreated vs CDDO-Me) and 0.0005 (CDDO-Me vs etoposide); n=4 independent experiments. **d,** LC–MS analysis of TAG, diacylglycerol (DAG), phosphatidylglycerol (PG), phosphatidylethanolamine (PE), and phosphatidic acid (PA) (mol%) in *PITRM1*-WT and *PITRM1*-KO2 microglia. Center line, median; box limits, 25th–75th percentiles; whiskers, minimum to maximum. Unpaired two-tailed *t*-test. TAG, \*\**P* = 0.0063; PG, \*\**P* =0.0060; PE, \*\**P* = 0.0037; PA \**P* = 0.0320; ns = not significant; n=4 independent experiments. **e,** Representative confocal images of IBA1 (red), BODIPY (green), and DAPI (blue) staining in microglia treated with CDDO-Me with or without TAG inhibitors (5 µM). Scale bar, 10 µm. Representative of at least two independent experiments.**f,** Representative images of p21 (green) and γH2AX (red) staining in untreated, CDDO-Me-treated, and CDDO-Me + TAG inhibitors (5 µM) conditions. DAPI (blue). Scale bar, 10 µm. Right: quantification of p21 and γH2AX fluorescence intensity in DAPI+ cells, normalized to control. Mean ± SEM; one-way ANOVA with Bonferroni *post hoc* correction. P21, \*\*\**P* = 0.0001, \*\*\*\**P* <0.0001; γH2AX, \*\*\**P* = 0.0006 (untreated vs CDDO-Me), 0.0009 (CDDO-Me vs CDDO-Me + TAG inhibitors); n = 4 independent experiments. **g,** Representative images of p21 (green) and γH2AX (red) staining in *PITRM1*-KO2 microglia with or without TAG inhibitors (10 µM). DAPI (blue). Scale bar, 10 µm. Right: quantification normalized to *PITRM1*-KO2. Mean ± SEM; unpaired two-tailed t test; \**P* = 0.0290; n = 4 independent experiments. **h,** Cytokine levels in supernatants from CDDO-Me-treated, and CDDO-Me + TAG inhibitors-treated microglia, normalized to CDDO-Me-treated. Mean ± SEM; unpaired two-tailed *t* test. TNF-α, \**P* = 0.0152; IL-6, \*\**P* = 0.0092; IFN-γ, \**P* = 0.0267; n = 5 independent experiments.

Integration of transcriptomic and metabolomic data identified enrichment of lipid pathways, including glycerophospholipid and glycerolipid metabolism, with decreased dihydroxyacetone phosphate and O-phosphoethanolamine and increased CDP-choline (Figure 3A, Supplementary Figure 10B) ^54^. Twenty lipid metabolism genes were dysregulated, affecting phosphatidylcholine, phosphatidate, phosphatidylethanolamine, phosphatidylserine, and CDP-diacylglycerol (DAG) pathways (Supplementary Figure 10B).

BODIPY staining showed altered lipid droplet (LD) distribution in CDDO-Me-treated microglia (Figure 3B). Lipidomics revealed decreased phosphatidylcholine alongside triacylglycerol (TAG) accumulation, consistent with a shift toward lipid storage (Figure 3C, Supplementary Figure 10C), whereas *PITRM1*-KO microglia exhibited reduced TAG and DAG, and increased membrane phospholipids (Figure 3D, Supplementary Figure 10D). Inhibition of TAG synthesis via DGAT1/2 reduced LD formation and senescence markers (p21, γH2AX) in both models (Figure 3E-G, Supplementary Figure 10E), and decreased TNF-α, IL-6 and IFN-γ secretion by CDDO-Me-treated microglia (Figure 3H), indicating that TAG synthesis is required for UPR^mt^-induced senescence.

### UPR^mt^ activation dysregulates methionine metabolism in human iPSC-derived microglia

We performed metabolite set enrichment analysis to identify metabolic pathways altered in CDDO-Me-treated human iPSC-derived microglia. Amino acid metabolism pathways were significantly enriched, including the malate-aspartate shuttle, lysine degradation, and β-alanine metabolism (Supplementary Figure 11A). Folate and cysteine metabolism were markedly dysregulated, with decreased levels of glycine, sarcosine, cysteine sulfinic acid, betaine, taurine, cystine, and S-adenosylmethionine (SAM) (Figures 3A and 4A).

**Figure 4.**
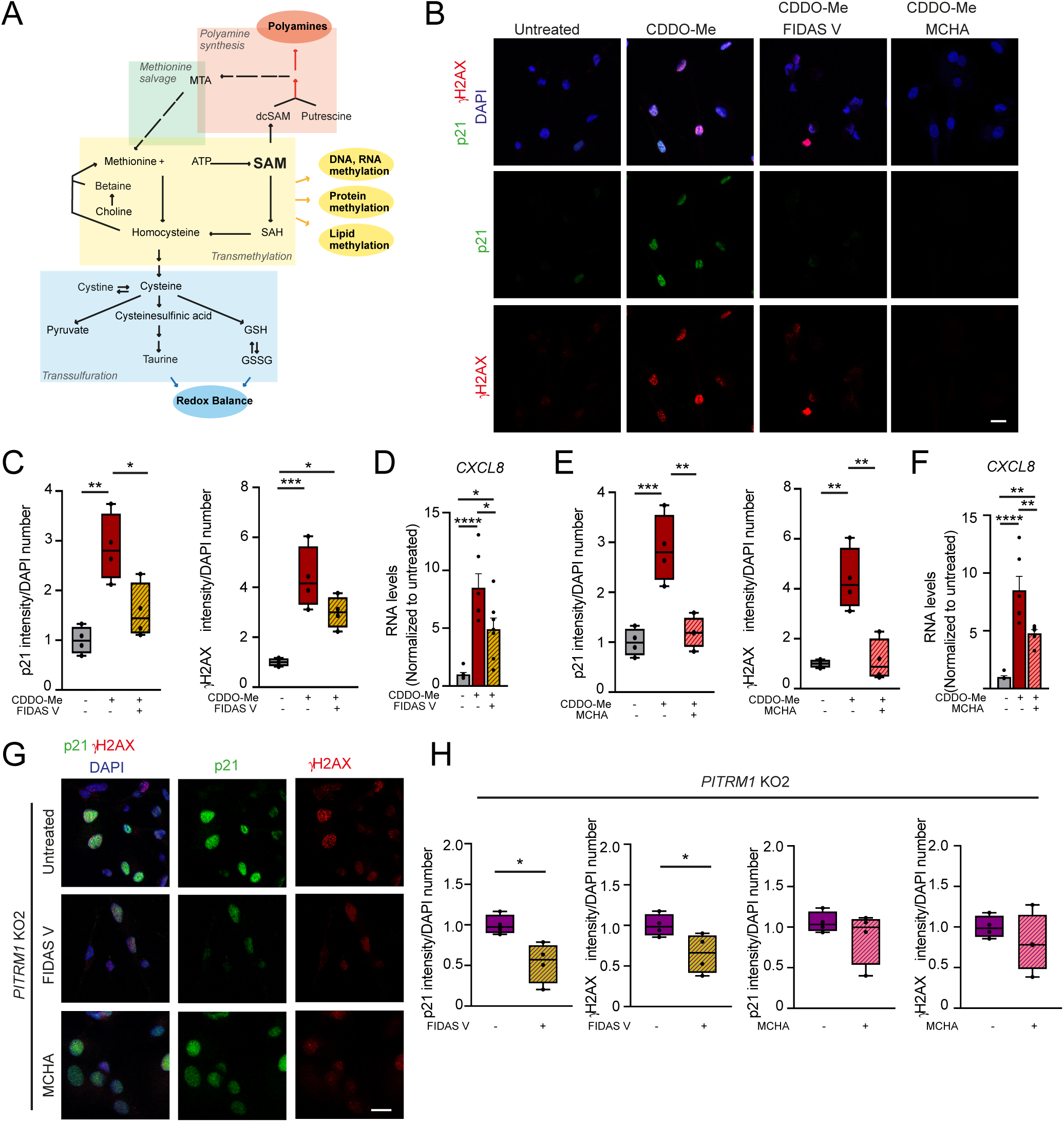
Reducing SAM availability attenuates the senescent phenotype in human iPSC-derived microglia. All experiments were performed using iPSC line C2, unless otherwise indicated. **a,** Schematic of key metabolites in methionine and cysteine metabolism, including transmethylation, methionine salvage, polyamine synthesis, and transsulfuration pathways. **b,** Representative confocal images of p21 (green) and γH2AX (red) in microglia untreated or treated with 1 µM CDDO-Me alone or combined with 25 µM FIDAS V or 250 µM MCHA for 6 h. Nuclei were stained with DAPI (blue). Scale bar, 10 µm. **c,** Quantification of p21^+^/DAPI^+^ and γH2AX^+^/DAPI^+^ fluorescence in untreated, CDDO-Me-treated, and CDDO-Me + FIDAS V groups. Data normalized to untreated controls. Center line, median; box, 25th–75th percentiles; whiskers, min–max. One-way ANOVA with Bonferroni *post hoc* correction. P21, \**P* = 0.0223, \*\**P* = 0.0023; γH2AX, \**P* = 0.0209; \*\*\**P* = 0.0007; n = 4 independent experiments. **d,** *CXCL8* mRNA levels under cotreatment conditions. Data normalized to untreated controls, mean ± SEM; one-way ANOVA with Bonferroni *post hoc* correction; \**P* = 0.0133 (CDDO-Me versus CDDO-Me + FIDAS V), 0.0305 (untreated versus CDDO-Me + FIDAS V); \*\*\*\**P* < 0.0001; n = 7 independent experiments. **e,** Quantification of p21^+^/DAPI^+^ and γH2AX^+^/DAPI^+^ fluorescence in untreated, CDDO-Me-treated, and CDDO-Me + MCHA groups. Data normalized to untreated controls. Center line, median; box, 25th–75th percentiles; whiskers, min–max. One-way ANOVA with Bonferroni *post hoc* correction. P21, \*\**P* = 0.0018, \*\*\**P* = 0.0008; γH2AX \*\**P* = 0.0012 (untreated versus CDDO-Me), 0.0016 (CDDO-Me versus CDDO-Me + MCHA); n = 4 independent experiments. **f,** *CXCL8* mRNA levels in cotreated cells. Mean ± SEM; Data normalized to untreated controls, mean ± SEM; one-way ANOVA with Bonferroni *post hoc* correction; **P = 0.0069 (untreated versus CDDO-Me + MCHA), 0.0078 (CDDO-Me versus CDDO-Me + MCHA), and ****P < 0.0001; n = 6 independent experiments. **g,** Representative confocal images of p21 (green) and γH2AX (red) in *PITRM1*-KO2 microglia, untreated or treated with 25 µM FIDAS V or 250 µM MCHA for 6 h. Nuclei: DAPI (blue). Scale bar, 10 µm. **h,** Quantification of p21^+^/DAPI^+^ and γH2AX^+^/DAPI^+^ fluorescence in *PITRM1*-KO (KO2) microglia under indicated treatments. Data normalized to untreated controls. Center line, median; box, 25th–75th percentiles; whiskers, min–max. Unpaired two-tailed t-test, \**P* = 0.0147 (p21), 0.0460 (γH2AX); n = 4 independent experiments.

Consistent with impaired glutathione synthesis, total GSH levels were decreased, although the GSH/GSSG ratio was increased (Supplementary Figure 11B), suggesting compensatory redox adaptation. Transcriptomic analysis revealed upregulation of methionine adenosyltransferase II (*MAT2A*), which catalyzes SAM synthesis from methionine and ATP, and cystathionine β-synthase (*CBS*), a key regulator of homocysteine metabolism (Supplementary Data 1). These data support coordinated rewiring of methionine and redox metabolism during UPR^mt^ activation and link these changes to microglial senescence.

To test the contribution of oxidative stress, we treated cells with the mitochondria-targeted antioxidant mitoquinone mesylate (MitoQ). MitoQ reduced p21 and DNA damage and restored *HSPD1*, *DDIT3* and *CXCL8* expression to baseline in CDDO-Me-treated microglia (Supplementary Figure 11C, D), consistent with ROS acting upstream of UPR^mt^ ^55^. However, MitoQ failed to rescue p21 or γH2AX levels in *PITRM1*-KO microglia (Supplementary Figure 11E), indicating that antioxidant treatment alone is insufficient to reverse the senescence phenotype in the context of persistent mitochondrial proteostasis defects.

Metabolic profiling of *PITRM1*-KO microglia revealed similar disruption of methionine, betaine, and taurine pathways (Supplementary Figure 11F, G; Supplementary Data 3). Both CDDO-Me-treated and *PITRM1*-KO microglia exhibited significant reductions in L-sarcosine, cystine, and SAM levels (Figure 4A; Supplementary Figure 11F). Together, these data identify impaired methionine metabolism as a shared metabolic signature of UPR^mt^-driven microglial senescence across pharmacological and genetic models.

### Targeting SAM metabolism attenuates the senescence phenotype in human iPSC-derived microglia

To determine whether methionine metabolism contributes to UPR^mt^-driven senescence, we modulated SAM availability in CDDO-Me-treated human iPSC-derived microglia. Supplementation with SAM-HCl increased DNA damage and p21 immunofluorescence compared with CDDO-Me alone (Supplementary Figure 12A-C). SAM-HCl also elevated *HSPD1* transcripts without affecting *DDIT3* and enhanced *CXCL8* expression (Supplementary Figure 12D), indicating partial UPR^mt^ activation together with amplification of the SASP.

Given the upregulation of *MAT2A* and reduced SAM levels upon CDDO-Me treatment, we hypothesized that increased SAM flux into downstream pathways, including polyamine synthesis, contributes to senescence. Inhibition of SAM synthesis with the MAT2A inhibitor FIDAS V significantly reduced p21 protein levels and *CXCL8* transcripts in CDDO-Me-treated microglia (Figure 4B-D; Supplementary Figure 12E). Similarly, blockade of spermidine synthesis with MCHA decreased DNA damage, p21, and *CXCL8* expression (Figure 4B, E-F). Neither FIDAS V nor MCHA altered UPR^mt^ markers (*HSPD1, DDIT3*) (Supplementary Figure 12F), indicating that attenuation of senescence occurred downstream of UPR^mt^ activation. In *PITRM1*-KO microglia, FIDAS V reduced p21 and DNA damage, whereas MCHA showed a similar trend (Figure 4G, H; Supplementary Figure 12G, H). Although TAG inhibition did not restore SAM levels, both FIDAS V and MCHA reduced LD accumulation (Supplementary Figure 12I, J). Together, these findings suggest that SAM availability regulates lipid remodeling and represents a key metabolic regulator of UPR^mt^-driven microglial senescence.

### UPR^mt^ activation induces a microglia-specific senescence phenotype in iPSC-derived tricultures

To examine whether microglial senescence persists in a multicellular context, we established a human iPSC-derived triculture system composed of 30% neurons, 60% astrocytes, and 10% microglia (Figure 5A). Cultures were maintained for 7 days prior to treatment with 1 µM CDDO-Me for 6 hours. Cell type-specific senescence was quantified using nuclear morphology-based deep learning analysis (Figure 5B). Consistent with monoculture experiments, CDDO-Me treatment selectively increased the predicted senescence score in microglia, whereas neurons and astrocytes showed reduced scores compared to controls (Figure 5B), indicating a microglia-specific senescence response within the triculture environment. To validate these findings in a disease-relevant context, we analyzed *PITRM1*-KO tricultures. As observed following pharmacological induction, *PITRM1*-deficient microglia displayed significantly higher predicted senescence scores compared to neurons or astrocytes (Supplementary Figure 13A). Together, these data demonstrate that mitochondrial proteostasis defects and UPR^mt^ activation drive a robust and cell type-specific senescence program in human iPSC-derived microglia, even within a complex multicellular milieu.

**Figure 5.**
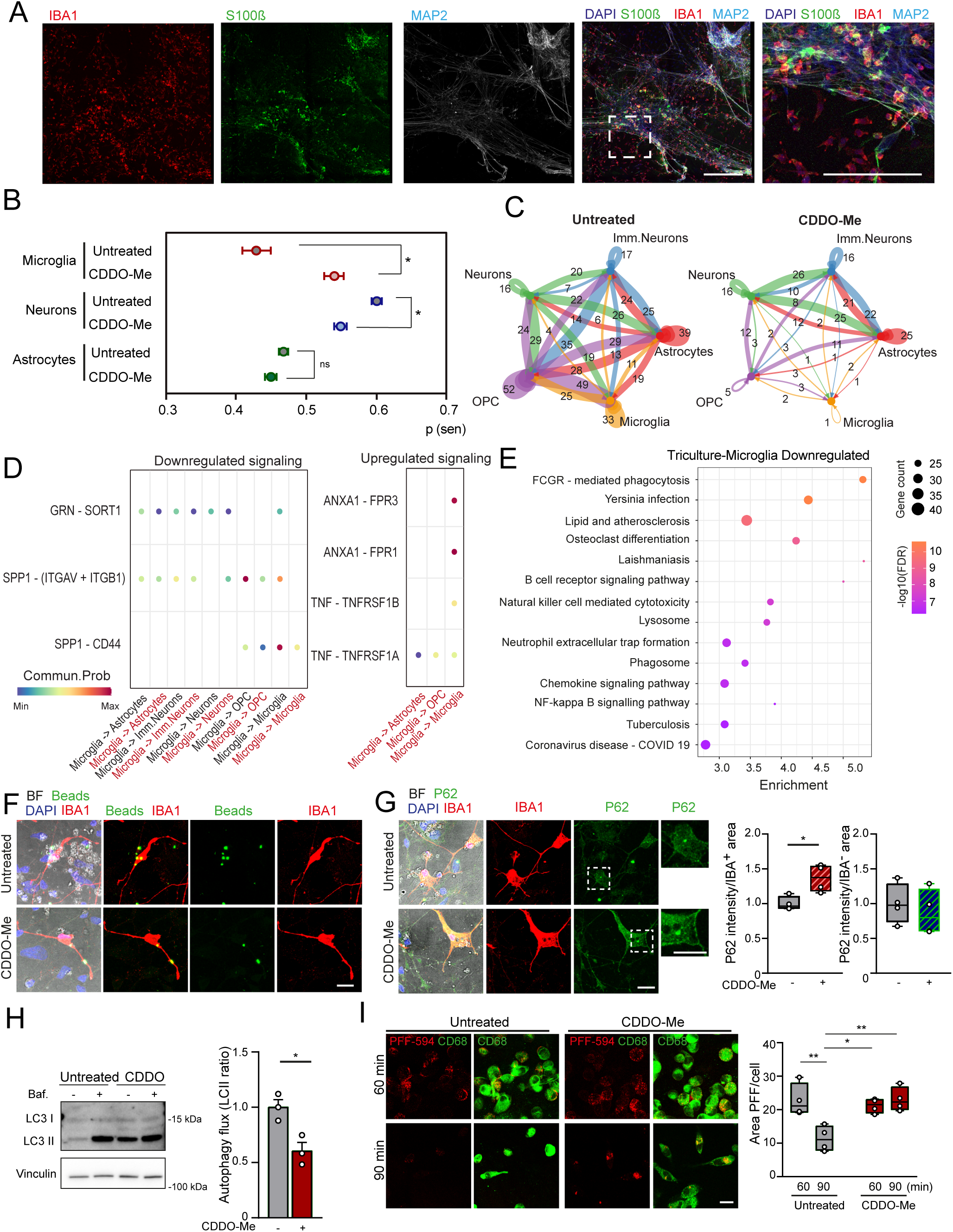
Mitochondrial proteotoxic stress impairs cell-cell communication and misfolded protein clearance in human iPSC-derived microglia in mono- and tricultures. All experiments were performed using iPSC line C2. **a,** Representative confocal images of control iPSC-derived tricultures showing microglia (IBA1, red), astrocytes (S100β, green), and neurons (MAP2, white). Nuclei were stained with DAPI (blue). Scale bars, 50 µm. **b,** Predicted senescence scores of untreated and CDDO-Me-treated human iPSC-derived microglia, neurons, and astrocytes in tricultures. Mean ± SEM; unpaired two-tailed *t* test compared with untreated controls in each cell type, \**P* = 0.0106 (microglia), 0.0138 (neurons); ns = not significant; n = 3 independent experiments. **c-d,** Cell-cell communication in tricultures analyzed using CellChat. **c,** Circle plots showing number of putative ligand–receptor (L-R) interactions in control versus CDDO-Me-treated cells. Edge width represents communication strength. **d,** Bubble plot depicting up- and downregulated L-R pairs in microglia within the triculture. Colors indicate communication probability, comparing CDDO-Me treatment with untreated control. Black font: interactions under control; red font: interactions under CDDO-Me. Significance determined via permutation test (n = 100); bubbles represent significant L-R interactions (nominal *P* < 0.01). **e,** Bubble plot showing downregulated Reactome pathways in the microglial cluster. Bubble diameter indicates gene count; color represents -log10(FDR). **f,** Fluorescence imaging of Alexa Fluor 488-labeled beads (green) and IBA1 (red) in untreated and CDDO-Me-treated microglia. Nuclei stained with DAPI (blue). BF: bright field. Scale bars, 10 µm. Images are representative of at least two independent experiments. **g,** Immunostaining for P62 (green) and IBA1 (red) in untreated and CDDO-Me-treated tricultures. Nuclei: DAPI (blue). BF: bright field. Right panel: quantification of P62 intensity in IBA1⁺ and IBA1⁻ areas. Data normalized to untreated controls. Center line, median; box, 25th–75th percentiles; whiskers, min–max; unpaired two-tailed *t* test, \**P* = 0.0167; n = 4 independent experiments. **h,** Western blot of LC3-II in microglia, untreated or treated with CDDO-Me, with or without bafilomycin (Baf, 4 hours). Autophagic flux quantified as the ratio of LC3-II in Baf-treated versus untreated cells, normalized to untreated control. Mean ± SEM; unpaired two-tailed *t* test, \**P* = 0.0192; n = 3 independent experiments. **i,** Confocal images of microglia untreated or treated with CDDO-Me, stained for CD68 (green) and α-synuclein preformed fibrils (PFF-594, red) at 60- and 90-min post-treatment. Nuclei: DAPI (blue). Scale bars, 10 µm. Right panel: quantification of PFF-594 area relative to total cell area. Center line, median; box, 25th–75th percentiles; whiskers, min–max; two-way ANOVA with Bonferroni *post hoc* correction, \**P* = 0.0170, \*\**P* = 0.0058 (untreated 60 min versus untreated 90 min), 0.0048 (untreated 90 min versus CDDO-Me 90 min); n = 4 independent experiments.

### UPR^mt^ activation induces cell type-specific metabolic dysregulation in iPSC-derived tricultures

To investigate intercellular mechanisms of UPR^mt^ activation, we performed single-cell RNA-sequencing in CDDO-Me-treated tricultures. UMAP integration and clustering identified 11 clusters, including two microglial, one immature neuronal, three neuronal, four astrocytic, and one oligodendrocyte progenitor cluster (OPC) (Supplementary Figure 13B-E; Supplementary Data 4). CDDO-Me did not alter cluster proportions (Supplementary Figure 13F).

UCell analysis revealed lower basal UPR^mt^ activity in neurons than glia, with CDDO-Me increasing scores across all cell types (Supplementary Figure 13G, H). Comparing monoculture and triculture DEGs identified 985 triculture-specific dysregulated genes (63 neuronal, 308 astrocytic, 614 microglial; Supplementary Figure 13I; Supplementary Data 5), highlighting context-dependent transcriptional responses. Reactome analysis showed disruption of inflammatory pathways, including lipid/atherosclerosis and antiviral responses (Supplementary Data 6). UPR^mt^ activation induced cell type-specific metabolic rewiring: microglia upregulated nucleotide sugar metabolism (R-HSA-00520, -01250), whereas neurons upregulated amino acid and one-carbon metabolism, as well as cysteine/methionine pathways (R-HSA-01230, -00260, -00220, -00250, -01240, -01200, -00670, -00270). These results indicate that glia-neuron crosstalk shapes inflammatory and metabolic responses to mitochondrial proteotoxic stress.

### Mitochondrial proteotoxic stress decreases predicted microglia-mediated cell-cell communication and phagocytic signaling in iPSC-derived tricultures

To define how UPR^mt^-driven senescence reshapes intercellular communication, we analyzed ligand-receptor interactions in the single-cell dataset using CellChat^56^. CDDO-Me treatment globally reduced both the number and strength of predicted interactions in tricultures, indicating an overall attenuation of cell-cell communication (Figure 5C). Cell type-resolved analysis revealed increased signaling among astrocytes and neuronal populations, but a marked reduction in autocrine and paracrine communication from microglia and OPCs (Supplementary Figure 14A).

In microglia, CDDO-Me enhanced ANXA1-FPR1/3 and TNF-TNFRSF1A/1B signaling, consistent with increased inflammatory crosstalk toward astrocytes, OPCs, and neighboring microglia (Figure 5D). Conversely, microglial SPP1-(ITGAV+ITGB1) signaling to neurons, astrocytes, OPCs, and microglia was reduced, as was SPP1-CD44 signaling to microglia and OPCs and GRN–SORT1 signaling to astrocytes, OPCs, and neurons (Figure 5D). Given the role of SPP1 in macrophage phagocytosis and microglial synapse engulfment ^57^, these changes suggest impaired clearance-related communication.

Consistently, Reactome analysis of microglial DEGs revealed enrichment of phagocytosis-related pathways (FcγR-mediated phagocytosis, lysosome, phagosome), infection-response pathways, and lipid and atherosclerosis pathways (Figure 5E; Supplementary Figure 14B). Functional assays in tricultures showed unchanged latex bead uptake but significant accumulation of p62 in CDDO-Me-treated microglia, indicating defective autophagic processing (Figure 5F,G). A similar phenotype was observed in tricultures containing *PITRM1*-KO microglia with wild-type neurons and astrocytes (Supplementary Figure 14C, D), demonstrating that microglia-intrinsic mitochondrial stress is sufficient to impair clearance mechanisms. To determine whether synaptic remodeling was affected, we assessed PSD95 accumulation. CDDO-Me treatment and *PITRM1* deficiency both increased PSD95 signal within microglia and across cultures, without changes in C1QA levels (Supplementary Figure 14E), consistent with impaired synaptic pruning ^58^. In monoculture, bead uptake was reduced after CDDO-Me treatment (Supplementary Figure 14F). LAMP1 intensity was significantly increased in IBA1-positive microglia, suggesting lysosomal stress or expansion (Supplementary Figure 14G). Moreover, bafilomycin A1 further reduced autophagic flux in CDDO-Me-treated cells (Figure 5H; Supplementary Figure 14H). Together, these findings demonstrate that mitochondrial proteotoxic stress induces a microglial senescence-like state characterized by impaired intercellular communication, reduced phagocytic signaling, and autophagolysosomal dysfunction, features highly relevant to neurodegenerative disease contexts.

### UPR^mt^ -induced misfolded protein accumulation in senescent microglia is reversed by inhibition of polyamine synthesis

To examine the relevance of UPR^mt^-driven senescence to protein aggregation, we cotreated control iPSC-derived microglia with CDDO-Me and Alexa Fluor 594-labeled α-synuclein preformed fibrils (PFFs). Immunostaining at 60 and 90 min revealed increased intracellular α-synuclein accumulation in CDDO-Me-treated microglia compared to controls, indicating impaired fibril clearance (Figure 5I). *PITRM1*-KO microglia similarly exhibited elevated intracellular α-synuclein relative to isogenic controls (Supplementary Figure 14I).

To test whether polyamine synthesis contributes to this phenotype, we cotreated microglia with α-synuclein PFFs and CDDO-Me in the presence or absence of the polyamine synthesis inhibitor MCHA. Live-cell imaging confirmed progressive α-synuclein accumulation in CDDO-Me-treated cells (Supplementary Figure 14J). Notably, MCHA significantly reduced intracellular α-synuclein levels compared to CDDO-Me alone, and this effect persisted over time (Supplementary Figure 14J). These findings demonstrate that UPR^mt^-induced misfolded protein accumulation in senescent microglia is dependent on SAM-driven polyamine metabolism and can be reversed by its inhibition.

### Microglial UPR^mt^ drives senescence and protein aggregation in MgBr assembloids

To investigate the impact of microglial mitochondrial stress on brain senescence, we generated MgBr assembloids by integrating WT or *PITRM1*-KO microglia into WT cortical organoids at day 30 (Supplementary Figure 15A, B). Microglia successfully integrated and persisted at 5, 15, and 35 days post-integration (DPI) (Supplementary Figure 15C-E). Expression of endodermal, neuronal progenitor, and mesodermal markers was comparable across conditions at 35 DPI (Supplementary Figure 16A, B), indicating preserved lineage composition. In contrast, senescence markers revealed a progressive increase in p21 expression in organoids containing *PITRM1*-KO microglia, without changes in DNA damage markers (Figure 6A-C; Supplementary Figure 16C, D). Notably, MAP2 expression was significantly reduced at 35 DPI (Figure 6D), suggesting impaired neuronal integrity associated with microglial mitochondrial stress. Assessment of amyloidogenic protein accumulation using thioflavin T demonstrated significantly increased fluorescence in *PITRM1*-KO MgBr assembloids at 35 DPI (Figure 6E). Consistently, pathological Aβ species were elevated (Figure 6F), whereas phosphorylated tau levels remained unchanged (Supplementary Figure 16E, F), consistent with an early-stage pathology in which Aβ deposition precedes tau alterations ^59–61^. Together, these findings indicate that microglial UPR^mt^ promotes a senescence-associated phenotype and accelerates amyloid accumulation in MgBr assembloids.

**Figure 6.**
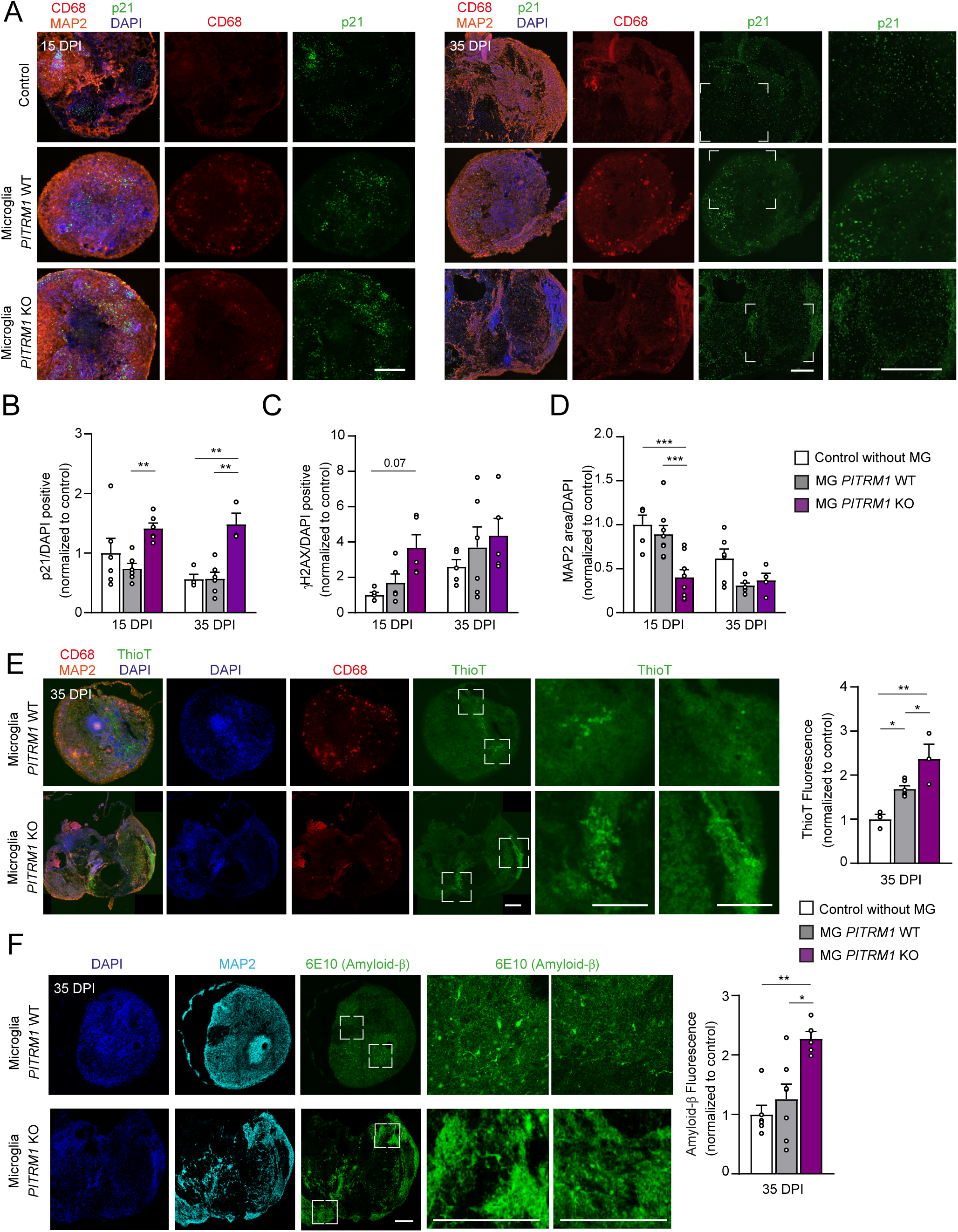
Mitochondrial proteotoxic stress in microglia induces senescence propagation and misfolded protein accumulation in MgBr assembloids. All experiments were performed using iPSC line C2 and *PITRM1* KO2. **a,** Immunostaining for p21 (green), CD68 (red), and MAP2 (orange) in MgBr assembloids at 15 and 35 days post integration (DPI). Nuclei were counterstained with DAPI (blue). High-magnification images of the sections are shown on the right. Scale bars, 25 µm. **b–d,** Quantification of p21⁺/DAPI⁺ cells (**b**), γH2AX⁺/DAPI⁺ cells (**c**), and MAP2/DAPI area (**d**) in whole sections at 15 and 35 DPI. Data are shown as mean ± SEM and normalized to control organoids without microglia integration. Two-way ANOVA with Bonferroni *post hoc* correction between genotypes at each time point. Significant comparisons: (b) \*\**P* = 0.0069 (15 DPI, *PITRM1*-WT vs *PITRM1*-KO), 0.0054 (35 DPI, Control vs *PITRM1*-KO), 0.0029 (35 DPI, *PITRM1*-WT vs *PITRM1*-KO); (d) \*\*\**P*=0.0002 (15 DPI, Control vs *PITRM1*-KO), 0.0004 (15 DPI, *PITRM1*-WT vs *PITRM1*-KO). Each n represents one individual organoid. For panel b, n=6/4 Control, 6/6 WT, and 6/3 KO at 15/35 DPI, respectively. For panel c, n=5/5 Control, 5/6 WT, and 5/5 KO. For panel d, n=5/6 Control, 8/7 WT, and 8/4 KO. **e,** Left: Immunostaining for CD68 (red) and MAP2 (orange) in 35 DPI MgBr assembloids. Nuclei stained with DAPI (blue); protein aggregates detected with Thioflavin T (ThioT, green). High-magnification images are shown on the right. Scale bars, 25 µm. Right: Quantification of ThioT fluorescence in whole sections. Mean ± SEM, normalized to control organoids without microglia. One-way ANOVA with Bonferroni *post hoc* correction; \**P*=0.0402 (Control vs *PITRM1*-WT), 0.0409 (*PITRM1*-WT vs *PITRM1*-KO), \*\**P*=0.0015; n=3 Control, n=6 WT, n=3 KO individual organoids. **f,** Left: Immunostaining for amyloid-β (6E10, green) and MAP2 (cyan) in 35 DPI MgBr assembloids. Nuclei stained with DAPI (blue). High-magnification images are shown on the right. Scale bars, 20 µm. Right: Quantification of amyloid-β fluorescence in whole sections. Mean□±□SEM; normalized to control organoids without microglia integration. One-way ANOVA with Bonferroni *post hoc* correction; \**P* = 0.0136, \*\**P* = 0.0053. n: n=5 Control, n=7 WT, n=5 KO individual organoids.

## Discussion

We define cell type-specific roles for mitochondrial stress responses in the human brain and identify the UPR^mt^ as a key driver of microglial senescence. UPR^mt^ activation in human iPSC-derived microglia induces coordinated dysregulation of SAM metabolism and TAG-centered lipid remodeling, leading to impaired autophagolysosomal and phagocytic function, altered intercellular communication, and compromised neuronal integrity. These findings link mitochondrial proteostasis defects to glia-driven brain aging and neurodegeneration.

Consistent with previous studies in mammalian and invertebrate systems, mitochondrial proteotoxic stress activated the UPR^mt^ in neurons, astrocytes, and microglia, accompanied by reduced mtDNA content, decreased respiratory complex levels, and increased chaperone expression ^35, 62^. However, glial cells, particularly microglia, exhibited earlier and more robust stress responses than neurons, suggesting an evolutionary specialization in which glia act as primary sensors, activating stress programs before neurons reach their own threshold ^14, 63^. Single-cell analyses in tricultures further revealed that astrocytes and microglia contribute disproportionately to mitochondrial stress signaling, suggesting glial specialization in stress sensing and response. Despite UPR^mt^ activation, microglia displayed a maladaptive program: downregulation of autophagy and proteasome genes combined with upregulation of translational machinery, reminiscent of ISR activation ^24^. In contrast, neurons and astrocytes mounted compensatory proteostatic programs. Convergence of UPR^mt^ and ISR signaling in microglia may therefore promote a neurotoxic state.

Because defining senescence in postmitotic brain cells is challenging ^28, 64,30, 65, 66^, we combined transcriptional profiling, molecular analyses, and machine learning-based nuclear morphology assessment to define a UPR^mt^-driven microglial senescence phenotype. This state is characterized by nuclear alterations, ISR-enriched transcriptional signatures, lipid droplet accumulation, and defective phagocytosis-features absent in neurons or astrocytes under identical stress conditions, underscoring the selective vulnerability of microglia.

Previous research in simple organisms indicate that mitochondrial stress enhances neuronal and astrocyte signaling and thereby contributes to brain homeostasis ^14, 17^. However, our findings suggests a more divergent response in humans. While glial cells in *C. elegans* and humans share fundamental roles, human glial cells exhibit greater complexity and specialization in neural function, development, and regeneration ^67^. Functionally, UPR^mt^ activation enhanced neuron-astrocyte communication but reduced microglial signaling, including decreased SPP1- and GRN-mediated phagocytic pathways. This was accompanied by increased TNF-dependent inflammatory signaling and impaired clearance of misfolded proteins. In MgBr assembloids, integration of *PITRM1*-KO microglia led to progressive p21 induction, reduced MAP2 expression, and increased amyloid accumulation, supporting a causal contribution of microglial mitochondrial stress to brain senescence and early neurodegenerative pathology, potentially by driving the SASP ^68^.

Lipid remodeling emerged as a central mechanism. Mitochondrial stress induced profound alterations in TAG metabolism, glycerophospholipid remodeling, oxidative stress, and depletion of methionine-derived metabolites including SAM. We show that UPR^mt^ activation triggers a transition toward a dysfunctional state reminiscent of LD-accumulating microglia and senescent microglia, observed near tau and amyloid pathology in the aging brain ^69, 70^. Importantly, this lipid accumulation is not merely a byproduct but a driver of dysfunction, as DGAT1/2 inhibition rescued both lipid storage and senescence markers. Interestingly, despite divergent TAG levels in pharmacological versus genetic models, both converged on a senescent phenotype, suggesting that lipid flux rather than absolute lipid content is a critical determinant of cellular health ^71,72, 73, 74^. This aligns with lipid dysregulation as a conserved hallmark of senescence across species ^75, 53^.

We further identify SAM metabolism as a key regulator of this process. Limiting SAM consumption or synthesis attenuated lipid accumulation and senescence markers, consistent with prior evidence that elevated SAM can promote UPR^mt^ activation ^76^. This metabolic dependency helps explain distinct stages of the stress response: while targeting mitochondrial ROS with MitoQ prevented senescence during pharmacological stress, it failed to rescue the defects driven by chronic genetic UPR^mt^ activation. Together, these findings support a two-phase model of microglial decline: an initial ROS-dependent activation phase followed by a chronic, self-sustaining phase driven by the coordinated dysregulation of SAM and lipid metabolism. Ultimately, these data position one-carbon metabolism as the central node coupling mitochondrial stress to lipid remodeling and the progression toward senescence.

This evidence supports a convergent mechanism in which mitochondrial stress and ISR signaling cooperate to drive a lipid-mediated neurotoxic microglial state. Our findings extend the concept of ISR-marked neurodegenerative microglial subsets ^24^, which in AD models promote “dark microglia” and exacerbate synapse loss and proteinopathy. We show that targeting specific branches of lipid metabolism may attenuate microglia-driven senescence and neurodegeneration. Notably, the mitochondrial proteostasis stress signature identified in iPSC-derived microglia overlaps with disease-associated clusters in the aging and AD brain, including ISR activation, early stress responses, and impaired phagocytic signaling, indicating that these maladaptive programs are active in human neurodegeneration.

Together, our findings highlight the context-dependent and cell type-specific nature of UPR^mt^ activation. While mitochondrial stress responses can support neuronal proteostasis ^8, 9, 14^, sustained microglial UPR^mt^ activation drives a lipid-mediated, ISR-associated senescent state that disrupts neuronal homeostasis. These results argue against indiscriminate modulation of mitochondrial stress pathways and instead support microglia-targeted strategies, potentially via lipid nanoparticle delivery or selective modulation of downstream metabolic branches, to mitigate aging- and neurodegeneration-associated pathology.

## Acknowledgments

This research was funded by Aligning Science Across Parkinson’s (Grant number: ASAP-000420) through the Michael J. Fox Foundation for Parkinson’s Research (MJFF) (to M.D.), ERC CoG (#101003329, to M.D.). M.D. received funding from DFG-ANR (#490761034), the MetaboBrain Index excellence program, and the EU Joint Program-Neurodegenerative Disease Research (JPND) project (GBA-PaCTS, 01ED2005B). We acknowledge the funding support of the Innovative Minds Program-Foundation Deutsche Demenzhilfe, EMBO Long term postdoctoral fellowship ALTF 1013-2019, Becas Chile-ANID 74190073 (to M.J.P.). The M.D. and M.S.K. laboratories are supported by funding from the European Union’s Horizon Europe Research and Innovation Program through the MSCA-Doctoral Network NADIS (no. 101073251). Work in *Pitrm1* mice models was funded by Fondazione Telethon and AISA (Project ID GSA22I001 to DB). We acknowledge Région Île-de-France for funding support toward the acquisition of the spectral flow cytometer SONY ID7000, which is used at the Cytometry Platform of the SFR Necker. We thank Dr. Daniel Ackerman of Insight Editing London for editing the manuscript prior to submission. The authors thank Dr. Graziella Cappelletti (The University of Milan, IT) for kindly providing the α-synuclein monomers.

## Author Contribution Statement

M.J.P. and M.D. conceived and designed the experiments; M.J.P., M.B., F.B., H.R., A.L., and S.K. contributed to the cell culture work; M.J.P., M.B., F.B., M.K., C.W., and A.L. contributed to the validation of the transcriptomic analysis and data analysis; C.W. performed the cell cycle analysis; I.N. performed the metabolomic analysis; Insa H., A.L., and C.W. performed the cytokine measurements; Indra H. and M.S.K. performed the senescence measurements and analysis; D.B., A.L., and M.J.P. performed the mouse experiments; H.R., M.J.P., and A.L. performed and analyzed the organoid experiments; M.J.P. and M.D. wrote the manuscript with contributions from all of the authors. All of the authors read and approved the final manuscript.

## Methods

### Human iPSCs

Three healthy control lines were used in the study: C1 and C2^8, 77, 78^, and the commercial line C3 (Kolf2.1J, RRID: CVCL_B5P3; Jackson Laboratory). C1 and C2 iPSC lines were derived with informed consent and approved by the Ethics Committee of the Medical Faculty and University Hospital Tübingen (Ethikkommission der Medizinischen Fakultät am Universitätsklinikum Tübingen, 199/2011BO1). *PITRM1*-knockout lines were generated from C2^8^. All lines were routinely tested for mycoplasma (Venor®GeM Classic, Minerva Biolabs Inc., Cat. #11-1250) and maintained in mTeSR™ Plus (STEMCELL Technologies; Cat. # 100-0276).

### Human iPSC differentiation

Human iPSCs were differentiated into cortical neurons, astrocytes, and microglia using established stepwise protocols (see Supplementary Methods). Cortical neurons were generated via neural progenitors^8, 79^ and used after 21 days in vitro. Astrocytes were derived from neural progenitors^80^ and used between passages 3-10. Microglia were produced through embryoid body-based hematopoietic specification ^78, 81^ followed by 10 days of terminal differentiation. Cells were maintained as monocultures or combined to generate multicellular systems. Tricultures were established by sequentially adding astrocytes and microglia to neurons at a defined ratio (neurons:astrocytes:microglia = 3:1:1) and maintained in maturation medium supplemented with microglial survival factors prior to experiments.

### Microglia-containing brain (MgBr) assembloids

Cortical organoids were generated ^82^ and later integrated with pre-differentiated iPSC-derived microglia to produce microglia-containing brain (MgBr) assembloids. Assembloids were maintained under orbital shaking and analyzed at defined days post-integration. Detailed differentiation conditions, media compositions, and assembly procedures are described in the Supplementary Methods.

### Drug treatments

Human iPSC-derived neurons, astrocytes, and microglia were exposed to increasing concentrations of CDDO-Me (CAS number 218600-53-4, Cayman Chemical, Cat. # 11883) for the durations indicated in the figure legends. iPSC-derived microglia were exposed to GTTP (CAS number 1131626-47-5; MedChemExpress, Cat. # 50-196-7948). Additional treatments included ultrapure lipopolysaccharides from Escherichia coli O111:B4 (LPS-EB Ultrapure; InvivoGen, Cat. # tlrl-eblps), SAM-HCl (CAS No. 86867-01-8; Sigma-Aldrich, Cat. # CIAH9ABED0C7), FIDAS V (CAS No. 1391934-98-7; MedChemExpress, Cat. # HY-136144), MCHA (Sigma-Aldrich, Cat. # CDS019932), MitoQ (CAS No. 845959-50-4; MedChemExpress, Cat. # HY-100116AR), and etoposide (CAS No. 33419-42-0; MedChemExpress, Cat. # HY-13629). For TAG inhibition, cells were treated with the DGAT1 inhibitor PF-04620110 (CAS No. 1109276-89-2; MedChemExpress, Cat. # HY-13009) and the DGAT2 inhibitor PF-06424439 (CAS No. 1469284-78-3; MedChemExpress, Cat. # HY-108341).

### LONP1 knockdown in iPSC-derived microglia

iPSC-derived microglia were transduced with lentiviral particles encoding an shRNA targeting human *LONP1* (shLONP1; TRCN0000046793, MISSION® shRNA library, Sigma-Aldrich) or a non-targeting control shRNA (VectorBuilder, Cat. # VB010000-0005mme). The shLONP1 construct (target sequence: CCAGTGTTTGAAGAAGACCAA) was cloned into the pLKO.1-puro backbone. Lentiviral particles were generated in HEK293T cells (RRID: CVCL_0063) by co-transfection of the shRNA plasmid with psPAX2 (RRID: Addgene_12260) and pMD2.G (RRID: Addgene_12259) using TransIT-X2 (Mirus Bio, Cat. # MIR 6003) according to the manufacturer’s protocol. Viral titers were normalized based on p24 capsid levels quantified with Lenti-X GoStix Plus (Takara Bio, Cat. # 631280), and equal amounts of virus were applied directly to microglial cultures without additional transfection reagents. Cells were harvested for analysis 24, 48, and 72 hours post-transduction.

### ATF5-inducible human iPSCs

iPSCs were transduced with lentiviral vectors encoding doxycycline-inducible constructs for *ATF5* overexpression (VectorBuilder, ID: VB241111-1111fuf) or an empty vector control (VectorBuilder, ID: VB201122-1083bzq), both containing a neomycin resistance cassette. Clonal selection was performed with G418 (150 µg/mL; Thermo Scientific, Cat. # 10131035) for two weeks. Stable clones were expanded and used for experiments without further antibiotic selection.

### Mouse studies

All experiments were conducted in accordance with European Directive 2010/63/EU and Italian law (D. Lgs. n. 26/2014) and approved by the Italian Ministry of Health (authorization 483/2023-PR). C57BL/6N-Atm1Brd *Pitrm1*+/tm1a(KOMP)Wtsi mice (allele ID: MGI:5085349), kindly provided by the Wellcome Trust Sanger Institute, were derived from the KOMP-CSD ES cell line (RRID: MMRRC_060296-UCD). Animals were housed in groups of three per cage (22 ± 2°C, 12 h light–dark cycle, 60% relative humidity) with *ad libitum* access to food (rodent diet 4RF25, Mucedola, Italy) and water. Two groups of female mice (6 and 15 months; 6 *Pitrm1*+/+ and 6 *Pitrm1*+/-) were used. Brains were either formalin-fixed for histology or half-brains were fresh-frozen for Western blot or embedded in OCT (Tissue-Tek, Sakura Finetek, Cat. # 4565). Cryosections (20 µm) were stored in antifreeze solution.

### Immunofluorescence and live-cell imaging

Cultures and tissue sections were fixed with paraformaldehyde (PFA, Sigma-Aldrich, Cat. # 1004968350), permeabilized, and blocked before incubation with primary and fluorescent secondary antibodies. Nuclei were counterstained with DAPI (Sigma-Aldrich, Cat. # D9542-10MG). Lipid droplets and protein aggregates were visualized using BODIPY (Thermo Scientific, Cat. # D3922) and thioflavin-T (Sigma□Aldrich, Cat. # T3516), respectively, where indicated. Images were acquired by Leica TCS SP8 confocal microscope (60×/1.4□NA oil objective), and quantitative analyses, including nuclear marker quantification, fluorescence intensity measurements, and colocalization, were performed using CellProfiler (Version 4.2.6, RRID: SCR_007358) and Fiji (Version 2.7.0, RRID:SCR_002285).

Cells were incubated at 37 °C with fluorescent probes to assess mitochondrial membrane potential (TMRM and MitoTracker Green; Invitrogen, Cat. # T668 and # M46750), intracellular calcium levels (Fluo-4 AM; Invitrogen, Cat. # F14217), or oxidative stress (DCF; Invitrogen, Cat. # C400). After washing, cells were imaged by Leica TCS SP8 confocal microscope (60×/1.4□NA oil objective). For calcium imaging, neurons and astrocytes were stimulated with KCl (Sigma-Aldrich, Cat. #P3911-25G) and glutamate (Sigma-Aldrich, Cat. # 49621-250G), respectively. Fluorescence signals were quantified using Fiji following background subtraction and normalization to baseline. Detailed acquisition settings and analysis workflows are provided in the Supplementary Methods.

### Phagocytosis assay

Microglia or tricultures were incubated with Alexa Fluor 488-labeled latex beads (Sigma□Aldrich, Cat. # L1030) for 3 hours at 37□°C; beads were pre-opsonized in fetal bovine serum (FBS, Thermo Scientific, Cat. # A5256801) for 1 hour at 37°C. Cells were washed with PBS (Thermo Scientific, Cat. # 14190169) and fixed with 4% PFA for immunostaining. For flow cytometry, samples were dissociated with Accutase (STEMCELL Technologies, Cat. # 07920), incubated with Zombie NIR viability dye (1:1000, Biolegend, Cat. # 423105) in PBS, washed in PBS with 2% FBS, and analyzed on an Acea NovoCyte® flow cytometer (Agilent Technologies) using FlowJo™ software (BD Biosciences; RRID:SCR_008520).

### Cytokine measurements

Cytokine levels in human iPSC-derived microglia were quantified via LEGENDplex^TM^ bead-based immunoassays (BioLegend GmbH). IL-1β, IL-4, IL-6, TNF-α, IFN-γ, MCP-1, and IL-8 in cell supernatants were measured with the Human Essential Immune Response Panel (BioLegend GmbH, Cat. # 740930) and a Sony ID7000^TM^ spectral cell analyzer (Sony Biotechnology, USA). Data (>300 beads per sample) were analyzed with LEGENDplex™ software v9.0 (BioLegend, https://www.biolegend.com/en-us/legendplex).

Cytokine mouse brains were measured using Mouse TNF-α DuoSet ELISA (R&D Systems, Cat. # DY410) and Mouse IL-1β/Il-1F2 DuoSet ELISA (R&D Systems, Cat. # DY201). Whole brains from 6- and 15-month-old mice were rinsed with PBS and homogenized in ice-cold buffer (50 mM Tris-HCl, pH 7.4, 150 mM NaCl, 1% Triton X-100, 1 mM EDTA, and protease/phosphatase inhibitors (Pierce, Thermo Scientific, Cat. # A32959) using 2 mm stainless steel beads. The lysates were centrifuged at 14,000□× g for 20 minutes, and the supernatants were collected for analysis. The absorbance was measured at 450□nm with wavelength correction at 540□nm using a PerkinElmer Envision 2105 plate reader. Cytokine concentrations were normalized to the total protein content determined by BCA assay (Pierce, Thermo Scientific, Cat. # 23225).

### Quantitative Real-Time PCR

mRNA was extracted from iPSC-derived neurons, astrocytes, and microglia using the RNeasy Mini Kit (QIAGEN, Cat. # 74106). cDNA was synthesized with the QuantiTect Reverse Transcription Kit (QIAGEN, Cat. # 205313). Quantitative PCR was performed with SYBR Green PCR Master Mix (QIAGEN, Cat. # 204145) on a Viia 7 Real-Time PCR System (Applied Biosystems). Gene expression was normalized to the housekeeping genes *RPLP0* or *ACTB*, and relative expression levels were calculated using the 2^−ΔΔCT method. Primer sequences are provided in Supplementary Table 2.

### Mitochondrial isolation

Mitochondrial isolation was performed with the QIAGEN Qproteome^®^ Mitochondria Isolation Kit (Qiagen, Cat. # 37612) with modifications. Pelleted cells (10 × 10^6) were incubated on ice for 10 minutes in 1 mL of lysis buffer with protease inhibitor. The suspension was centrifuged at 1000 × g for 10 minutes at 4°C. Half of the supernatant was retained as the total cellular fraction, and the remainder was subjected to three additional centrifugation rounds (1000 × g, 10 minutes, 4°C) to obtain the cytosolic fraction. The pellet was resuspended in 1.5 mL of disruption buffer with protease inhibitors and mechanically disrupted via 26-gauge needle ten times on ice. After a first centrifugation (1,000 × g, 10□min, 4□°C), the pellet was discarded, and the supernatant was further centrifuged at 14,000 × g for 30□min at 4□°C. The final mitochondrial pellet was resuspended in 50 μL of storage buffer with protease inhibitors. Protein concentration was measured with the Pierce™ BCA Protein Assay Kit; 20□µg was used for immunoblotting and 30□µg for protein aggregation assays.

### Western blot

Protein lysates from cells and mouse brain tissue were prepared using detergent-based extraction buffers containing protease and phosphatase inhibitors. Protein concentrations were determined by BCA assay, and equal amounts of protein were resolved by SDS–PAGE (Surepage, GenScript, Cat. # M00654) and transferred to PVDF membranes (Thermo Scientific, Cat. # 10617354). Membranes were blocked and incubated with primary antibodies overnight, followed by HRP-conjugated secondary antibodies. Proteins were visualized using chemiluminescence and imaged with digital detection systems. Band intensities were quantified using ImageJ and normalized to loading controls. Antibodies are listed in Supplementary Table 1, and detailed buffer compositions and electrophoresis conditions are provided in Supplementary Methods.

### Protein aggregation measurement

Protein aggregation was assessed using the PROTEOSTAT® Protein Aggregation Assay Kit (Enzo Life Sciences, Cat. # ENZ-51023-KP050) with modifications. The detection dye was diluted 1:2 in 1× assay buffer, and 2□µL was added per well of a black 96-well plate with clear bottom. Thirty micrograms of protein per sample were diluted to 100□µL in assay buffer, incubated 15□min at room temperature in the dark, and fluorescence was measured (550□nm excitation/600□nm emission) using a PerkinElmer EnVision 2105 plate reader. For imaging-based detection, the PROTEOSTAT® Aggresome Detection Kit (Enzo Life Sciences, Cat. # ENZ-51035-KIT) was used. Fixed cells were permeabilized with 0.1% Triton X-100 in PBS, incubated with the detection reagent for 30□min in the dark, counterstained with DAPI, and imaged on a Leica TCS SP8 confocal microscope (60×/1.4□NA oil objective).

### Bulk RNA-sequencing and data analysis

Total RNA from iPSC-derived neurons, astrocytes, and microglia (three biological replicates per condition) treated with or without CDDO-Me was subjected to poly(A)-selected, strand-specific bulk RNA sequencing (paired-end, 150 bp; Illumina NovaSeq). Differential gene expression analysis was performed using DESeq2 (version 1.44.0, RRID: SCR_015687), applying a fold-change cutoff >2 and Benjamini–Hochberg FDR < 0.05. Functional enrichment analysis of differentially expressed genes was conducted using gene ontology and pathway analysis tools, as described in Supplementary Methods. Overlap between cell type-specific gene sets and published human microglial signatures was assessed using Fisher’s exact test. Gene set enrichment analysis (GSEA, version v4.3.2, https://www.gsea-msigdb.org; RRID: SCR_003199) was performed using the Molecular Signatures database (MSigDB, version v6.2, RRID: SCR_016863). To evaluate enrichment of senescence-, SASP-, and cell cycle-related pathways^7, 25–28, 83^, genes were ranked based on differential expression between treated and control samples, and significance was determined using normalized enrichment scores and FDR correction. Full parameters and gene set details are provided in Supplementary Methods.

### Cell cycle analysis by flow cytometry

iPSC-derived cells were dissociated using Accutase, resuspended in PBS, and stained with Hoechst 33342 (2.5 µg/mL; Sigma-Aldrich, Cat. # 14533) for 30 minutes at 37°C. After washing, cells were incubated with Zombie NIR viability dye (1:1000 dilution in PBS). The cells were then washed and resuspended in PBS containing 2% FBS. Flow cytometry was performed on a Sony ID7000™ spectral cell analyzer with a maximum event rate of 200 events per second. The data were analyzed in FlowJo™ software using the Watson (Pragmatic) model. The G1, S, and G2/M phases were distinguished based on the Hoechst signal intensity. Model fitting was visualized by overlaying the model sum and the actual signal distribution. Peak 1 was set to “n” and Peak 2 was constrained to match the coefficient of variation of the G1 peak. The final model selection was based on achieving a low root mean square deviation value.

### Senescence β-Galactosidase staining

Senescence-associated β-galactosidase staining was performed using the Senescence β-Galactosidase Staining Kit (Cell Signaling Technology, Cat. # 9860). Cells were cultured on Matrigel-coated coverslips and fixed with the provided fixation solution. For mouse tissue, OCT-embedded sections were cryosectioned, mounted on Epredia™ SuperFrost™ Plus slides (Thermo Scientific, Cat. # 10149870), refixed, and stained following the manufacturer’s instructions. Images were acquired using a Leica DM750 brightfield microscope (40×/0.4□NA) for cells or an EVOS M7000 Imaging System (20×/0.4□NA; Thermo Scientific) for tissue sections.

### NAD consumption assay

NAD consumption was assessed with the fluorescent NAD analog nicotinamide 1,N^6^-ethenoadenine dinucleotide (Sigma-Aldrich, Cat. # N2630). iPSC-derived microglia (1 × 10^5 per condition) were detached with Accutase and resuspended in 100 μL PBS. Eighty micromolar of the analog (340□nm excitation/460□nm emission) was added, and fluorescence was recorded every minute for 1□h at 37□°C using a SpectraMax M2 reader (Molecular Devices). NADase activity was calculated from the linear slope and normalized to total protein content. Protein concentrations were determined with a Pierce™ BCA protein assay kit.

### Nuclear senescence scoring

iPSC-derived monocultures and tricultures were stained with DAPI and immunolabeled for MAP2, IBA1, and GFAP. Images were acquired using a Leica TCS SP8 confocal microscope equipped with a 40×/1.4 NA oil immersion objective. Quantitative senescence scoring was performed as previously described ^51^. Senescence prediction was based on nuclear morphology. Nuclei were segmented from DAPI images using a U-Net architecture, and morphometric features (area, convexity, and aspect ratio) were extracted. Segmented nuclei were converted into two-channel masks and analyzed using an ensemble of deep neural networks to generate senescence probability scores. The models were originally trained on fibroblasts and validated across multiple cell types ^51^. For triculture experiments, images were split into individual channels (IBA1, S100β, and MAP2). A minimum integrated intensity threshold was applied to each channel to identify marker-positive cells. Objects exceeding the threshold were classified according to the channel with the highest integrated density signal. Analyses were performed using Python 3.9.18 (RRID:SCR_008394), TensorFlow GPU 2.15.0 (RRID:SCR_016345), Keras 2.15.0 (RRID:SCR_026159), NumPy 1.23.5 (RRID:SCR_008633), and Pandas 1.2.4 (RRID:SCR_018214).

### Lipidomic and metabolomic profiling of iPSC-derived microglia

Lipidomic analysis was conducted by Lipotype GmbH (Dresden, Germany) using high-resolution shotgun mass spectrometry on an Orbitrap platform with class-specific internal standards. Lipid species were identified based on MS and MS/MS fragmentation patterns and quantified relative to internal standards. For metabolomics, polar metabolites were extracted using methanol/acetonitrile-based solvents, separated by hydrophilic interaction liquid chromatography, and analyzed by high-resolution Orbitrap mass spectrometry operated in full-scan polarity-switching mode. Metabolites were annotated based on accurate mass and retention time matching and quantified from peak areas. Statistical and pathway enrichment analyses were performed using established metabolomics workflows. Detailed extraction procedures, chromatographic gradients, mass spectrometry acquisition parameters, internal standards, and data processing pipelines are provided in Supplementary Methods.

### Single-cell RNA sequencing and analysis

Single-cell suspensions from iPSC-derived tricultures were processed using the 10x Genomics Chromium Next GEM Single Cell 3′ v3.1 kit (10x Genomics, Cat. # 1000128). Libraries were sequenced on an Illumina NovaSeq 6000 (SP Flow Cell, Illumina) paired-end to a depth of ∼300 million reads per library (GENEWIZ GmbH). After Cell Ranger processing, a total of 7,160 high-quality cells were retained for analysis. Analyses were performed in Seurat (Seurat version 4.1.0, RRID: SCR_007322; R version 4.3.3, RRID: SCR_001905). Low-quality cells were excluded based on UMI count (<2,500), detected genes (<500), and mitochondrial transcript fraction (>20%). Doublets were removed using DoubletFinder (DoubletFinder version 3, RRID: SCR_018771). Data were normalized using SCTransform (version 0.4.1, RRID: SCR_022146), regressing out cell cycle and mitochondrial effects, and integrated using Harmony (version 1.2.0, RRID:SCR_022206). Clustering was performed using a shared nearest-neighbor graph based on the first 40 principal components, followed by UMAP visualization. Cluster identities were assigned using conserved marker genes. Differential expressions within each cell type was assessed using the MAST test with Bonferroni correction. Gene set enrichment and pathway analyses were conducted on differentially expressed genes using standard functional enrichment tools. Cell-cell communication was analyzed using CellChat (version 2.1.0, RRID: SCR_021946). Additional details are provided in Supplementary Methods.

### Autophagic flux analysis

Autophagic flux in iPSC-derived microglia was assessed by treatment with 100□nM bafilomycin A1 (Sigma-Aldrich, Cat. # B1793) for 4 hours prior to protein extraction. LC3B levels were analyzed by Western blot, and the LC3-II band intensity was quantified using ImageJ. Flux was calculated as the ratio of LC3-II in bafilomycin-treated cells versus untreated cells. Ratios were compared between control and CDDO-Me-treated conditions to evaluate autophagosome accumulation.

### α-Synuclein pre-formed fibril treatment

Recombinant human α-synuclein was expressed, purified, and assembled into pre-formed fibrils (PFFs) as previously described ^79^, with fibril formation validated by biochemical and ultrastructural assays (details in Supplementary Methods). Prior to use, PFFs were sonicated to generate short fibrillar species and added to iPSC-derived microglia at a final concentration of 1 µg/mL. Where indicated, cells were co-treated with the indicated compounds. α-Synuclein uptake and intracellular processing were assessed by immunofluorescence and live-cell imaging at defined time points. Particle quantification was performed using automated object detection and tracking using CellProfiler (version 4.2.6, RRID:SCR_007358).

### Statistics and reproducibility

Statistical analyses were performed using GraphPad Prism (version 10.2.3, RRID: SCR_002798). No statistical methods were used to predetermine sample sizes; sample sizes were comparable to those reported in previous iPSC-based studies. No formal randomization was performed. Cell cultures were assigned to experimental conditions based on availability and processed in parallel to minimize batch effects. Animals were grouped by genotype and age. Analyses were conducted blinded when feasible (e.g., omics and automated image analyses using coded samples) but were otherwise unblinded. Samples were excluded if identified as statistical outliers or in cases of predefined technical failure. Data were assumed to follow a normal distribution but were not formally tested. Two-group comparisons were conducted using unpaired two-tailed Student’s t-tests, and multiple-group comparisons were analyzed by ANOVA with appropriate multiple-comparison corrections, as specified in the figure legends. Data are presented as mean ± SEM from at least two independent differentiations with technical replicates.

### Data Availability

All data supporting the findings of this study are available within the article and its supplementary information files. Bulk and single-cell RNA-sequencing datasets generated in this study have been deposited in the Gene Expression Omnibus (GEO) under accession numbers GSE274283 and GSE274289. Detailed experimental protocols are available on protocols.io at https://dx.doi.org/10.17504/protocols.io.8epv5kbm4v1b/v1. Source data for all main and supplementary figures are provided in the accompanying files Source Data. The full collection of datasets, images, and custom analysis scripts used in this study has been deposited on Zenodo under DOI 10.5281/zenodo.17294146, 10.5281/zenodo.17831119, 10.5281/zenodo.17608082, 10.5281/zenodo.17294305 and 10.5281/zenodo.17294798. Previously published datasets used for comparative analyses are detailed in the Methods section with appropriate references.

