## Supplementary Figures and Text for "The mitochondrial unfolded protein response in human microglia disrupts neuronal-glial communication and promotes senescence"

- **Supplemental Figures 1-16. Page 1 - 40**
- **Supplemental Methods. Page 41 - 54**
- **Supplemental Tables. Page 55 - 58**
- **Supplemental Unprocessed Blots. Page 59 - 68**

#### Supplemental Figures

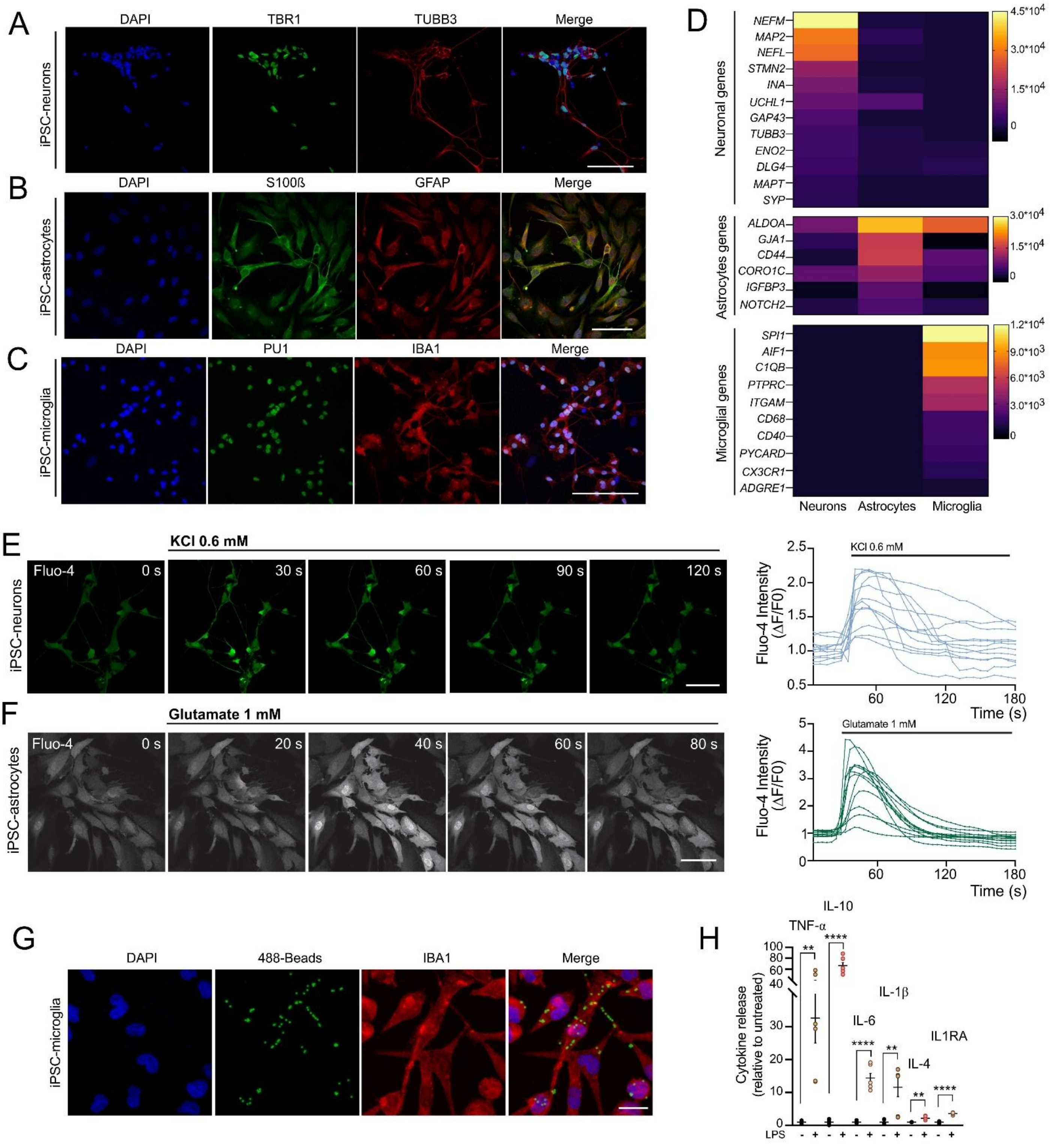


Supplementary Figure 1

**Supplementary Figure 1. Characterization of human iPSC-derived neurons and glia**. **A-C)** Representative confocal images of immunofluorescence staining showing cell type-specific markers in human iPSC-derived (A) cortical neurons (TBR1, green; TUBB3, red), (B) astrocytes (S100ß, green; GFAP, red), and (C) microglia (PU1, green; IBA1, red) (C1). Nuclei were counterstained with DAPI (blue). Scale bars, 50 μm. **D)** Heatmap of bulk RNA-sequencing data showing mean read counts of cell type-specific genes in human iPSC-derived neurons, astrocytes, and microglia. **E-F)** Representative confocal images of intracellular calcium dynamics measured in live cells using the Fluo-4 dye. Scale bars, 50 μm. Human iPSC-cortical neurons were stimulated with 0.6 mM KCl (E), and human iPSC-astrocytes were stimulated with 1 mM glutamate (F). Right: fluorescence intensity traces from individual cells over time. Images were acquired every 6 s (neurons) or 4 s (astrocytes) for 3 min. Fluorescence values were normalized to baseline prior to stimulation. n = 2 independent differentiations (C1). **G)** Representative confocal images showing microglial phagocytosis of fluorescent latex beads (green) with IBA1 immunostaining (red). Scale bar, 10 μm. **H)** Cytokine levels in the supernatant from human iPSC-derived microglia, either untreated or treated with lipopolysaccharide (LPS) for 24 hours. Mean ± SEM; unpaired two-tailed *t* test compared with the untreated group, $^{**}P = 0.0019$ (TNF-α), 0.0042 (IL-1β), 0.0020 (IL-4), and $^{****}P < 0.0001$; n = 6 independent experiments (C1).

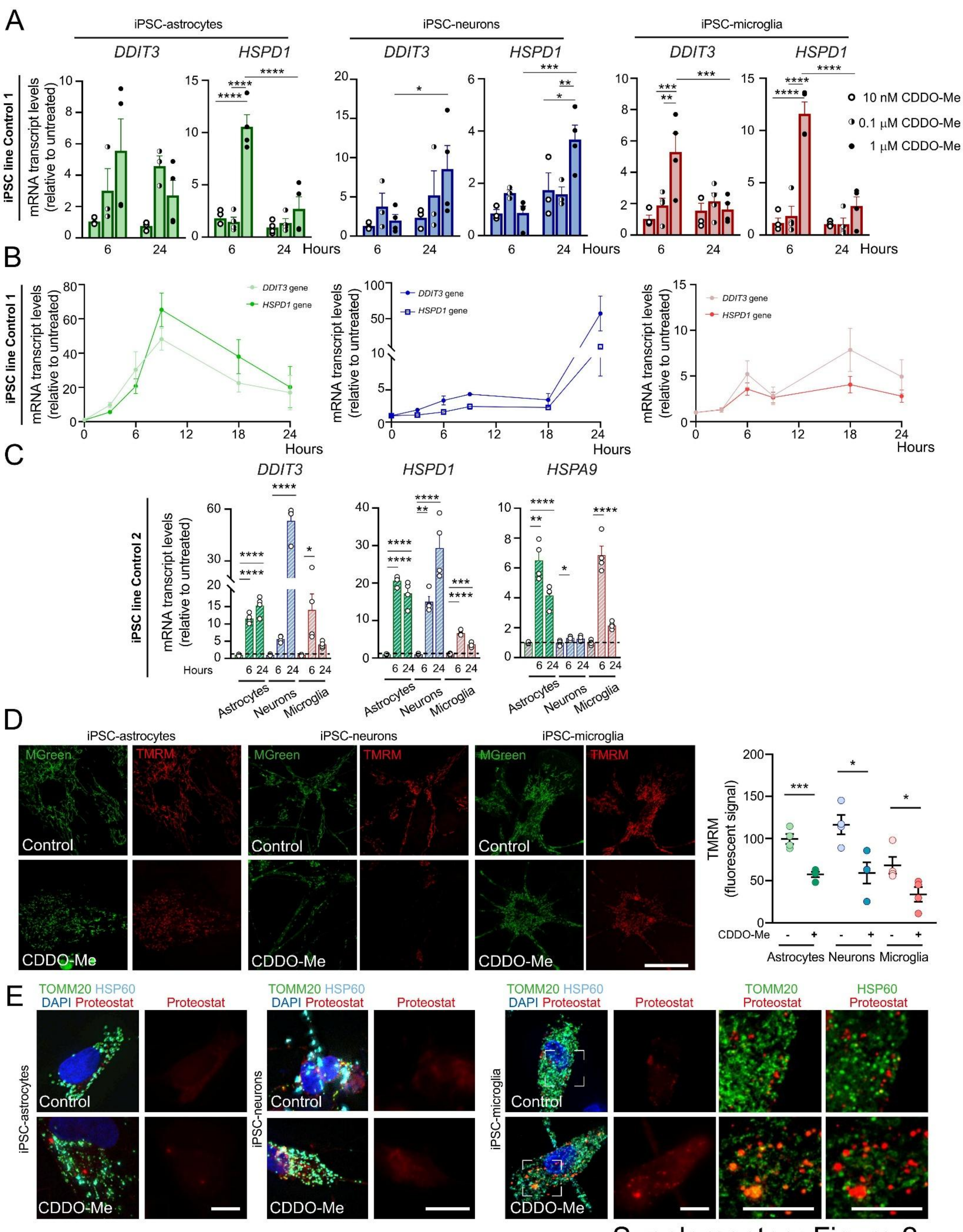


Supplementary Figure 2

**Supplementary Figure 2. Characterization of UPR$^{mt}$ activation in human iPSC-derived neurons and glia. A-C)** mRNA expression levels of the UPR$^{mt}$ genes *HSPD1* and *DDIT3* in human iPSC-derived microglia, astrocytes, and neurons. **A)** mRNA levels measured after 6 hours and 24 hours of CDDO-Me treatment at three different concentrations. Mean ± SEM; two-way ANOVA with Bonferroni *post hoc* correction. Astrocytes: ****$P$ < 0.0001; Neurons: *$P$ = 0.0267 (*DDIT3*), *$P$ = 0.0130 (*HSPD1*), **$P$ = 0.0073, ***$P$ = 0.0001; Microglia: **$P$ = 0.0037, ***$P$ = 0.0004 (*DDIT3,* 6 hours, 0.01 versus 1 µM), 0.0007 (*DDIT3,* 1 µM, 6 hours versus 24 hours), and ****$P$ < 0.0001; astrocytes and neurons: n = 3 at 10 nM and, 0.1 µM and n = 4 at 1 µM CDDO-Me; microglia: n = 4 at 10 nM, 0.1 µM and 1 µM CDDO-Me independent experiments (C1). **B)** mRNA levels after 1 µM CDDO-Me treatment at different time points. Mean ± SEM; n = 4 (astrocytes and neurons), and 8 (microglia) independent experiments (C1). **C)** mRNA levels of *HSPD1, HSPA9* and *DDIT3* after treatment with 1 µM CDDO-Me for 6 hours or 24 hours. Mean ± SEM, with the data normalized to those of the control (gray) in each condition; one-way ANOVA with Bonferroni *post hoc* correction. *DDIT3*: *$P$ = 0.0159, ****$P$ < 0.0001; *HSPD1:* **$P$ = 0.0060, ***$P$ = 0.0002, and ****$P$ < 0.0001; *HSPA9*: *$P$ = 0.0392, **$P$ = 0.0010, ****$P$ < 0.0001; n = 4 independent experiments (C2). **D)** Representative confocal images showing mitochondrial morphology (Mitogreen, green) and mitochondrial membrane potential (TMRM, red) in control and CDDO-Me-treated human iPSC-derived astrocytes, neurons, and microglia (C1). Changes in TMRM fluorescence intensity are shown in the right panel. Mean ± SEM; unpaired two-tailed *t* test; *$P$ = 0.0153 (neurons), 0.0400 (microglia) and ***$P$ = 0.0008. n = 4 independent experiments. Scale bar, 10 µm. **E)** Representative images showing immunofluorescence staining for TOMM20 (green) and HSP60 (cyan) in human iPSC-derived neurons and glia (C2). PROTEOSTAT (red) and DAPI (blue) staining are shown. Right panels display high-magnification areas in microglia with the colocalization of PROTEOSTAT (red), TOMM20 (green), and HSP60 (green). Scale bar, 10 µm.

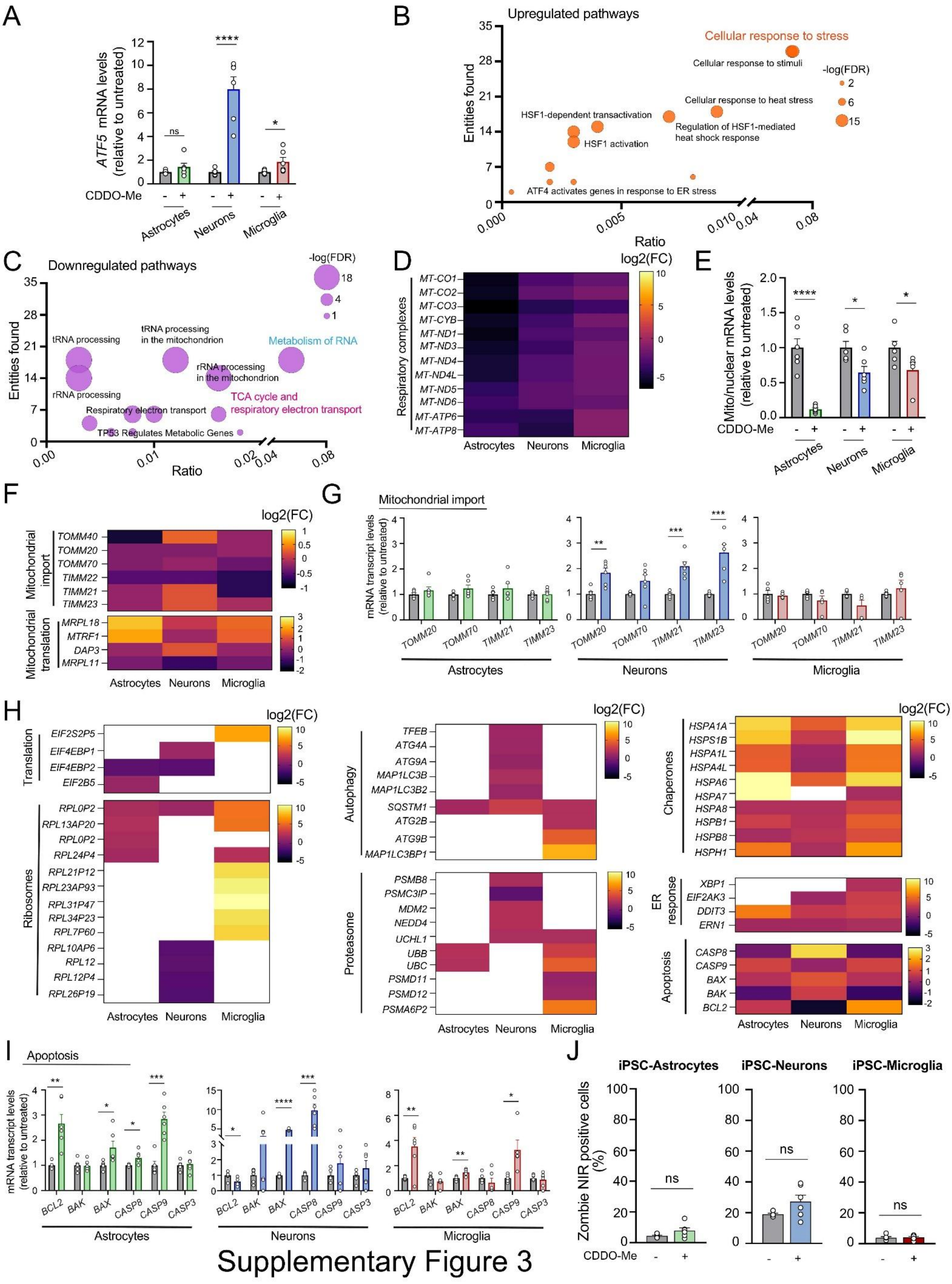


Supplementary Figure 3

**Supplementary Figure 3. The UPR$^{mt}$ elicits mitochondrial dysfunction and stress-dependent chaperone induction. A)** mRNA expression levels of the UPR$^{mt}$ gene *ATF5* in human iPSC-derived microglia, astrocytes, and neurons after treatment with 1 µM CDDO-Me. Mean ± SEM; the data are normalized to control (grey) in each condition; unpaired two-tailed *t* test, **P* = 0.0495, *****P* < 0.0001 and ns = not significant; n = 6 independent experiments (C1). **B, C)** Bubble plots showing the top Reactome pathways that were upregulated (B) or downregulated (C) after CDDO-Me treatment, shared among neurons, astrocytes, and microglia. Bubble diameter represents -log10(FDR). **D)** Heatmap of bulk RNA-sequencing data illustrating mean read counts of respiratory complex subunit genes in human iPSC-derived astrocytes, neurons and microglia. **E)** Ratios of the mRNA expression levels of mitochondrial genome-encoded (*mt-ND1*) and nuclear genome-encoded (*NDUF2*) complex I subunits in human iPSC-derived astrocytes, neurons and microglia after treatment with 1 µM CDDO-Me. Mean ± SEM; data are normalized to the control in each condition; unpaired two-tailed *t* test; **P* = 0.0177 (neurons), 0.0269 (microglia) and *****P* < 0.0001; n = 6 independent experiments (C1). **F)** Heatmap of bulk RNA-sequencing data illustrating mean read counts of mitochondrial genes in human iPSC-derived astrocytes, neurons and microglia. **G)** mRNA expression levels of mitochondrial import genes in human iPSC-derived astrocytes, neurons and microglia after treatment with 1 µM CDDO-Me. Mean ± SEM; data are normalized to control in each condition; unpaired two-tailed *t* test; ***P* = 0.0020 and ****P* = 0.0002 (*TIMM21*), 0.0004 (*TIMM23*); n = 5 (microglia) or n = 6 (neurons, astrocytes) independent experiments (C1). **H)** Heatmap of bulk RNA-sequencing data illustrating mean read counts of cellular protein stress response and apoptosis genes in human iPSC-derived astrocytes, neurons and microglia. **I)** mRNA expression levels of apoptotic genes in human iPSC-derived astrocytes, neurons and microglia after treatment with 1 µM CDDO-Me. Mean ± SEM; data normalized to control in each condition; unpaired two-tailed *t* test. Astrocytes: **P* = 0.0241 (*BAX*), 0.0293 (*CASP8*), ***P* = 0.0013, ****P* = 0.0002; Neurons: **P* = 0.0225, ****P* = 0.0008, *****P* < 0.0001; microglia: **P* = 0.0213, ***P* = 0.0064 (*BCL2*), 0.0024 (*BAX*); n = 6 independent experiments (C1). **J)** Percentage of Zombie NIR-positive cells analyzed by flow cytometry in untreated and

1 μM CDDO-Me-treated iPSC astrocytes (6 h), neurons (24 h) and microglia (6 h). Mean ± SEM; unpaired two-tailed *t* test, ns = not significant; n = 6 independent experiments (C2).

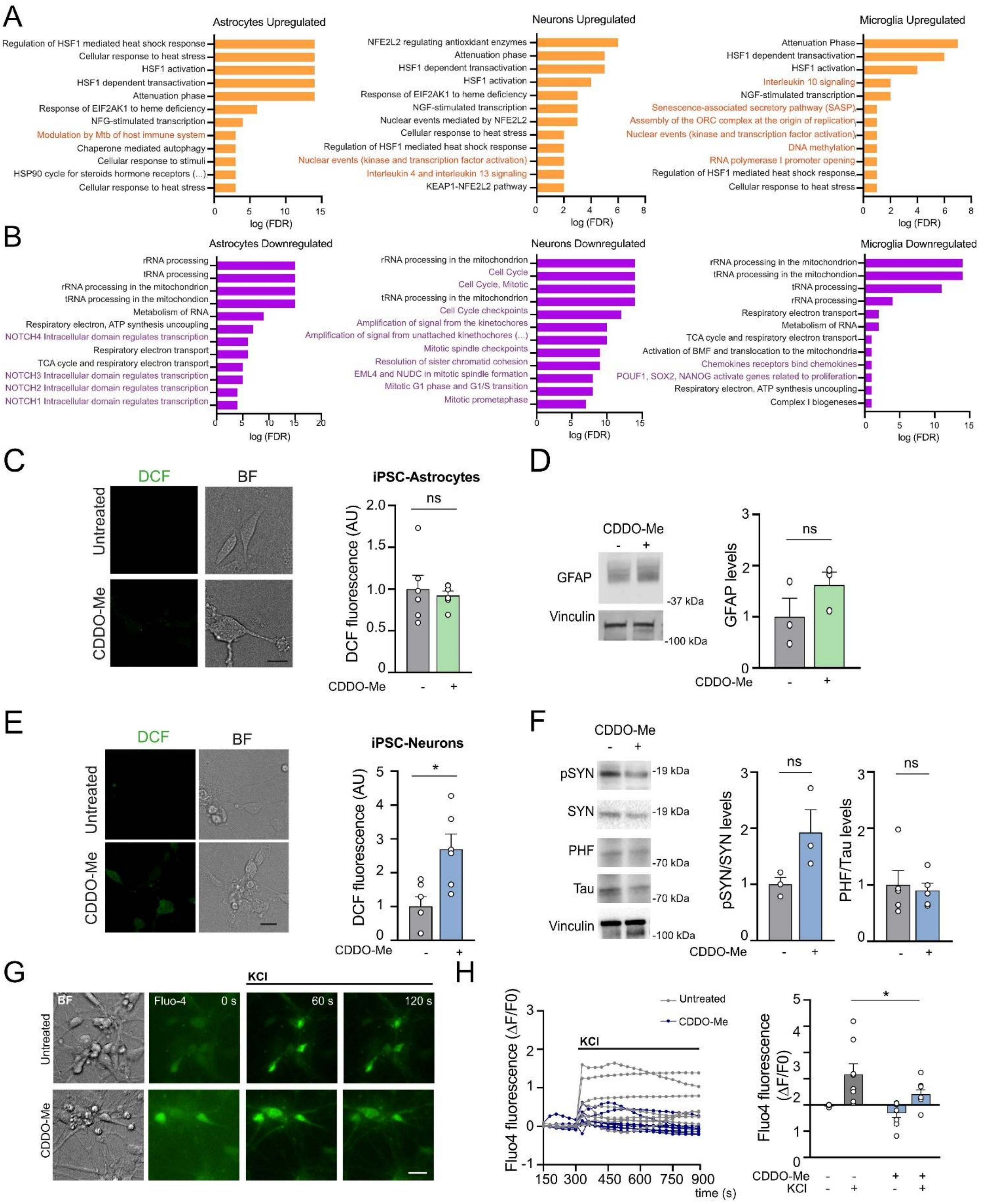


Supplementary Figure 4

**Supplementary Figure 4. LONP1 inhibition induces a cell type-specific response in iPSC-derived neurons and glia. A, B)** Top enriched Reactome pathways of upregulated (A) and downregulated (B) differentially expressed genes (DEGs) in human iPSC-derived microglia, neurons, and astrocytes. Cell type-specific pathways are highlighted in orange (A) and purple (B). **C)** Live-cell imaging of 2, 7-Dichlorofluorescin (DCF) fluorescence as a measure of reactive oxygen species in untreated and 1 µM CDDO-Me-treated iPSC-derived astrocytes. BF: bright field. Scale bar, 10 µm. Quantification of fluorescence intensity is shown on the right. Mean ± SEM; unpaired two-tailed *t* test; ns = not significant; n = 6 independent experiments (C2). **D)** Representative Western blot showing GFAP levels in untreated and 1 µM CDDO-Me-treated iPSC-derived astrocytes. Vinculin served as a loading control. Right, quantification of GFAP levels normalized to vinculin and to untreated control. Mean ± SEM; unpaired two-tailed *t* test; ns = not significant; n = 3 independent experiments (C2). **E)** Live-cell DCF imaging in untreated and 1 µM CDDO-Me-treated iPSC-derived neurons. BF: bright field. Scale bar, 10 µm. Quantification is shown on the right. Mean ± SEM; unpaired two-tailed *t* test; *$P$ = 0.0101; n = 6 independent experiments (C2). **F)** Representative immunoblot showing phosphorylated α-synuclein (pSYN; Ser129), total α-synuclein (SYN), phosphorylated tau (PHF; Thr181), total tau, and vinculin (loading control) in untreated and 1 µM CDDO-Me-treated iPSC-derived neurons (24 h). Right, quantification of immunoblot data showing pSYN/total SYN and PHF/total tau ratios normalized to control. Mean ± SEM; unpaired two-tailed *t* test; ns = not significant; n = 3 (pSYN/total SYN), and 5 (PHF/total tau) independent experiments (C2). **G, H)** Live-cell calcium imaging using Fluo-4 in human iPSC-derived neurons stimulated with 100 mM KCl. Images were acquired every 30 s for 15 min. BF: bright field. Scale bar, 10 µm. Representative images are shown in G; single-cell fluorescence traces over time are shown in H. Fluorescence intensity was normalized to baseline prior to stimulation. Right, quantification of calcium responses before and after KCl stimulation. Mean ± SEM; the data are normalized to the control in each condition; two-way repeated-measures ANOVA with Bonferroni *post hoc* correction, *$P$ = 0.0448; n = 8 cells from 2 independent experiments (C2).

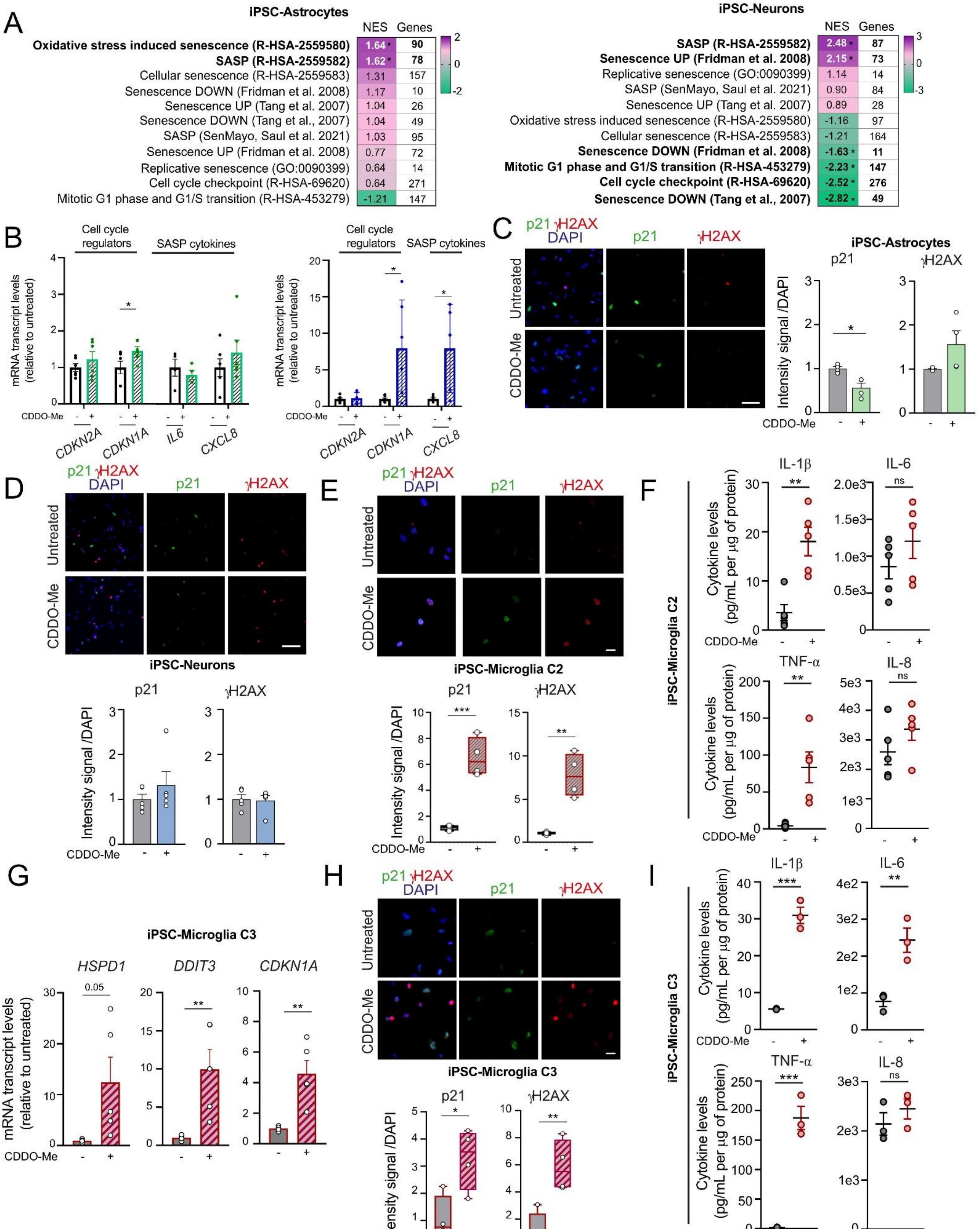


Supplementary Figure 5

**Supplementary Figure 5. CDDO-Me treatment induces a senescence-associated phenotype in human iPSC-derived microglia. A)** Heatmap showing relative enrichment of senescence-associated gene sets from RNA-sequencing data of human iPSC-derived neurons and astrocytes. Normalized enrichment scores (NESs) for treated versus untreated cells were calculated using Gene Set Enrichment Analysis. Upregulation is shown in purple and downregulation is shown in green. *FDR $\leq$ 0.05. **B)** mRNA expression levels of cell cycle regulators (*CDKN2A*, *CDKN1A*) and SASP cytokines (*IL6* and *CXCL8*) in human iPSC-derived astrocytes (green) and neurons (blue) after 1 µM CDDO-Me treatment. Mean ± SEM; data normalized to untreated controls; unpaired two-tailed *t* test; **P* = 0.0498 (*CDKN1A*, astrocytes), 0.0280 (*CDKN1A*, neurons), 0.0199 (*CXCL8*, neurons); astrocytes: n = 6 (*CDKN2A, CDKN1A, IL8*), n = 4 (*IL6*); neurons: n = 6 (*CDKN1A, CDKN2A, IL8*) independent experiments (C1). **C, D)** Representative immunofluorescence images of p21 (green) and γH2AX (red) in untreated and CDDO-Me-treated human iPSC-derived astrocytes (C) and neurons (D). Nuclei were stained with DAPI (blue). Scale bar, 50 µm. Right, quantification of p21- and γH2AX-positive cells normalized to DAPI-positive nuclei. Mean ± SEM; unpaired two-tailed *t* test; **P* = 0.0101; n = 4 (C) and n = 5 (D) independent experiments (C2). **E)** Representative immunofluorescence images of p21 (green) and γH2AX (red) in untreated and CDDO-Me-treated microglia. Nuclei were stained with DAPI (blue). Scale bar, 10 µm. Below, quantification of fluorescence intensity normalized to control. Center line, median; box, 25th–75th percentiles; whiskers, min–max; data are normalized to the control; unpaired two-tailed *t* test; ***P* = 0.0018 and ****P* = 0.0004; n = 4 independent experiments (C2). **F)** Cytokine levels in the supernatants of untreated or CDDO-Me-treated microglia (6 hours). Mean ± SEM; unpaired two-tailed *t* test; ***P* = 0.0024 (IL-1β), 0.0053 (TNF-α); ns = not significant; n = 5 independent experiments (C2). **G)** mRNA expression levels of $UPR^{mt}$ genes (*HSPD1, DDIT3*), and the cell cycle regulator *CDKN1A* in human iPSC-derived microglia after treatment with 1 µM CDDO-Me. Mean ± SEM; normalized to untreated control in each condition; unpaired two-tailed *t* test; ***P* = 0.0096 (*DDIT3*), 0.0034 (*CDKN1A*); n = 5 independent experiments (C3). **H)** Representative confocal images of p21 (green) and γH2AX (red) in untreated and CDDO-

Me-treated human iPSC-derived microglia for (6 hours, C3). Nuclei were stained with DAPI (blue). Scale bar, 10 μm. Below, quantification of fluorescence intensity normalized to DAPI-positive nuclei. Center line, median; box, 25th–75th percentiles; whiskers, min–max; unpaired two-tailed *t* test; **P* = 0.0182, ***P* = 0.0059; n = 4 independent experiments. **I)** Cytokine levels in the supernatants from untreated and CDDO-Me-treated microglia (6 hours, C3). Mean ± SEM; unpaired two-tailed *t* test; ***P* = 0.0099 and ****P* = 0.0003 (IL-1β), 0.0007 (TNF-α); ns = not significant; n = 3 independent experiments (C3).

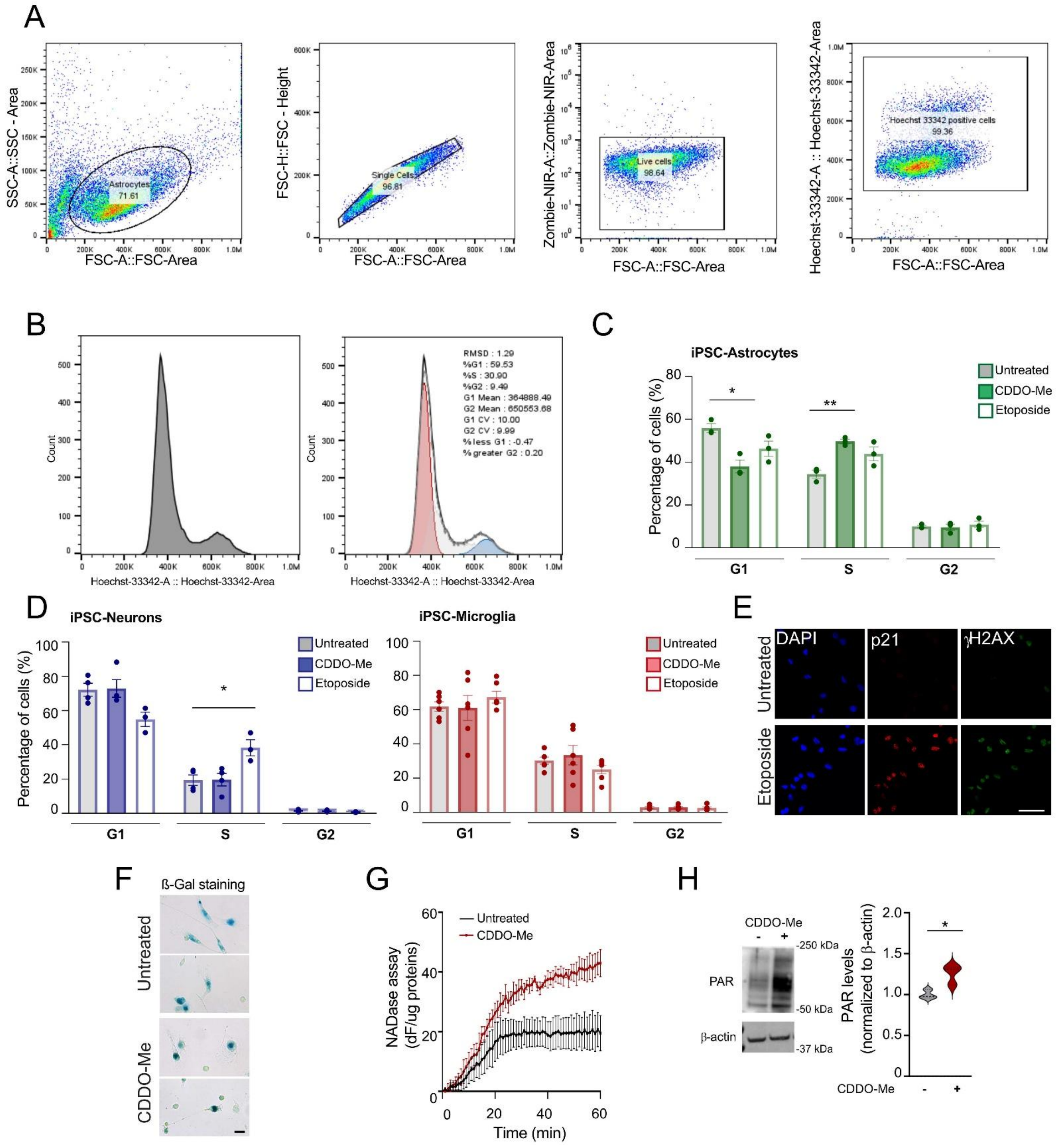


Supplementary Figure 6

**Supplementary Figure 6. Characterization of the senescence signature and phenotype of human iPSC-derived neurons and glia. A)** Representative flow cytometry gating strategy in astrocytes. Cells were first gated based on forward scatter area (FSC-A) versus side scatter area (SSC-A), followed by doublet exclusion using FSC-A versus FSC-H. Live cells were selected using FSC-A versus Zombie NIR-A, and Hoechst 33342-positive nuclei were subsequently gated. **B)** Left, representative input histogram for cell cycle analysis showing Hoechst 33342-A intensity versus cell count (astrocyte control). Right, corresponding cell cycle distribution using the Watson (Pragmatic) model: G1 (red), S (light gray), and G2 (blue). The model fit is shown in dark gray with the raw data overlaid in black. **C, D)** Quantification of the percentage of cells in G1, S, and G2 phases in iPSC-derived astrocytes, neurons and microglia under the indicated treatment conditions. Mean ± SEM; one-way ANOVA with Bonferroni *post hoc* correction; astrocytes: **P* = 0.0141, ***P* = 0.0085, n = 3; neurons: **P* = 0.0455, n = 3; microglia: n = 6 independent experiments (C2). **E)** Representative images of iPSC-derived microglia (C2) under untreated condition or after 10 µM etoposide treatment (6 hours). Cells were stained for p21 (red), gH2AX (green), and DAPI (blue). Scale bar, 10 µm. **F)** Representative brightfield images of ß-galactosidase (ß-Gal) staining in human iPSC-derived microglia (C1) treated with or without 1 µM CDDO-Me for 6 hours. Images are representative of at least two independent experiments. Scale bar, 10 µm. **G)** Median fluorescence intensity of a NAD fluorescent analog measured over time in untreated and CDDO-Me-treated microglia. Mean ± SEM; data are normalized to total protein content in each sample; n = 3 independent experiments (C1). **H)** Representative immune blot showing poly (ADP-ribose) (PAR) and β-actin levels (loading control) in human iPSC-derived microglia untreated and treated with 1 µM CDDO-Me for 6 hours. Right, densitometric quantification normalized to control (gray). Violin plots indicate medians; unpaired two-tailed *t* test; **P* = 0.0280; n = 3 independent experiments (C1).

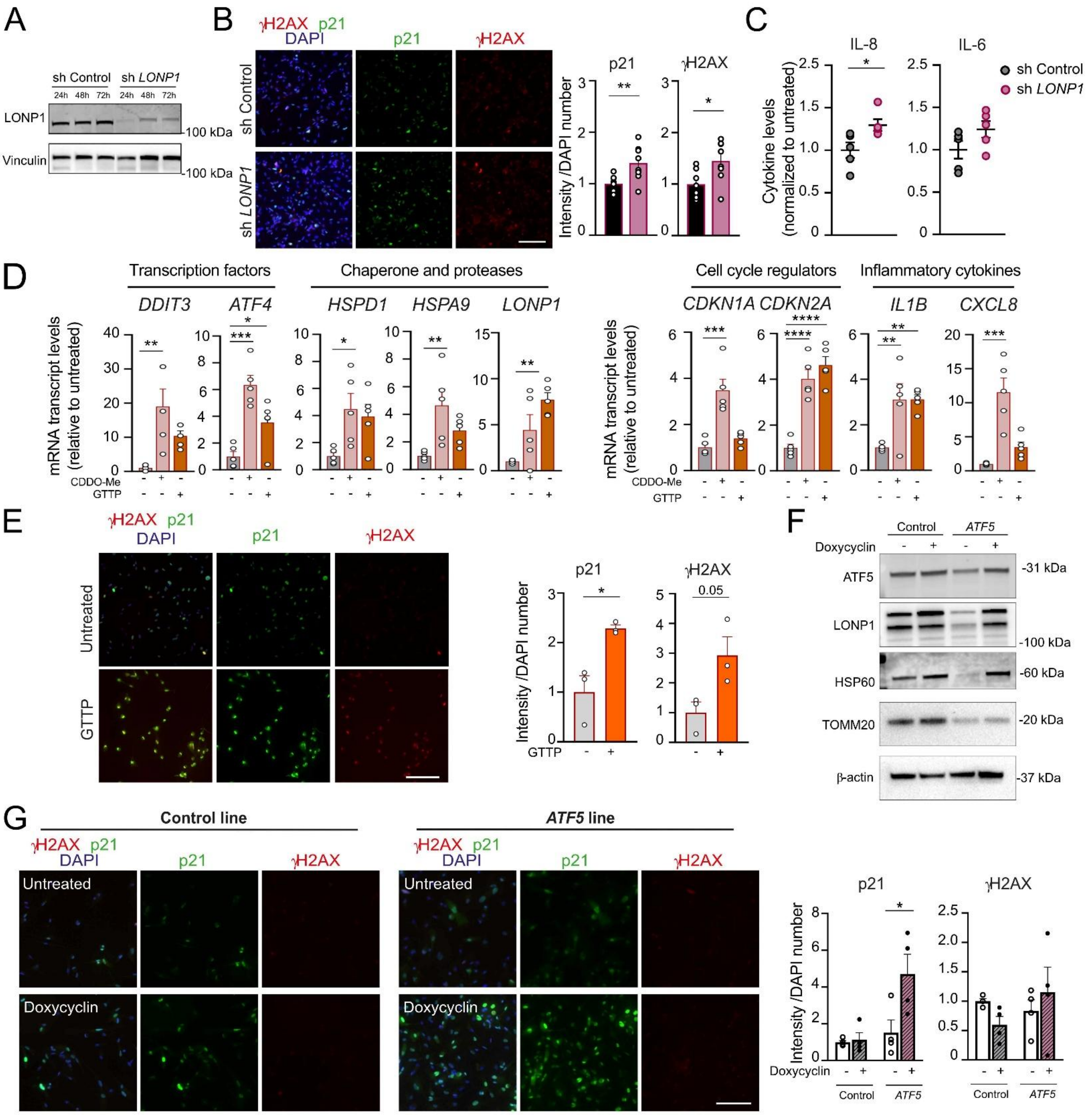


Supplementary Figure 7

**Supplementary Figure 7. Pharmacological and genetic induction of mitochondrial protein stress triggers a senescence-associated phenotype in human iPSC-derived microglia. A)** Representative Western blot showing LONP1 protein levels in iPSC-derived microglia at 24, 48 and 72 hours after infection with a lentivirus carrying a control or *LONP1*-targeting short hairpin RNA (shRNA). Vinculin serves as loading control. **B)** Representative immunofluorescence images of p21 (green) and γH2AX (red) in iPSC-derived microglia 48 hours after infection with control or *LONP1*-targeting shRNA. Nuclei were stained with DAPI (blue). Scale bar, 100 μm. Right, quantification of fluorescence intensity normalized to control. Mean ± SEM; unpaired two-tailed *t* test, $^{**}P = 0.0092$ and $^{*}P = 0.0173$; n = 4 technical replicates in 2 independent experiments (C2). **C)** Cytokine levels in supernatants from iPSC-derived microglia at 48 hours after infection with a lentivirus carrying a control or *LONP1*-targeting shRNA. Mean ± SEM; unpaired two-tailed *t* test; $^{*}P = 0.0332$; n = 5 independent experiments (C2). **D)** mRNA expression levels of $UPR^{mt}$ genes (*DDIT3*, *ATF4*, *HSPD1*, *HSPA9*, *LONP1*), cell cycle regulators (*CDKN1A*, *CDKN2A*) and SASP cytokines (*IL1B, CXCL8*) in human iPSC-derived microglia after treatment with 1 μM CDDO-Me or 1 μM GTTP for 6 hours. Mean ± SEM; data are normalized to the untreated control in each condition; one-way ANOVA with Bonferroni *post hoc* correction. Sequentially: $^{**}P = 0.0030$ (*DDIT3*), $^{*}P = 0.0407$ (*ATF4*), $^{***}P = 0.0002$ (*ATF4*), $^{*}P = 0.0310$ (*HSPD1*), $^{**}P = 0.0092$ (*HSPA9*), $^{**}P = 0.0016$ (*LONP1*), $^{***}P = 0.0002$ (*CDKN1A*), $^{****}P < 0.0001$ (*CDKN2A*), $^{**}P = 0.0090$ (*IL1B,* control versus CDDO-Me), 0.0088 (*IL1B*, control versus GTTP), $^{***}P = 0.0002$ (*CXCL8*); n = 5 independent experiments (C2). **E)** Representative immunofluorescence images of p21 (green) and γH2AX (red) in iPSC-derived microglia untreated, and after treatment with GTTP. Nuclei were stained with DAPI (blue). Scale bar, 100 μm. Right, quantification of fluorescence intensity normalized to control. Mean ± SEM; unpaired two-tailed *t* test, $^{*}P = 0.0188$; n = 3 independent experiments (C2). **F)** Representative Western blot showing ATF5, LONP1, HSP60, and TOMM20 protein levels in control and *ATF5*-overexpressing iPSC-derived microglia treated with or without doxycycline for 24 hours. Actin serves as loading control. **G)** Representative immunofluorescence images of p21 (green) and γH2AX (red) in control and

*ATF5*-overexpressing iPSC-derived microglia treated with or without doxycycline for 24 hours. Nuclei were stained with DAPI (blue). Scale bar, 50 μm. Right, quantification of fluorescence intensity normalized to control. Mean ± SEM; the data are normalized to the control in each condition; Two-way ANOVA with Bonferroni *post hoc* correction between genotypes at each time point, $^*P = 0.0108$; n = 4 independent experiments (C2).

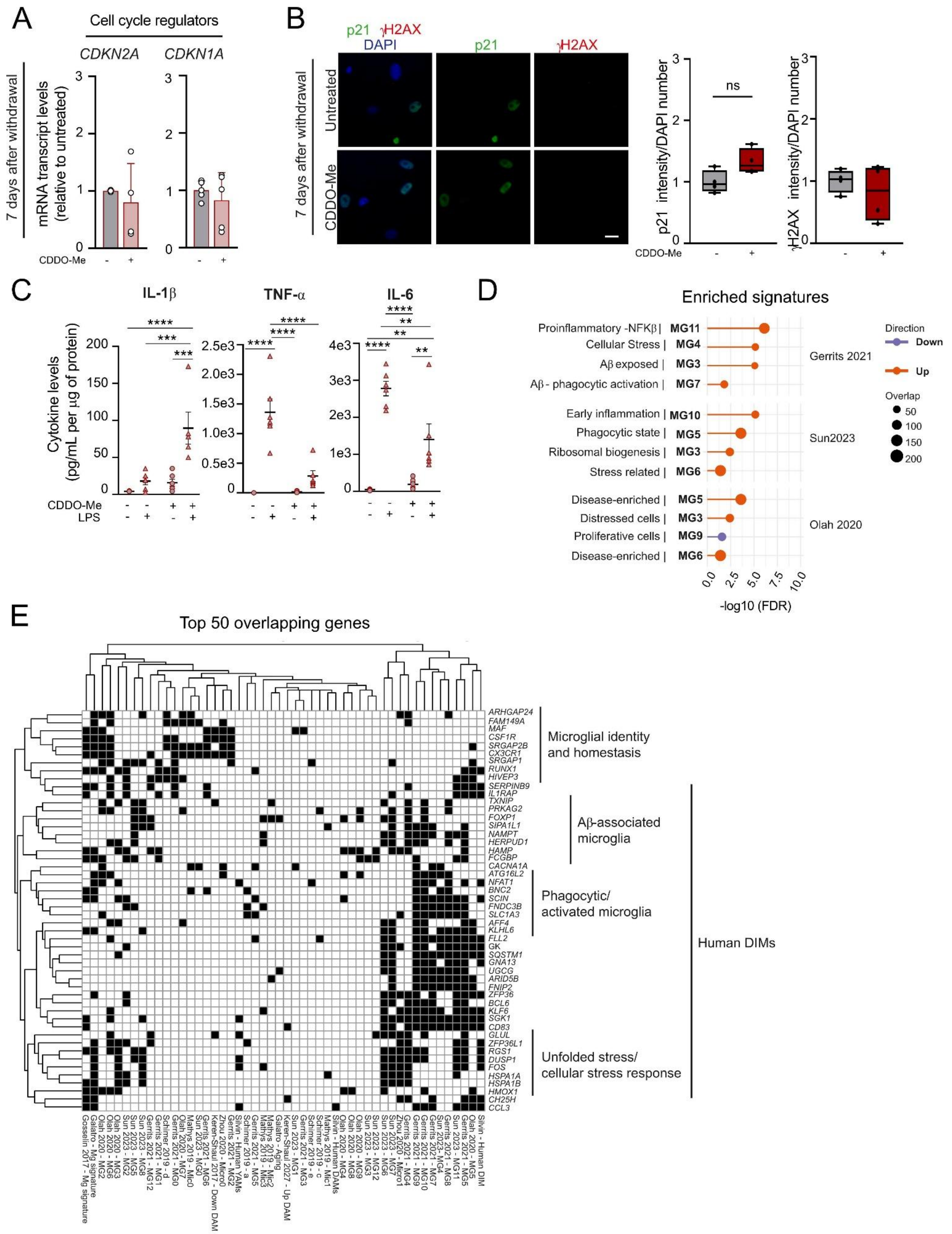


### Supplementary Figure 8

**Supplementary Figure 8. CDDO-Me treatment induces a neurodegenerative disease-associated transcriptional signature in human iPSC-derived microglia. A)** mRNA expression levels of cell cycle regulators (*CDKN2A*, *CDKN1A*) in human iPSC-derived microglia untreated and treated with 1 µM CDDO-Me for 6 hours followed by a 7-day washout period. Mean ± SEM, normalized to untreated controls; n = 4 (*CDKN2A*) and 5 (*CDKN1A*) independent experiments (C2). **B)** Representative immunofluorescence images of p21 (green) and γH2AX (red) in human iPSC-derived microglia, either untreated or treated with CDDO-Me for 6 hours followed by a 7-day washout period. Nuclei were counterstained with DAPI (blue). Scale bar, 10 µm. Right, quantification of fluorescence intensity normalized to DAPI-positive nuclei. Center line, median; box, 25th–75th percentiles; whiskers, min–max; unpaired two-tailed *t*-test; ns = not significant; n = 4 independent experiments (C2). **C)** Cytokine levels in supernatants from human iPSC-derived microglia, either untreated or treated with CDDO-Me, LPS, or CDDO-Me + LPS. Mean ± SEM; two-way ANOVA with Bonferroni *post hoc* comparisons. IL-1β: ***$P$ = 0.0004 (CDDO-Me versus CDDO-Me ± LPS), 0.0005 (LPS versus CDDO-Me + LPS), and ****$P$ < 0.0001; TNF-α: ****$P$ < 0.0001; IL-6: **$P$ = 0.0034 (untreated versus CDDO-Me + LPS), 0.0029 (LPS versus CDDO-Me + LPS), 0.0092 (CDDO-Me versus CDDO-Me ± LPS), and ****$P$ < 0.0001; n = 6 independent experiments (C2). **D)** Bubble plot showing -log10(FDR) values from Fisher's exact test comparing genes upregulated in CDDO-Me-treated microglia with annotated clusters from the Human Microglia Atlas (HuMicA). Clusters represent microglial states identified in aging and neurodegenerative human brain datasets. **E)** Heatmap displaying the presence (black) or absence (white) of the top 50 genes overlapping between the CDDO-Me signature and significantly enriched microglial states from human datasets in the HuMicA database.

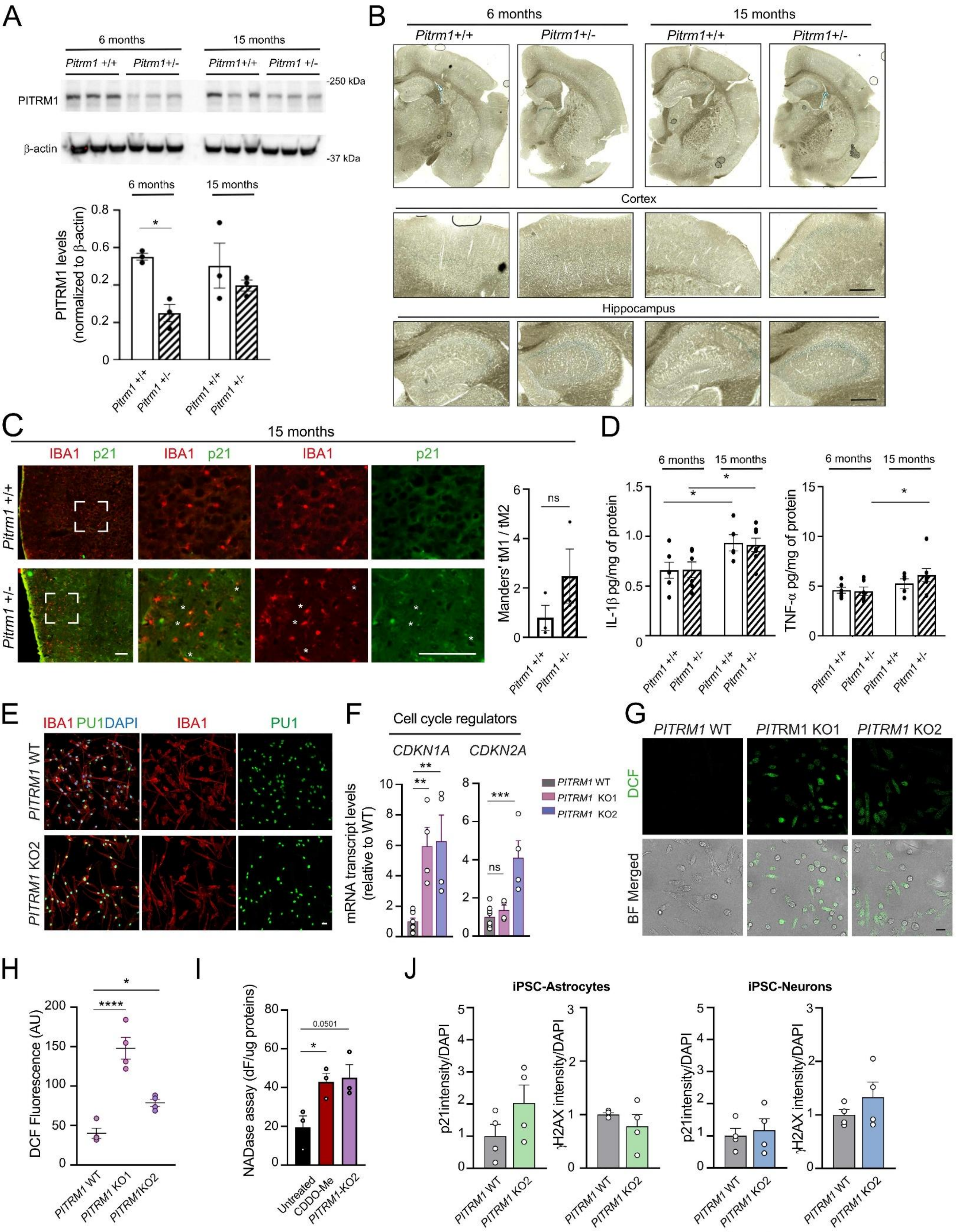


Supplementary Figure 9

**Supplementary Figure 9. *PITRM1* loss of function in mouse and human iPSC-derived microglia is associated with a senescence-like phenotype. A)** Representative Western blot showing PITRM1 protein levels in cortical lysates from *Pitrm1*$^{+/-}$ and *Pitrm1*$^{+/+}$ mice. Bottom, densitometric quantification normalized to β-actin in each sample. Mean ± SEM; two-way ANOVA with Fisher's LSD test, *$P$ = 0.0126; n = 3 animals per condition. **B)** Representative brightfield images showing ß-galactosidase staining in *Pitrm1*$^{+/-}$ and *Pitrm*$^{+/+}$ mice at 6 months and 15 months of age. Whole brains are shown in the upper panels (scale bar, 1,000 µm); higher magnification views of cortex and hippocampus are shown below (scale bar, 500 µm). Images are representative of three animals per genotype. **C)** Representative confocal images of p21 (green) and IBA1 (red) in the cortices of *Pitrm1*$^{+/-}$ and *Pitrm1*$^{+/+}$ mice at 15 months of age. White asterisks indicate p21-positive microglia. Scale bar, 10 µm. Right, quantification of Mander's colocalization ratio for p21 and IBA1 in cortical microglia. Mean ± SEM; unpaired two-tailed *t* test, ns = not significant; n=3 animals per condition. **D)** ELISA quantification of interleukin-1 beta (IL-1β) and tumor necrosis factor alpha (TNF-α) levels in total brain homogenates from *Pitrm1*$^{+/-}$ and *Pitrm1*$^{+/+}$ mice at 6 and 15 months of age. Mean ± SEM; two-way ANOVA with Fisher's LSD test, *$P$ = 0.0134 (IL-1β, *Pitrm1*$^{+/+}$ 6 versus 15 months), 0.0481 (IL-1β, *Pitrm1*$^{+/-}$ 6 versus 15 months), 0.0242 (TNF-α, *Pitrm1*$^{+/-}$ 6 versus 15 months). n = 6 animals per genotype per age group. **E)** Representative confocal images of microglia differentiated from *PITRM1*-wild type (WT) and *PITRM1*-knockout (clone KO2) human iPSCs. IBA1 (red), PU1 (green), and DAPI (blue) are shown. Scale bar, 10 µm. **F)** mRNA expression levels of cell cycle regulators (*CDKN1A*, *CDKN2A*) in microglia derived from *PITRM1*-WT and *PITRM1*-KO human iPSCs (KO1, KO2). Mean ± SEM; data are normalized to WT control; one-way ANOVA with Bonferroni *post hoc* correction, **$P$ = 0.0060 (WT versus KO1), 0.0037 (WT versus KO2) and ***$P$ = 0.0004. n = 4 independent biological replicates per KO clone and matched to WT. **G, H)** Representative confocal images of 2',7'-dichlorodihydrofluorescein (DCF)-based reactive oxygen species measurements in human *PITRM1*-WT and *PITRM1*-KO iPSC-derived microglia. Quantification of DCF fluorescence intensity is shown in H. Mean ± SEM; one-way ANOVA with Bonferroni *post hoc* correction,

*$P$ = 0.0318 and ****$P$ < 0.0001; n = 4 independent experiments. Scale bar, 10 µm. **I)** Quantification of NAD fluorescent analog consumption rates in untreated, CDDO-Me-treated, and *PITRM1*-KO2 iPSC-derived microglia. Mean ± SEM; data are normalized to total protein content; one-way ANOVA with Bonferroni *post hoc* correction; *$P$ = 0.0398; n = 3 independent experiments. **J)** Quantification of p21 and γH2AX immunofluorescence in astrocytes and neurons from *PITRM1*-WT and *PITRM1*-KO2 iPSC lines. Mean ± SEM; unpaired two-tailed *t* test; n = 4 independent experiments.

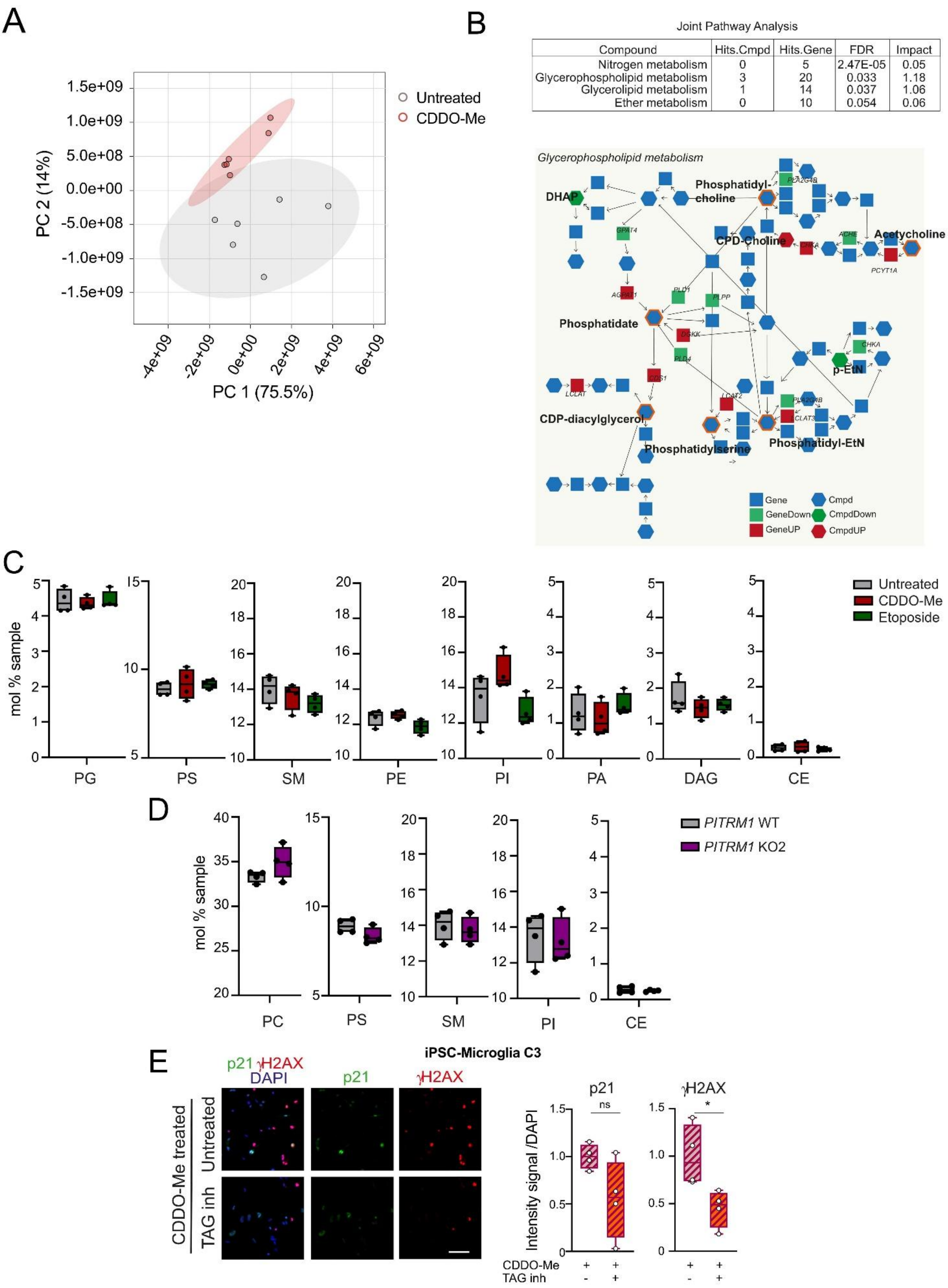


| Compound | Hits.Cmpd | Hits.Gene | FDR | Impact |
|---|---|---|---|---|
| Nitrogen metabolism | 0 | 5 | 2.47E-05 | 0.05 |
| Glycerophospholipid metabolism | 3 | 20 | 0.033 | 1.18 |
| Glycerolipid metabolism | 1 | 14 | 0.037 | 1.06 |
| Ether metabolism | 0 | 10 | 0.054 | 0.06 |

Supplementary Figure 10

**Supplementary Figure 10. Mitochondrial proteotoxic stress disrupts lipid metabolism in human iPSC-derived microglia. A)** Principal component analysis (PCA) score plot showing metabolomic profiles of untreated and 1 μM CDDO-Me-treated human iPSC-derived microglia after 6 hours of treatment. **B)** Upper panel: table of significant pathways derived from the joint pathway analysis of transcriptomic and metabolomic datasets from control and CDDO-Me-treated iPSC-derived microglia (FDR$\leq$0.05). The table lists hit compounds, hit genes, FDR values, and pathway impacts. Lower panel: schematic representation of key dysregulated metabolites and genes in the glycerophospholipid metabolism pathway. Orange borders indicate compounds that were not measured directly but are hypothesized to be affected by alterations in gene expression. **C)** Liquid chromatography-mass spectrometry (LC-MS) quantification of lipid classes expressed as molar percentage (mol%) in iPSC-derived microglia under three conditions: untreated control, CDDO-Me treatment and etoposide treatment. Lipid species include phosphatidylglycerol (PG), phosphatidylserine (PS), sphingomyelin (SM), phosphatidylethanolamine (PE), phosphatidylinositol (PI), phosphatidic acid (PA), diacylglycerol (DAG) and cholesteryl ester (CE). Center line, median; box, 25th–75th percentiles; whiskers, min–max; one-way ANOVA with Bonferroni *post hoc* correction; n=4 independent experiments (C2). **D)** LC-MS analysis of phosphatidylcholine (PC), PS, SM, PI and CE (mol%) in *PITRM1*-WT and *PITRM1*-KO2 iPSC-derived microglia. Center line, median; box, 25th–75th percentiles; whiskers, min–max; unpaired two-tailed *t* test; n=4 independent experiments. **E)** Representative confocal images of p21 (green) and γH2AX (red) in CDDO-Me-treated human iPSC-derived microglia (C3) for 6 hours. Where indicated, cells were co-treated with triacylglycerol (TAG) synthesis inhibitors for 6 hours. Nuclei were stained with DAPI (blue). Scale bar, 10 μm. Right, quantification of fluorescence intensity in DAPI-positive cells. Center line, median; box, 25th–75th percentiles; whiskers, min–max; unpaired two-tailed *t* test, **P* = 0.0152; ns = not significant; n = 4 independent experiments.

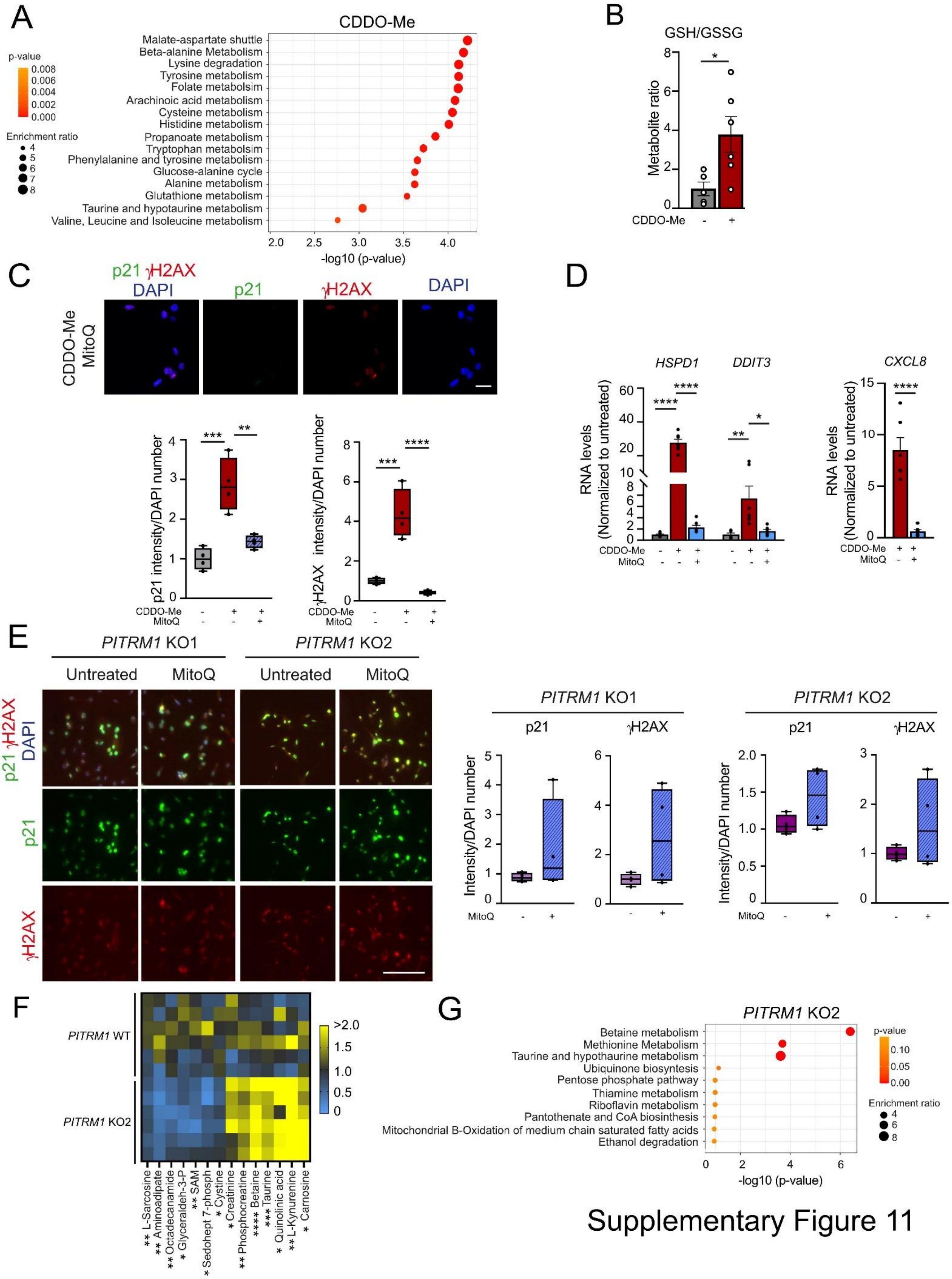


### Supplementary Figure 11

**Supplementary Figure 11. UPR$^{mt}$ induces global metabolic and redox remodeling in senescent microglia. A)** Bubble plot of metabolite set enrichment analysis (MSEA) comparing control and 1 μM CDDO-Me-treated human iPSC-derived microglia. Bubble size represents the enrichment ratio; color scale indicates *P* value. **B)** Relative glutathione redox ratio (GSH/GSSG) in control and CDDO-Me-treated microglia. Mean ± SEM; unpaired two-tailed *t* test, **P* = 0.0275; n = 6 independent experiments (C2). **C)** Representative confocal images of p21 (green) and γH2AX (red) immunofluorescence in human iPSC-derived microglia treated with 1 μM CDDO-Me + 20 nM MitoQ for 6 hours. Nuclei were stained with DAPI (blue). Scale bar, 10 μm. Bottom, quantification of the fluorescence intensity in DAPI-positive cells from control, CDDO-Me, and CDDO-Me + MitoQ groups. Center line, median; box, 25th–75th percentiles; whiskers, min–max; one-way ANOVA with Bonferroni *post hoc* correction, ***P* = 0.0034, ****P* = 0.0005 (p21), 0.0003 (γH2AX), and *****P* < 0.0001; n = 4 independent experiments (C2). **D)** Expression of UPR$^{mt}$ genes *HSPD1* and *DDIT3*, and the SASP cytokine *CXCL8* under cotreatment conditions. Mean ± SEM; data are normalized to the untreated control in each condition; one-way ANOVA with Bonferroni *post hoc* correction or unpaired two-tailed *t* test, **P* = 0.0133, ***P* = 0.0060, and *****P* <0.0001; n = 7 independent experiments (C2). **E)** Representative confocal images of p21 (green) and γH2AX (red) immunofluorescence in human *PITRM1*-KO (KO1 and KO2) iPSC-derived microglia. *PITRM1*-KO cells were either untreated or treated with 20 nM MitoQ for 6 hours. Nuclei were stained with DAPI (blue). Scale bar, 10 μm. Right, quantification of fluorescence intensity in DAPI-positive cells across conditions. Center line, median; box, 25th–75th percentiles; whiskers, min–max; unpaired two-tailed *t* test, n = 4 independent experiments. **F)** Heatmap of the 14 significantly altered metabolites in *PITRM1*-KO2 versus *PITRM1*-WT microglia. Metabolites are ordered by relative abundance. Yellow indicates increased abundance; blue indicates decreased abundance. Data are normalized to WT controls; unpaired two-tailed *t* test. **P* = 0.0155 (glyceraldehyde-3-phosphate), 0.0477 (sedoheptulose-7-phosphate), 0.0319 (cystine), 0.0495 (creatinine), 0.0172 (quinolinic acid), 0.0148 (carnosine); ***P* = 0.0041 (L-sarcosine), 0.0082 (aminoadipate), 0.0040 (octadecanamide), 0.0068 (S-adenosyl-L-methionine), 0.0029 (phosphocreatine), 0.0037 (L-kynurenine); ****P* = 0.0002 (taurine); *****P* <0.0001; n = 6 independent experiments. **G)** Bubble plot of MSEA comparing *PITRM1*-KO2 and *PITRM1*-WT microglia. Bubble size indicates enrichment ratio; color denotes *P* value.

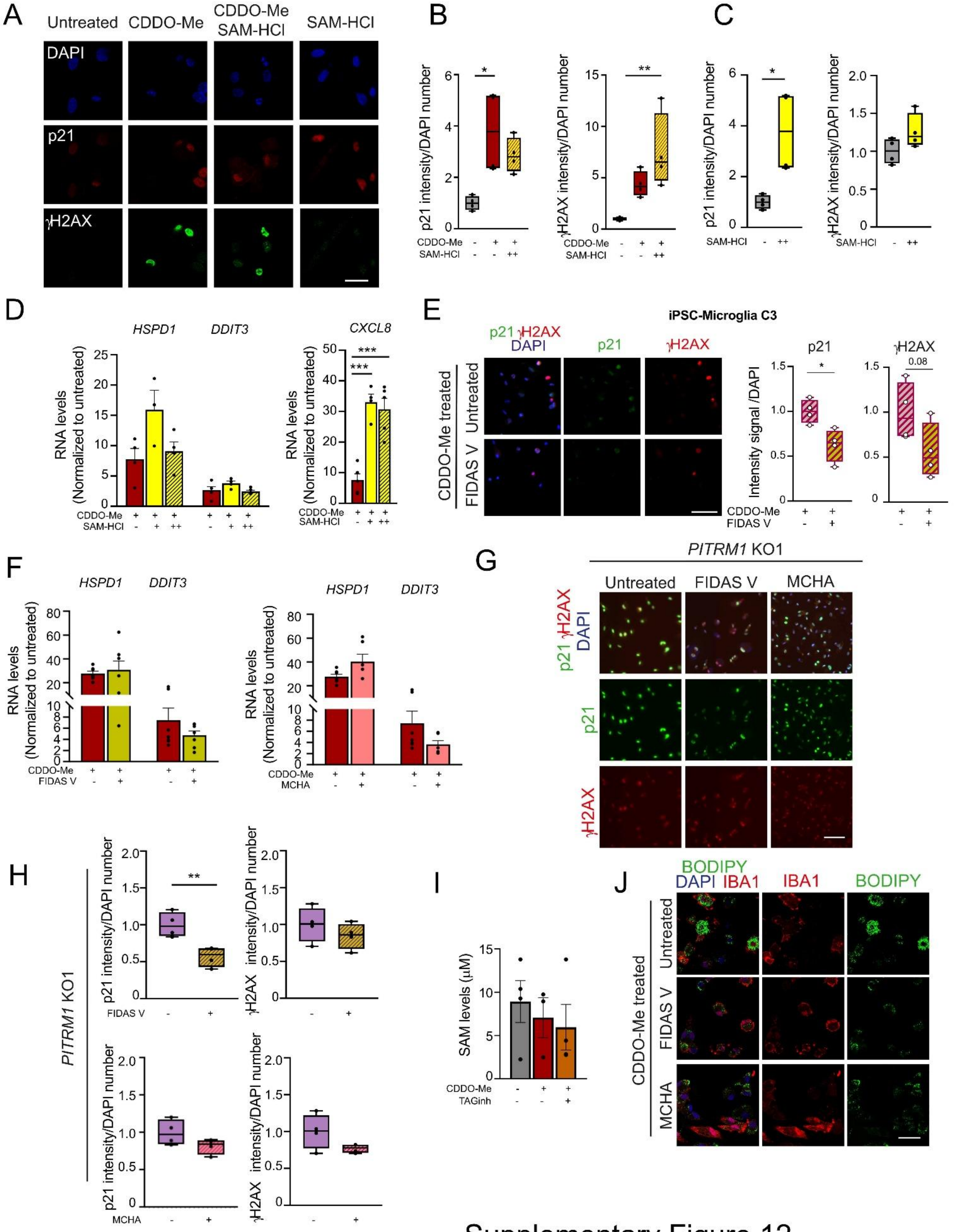


Supplementary Figure 12

**Supplementary Figure 12. The UPR$^{mt}$-driven senescent phenotype in microglia is linked to changes in methionine metabolism. A)** Representative confocal images of p21 (red) and γH2AX (green) immunofluorescence in human iPSC-derived microglia (C2). Cells were untreated or treated for 6 hours with 0.5 mM S-adenosylmethionine hydrochloride (SAM-HCl), 1 µM CDDO-Me, or the combination. Nuclei were counterstained with DAPI (blue). Scale bar, 10 µm. **B, C)** Quantification of fluorescence intensity in DAPI-positive cells comparing control versus 1 µM CDDO-Me alone and CDDO-Me + 0.5 mM SAM-HCl (B), or control versus 0.5 mM SAM-HCl alone (C). Center line, median; box, 25th–75th percentiles; whiskers, min–max; one-way ANOVA with Bonferroni *post hoc* correction (B) or unpaired two-tailed *t* test (C), **P* = 0.0113 (CDDO-Me versus control), 0.0139 (SAM-HCl versus control); ***P* = 0.0071; n = 4 independent experiments (C2). **D)** mRNA expression of UPR$^{mt}$ genes *HSPD1* and *DDIT3*, and the SASP cytokine gene *CXCL8*, following treatment with CDDO-Me alone or in combination with 0.1 mM (+) or 0.5 mM (++) SAM-HCl. Data are normalized to untreated controls Mean ± SEM; one-way ANOVA with Bonferroni *post hoc* correction, ****P* = 0.0002 (CDDO-Me versus CDDO-Me + 0.1 mM SAM-HCl), 0.0003 (CDDO-Me versus CDDO-Me + 0.5 mM SAM-HCl); n = 4 (*HSPD1*), n = 5 (*DDIT3* and *CXCL8*) independent experiments (C2). **E)** Representative confocal images of p21 (green) and γH2AX (red) immunofluorescence in CDDO-Me-treated human iPSC-derived microglia (C3). Where indicated, cells were treated with 25 µM FIDAS V for 6 h. Nuclei were stained with DAPI (blue). Scale bar, 10 µm. Right, quantification of fluorescence intensity in DAPI-positive cells. Center line, median; box, 25th–75th percentiles; whiskers, min–max, unpaired two-tailed *t* test, **P* = 0.0268; n = 4 independent experiments. **F)** mRNA expression of UPR$^{mt}$ genes *HSPD1* and *DDIT3* following treatment with 1 µM CDDO-Me alone or in combination with 25 µM FIDAS V or 250 µM MCHA. Data are normalized to untreated controls. Mean ± SEM; unpaired two-tailed *t* test; n = 7 independent experiments (C2). **G)** Representative confocal images of p21 (green) and γH2AX (red) immunofluorescence in human *PITRM1*-KO1 iPSC-derived microglia. Cells were untreated or treated with 25 µM FIDAS V or 250 µM MCHA for 6 hours. Nuclei were stained with DAPI (blue). Scale bar, 10 µm. **H)** Quantification of fluorescence intensity in DAPI-positive *PITRM1*-

KO1, *PITRM1*-KO1 + FIDAS V and *PITRM1*-K1 + MCHA cells. Center line, median; box, 25th–75th percentiles; whiskers, min–max; unpaired two-tailed *t* test, ***P* = 0.0071; n = 4 independent experiments. **I)** Intracellular SAM levels measured using the Bridge-It® fluorescence assay in untreated control cells, CDDO-Me-treated cells and cells cotreated with CDDO-Me and a triacylglycerol (TAG) synthesis inhibitor. Mean ± SEM; one-way ANOVA with Bonferroni *post hoc* correction; n = 4 independent experiments (C2). **J)** Representative confocal images of IBA1 (red), BODIPY (green), and DAPI (blue) immunofluorescence in human iPSC-derived microglia (C2) treated with CDDO-Me with or without 25 µM FIDAS V or 250 µM MCHA for 6 hours. Scale bars, 10 µm. Images are representative of at least two independent experiments.

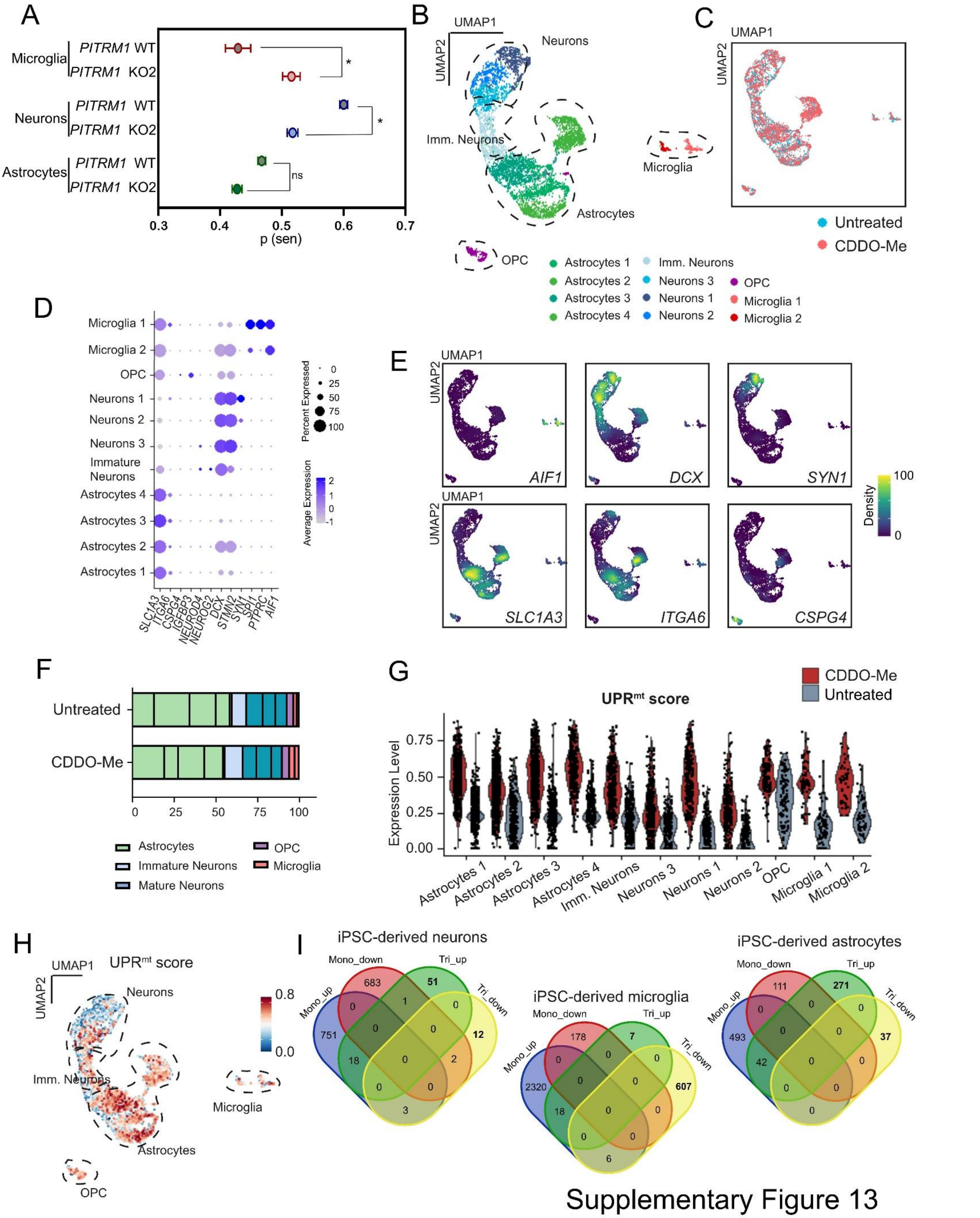


Supplementary Figure 13

**Supplementary Figure 13. Single-cell transcriptomic analysis reveals cell type-specific responses in control and CDDO-Me-treated human iPSC-derived tricultures. A)** Predicted senescence scores for *PITRM1*-WT and *PITRM1*-KO2 human iPSC-derived microglia, neurons, and astrocytes within the triculture system. Mean ± SEM; unpaired two-tailed *t* test versus control within each cell type, **P* = 0.0304 (microglia), 0.0440 (neurons); ns= not significant; n = 3 independent experiments. **B)** UMAP cluster integration plot showing unsupervised clustering of control and CDDO-Me-treated human iPSC-derived tricultures. **C)** UMAP dimensionality reduction plot of integrated datasets from control and CDDO-Me-treated tricultures. **D)** Dot plot showing expression of selected cell type-specific marker genes in CDDO-Me-treated versus control tricultures. **E)** Cell density visualization using Nebulosa kernel-based estimation to display UMAP density features for cells expressing *AIF1, DCX, SYN1, SLC1A3, ITGA6,* or *CSPG4*. **F)** Cluster frequency distribution across datasets showing the relative abundance of each cell cluster in untreated and CDDO-Me-treated tricultures. **G)** Violin plots of $UPR^{mt}$ signature scores in untreated and CDDO-Me-treated tricultures using the UCell algorithm. **H)** UMAP projection of $UPR^{mt}$ UCell scores at single-cell resolution visualized as cellular density features. **I)** Venn diagrams comparing differentially expressed genes in iPSC-derived neurons, astrocytes and microglia between monoculture and triculture systems. Blue, genes upregulated in monoculture; red, genes downregulated in monoculture; green, genes upregulated in triculture; yellow, genes downregulated in triculture.

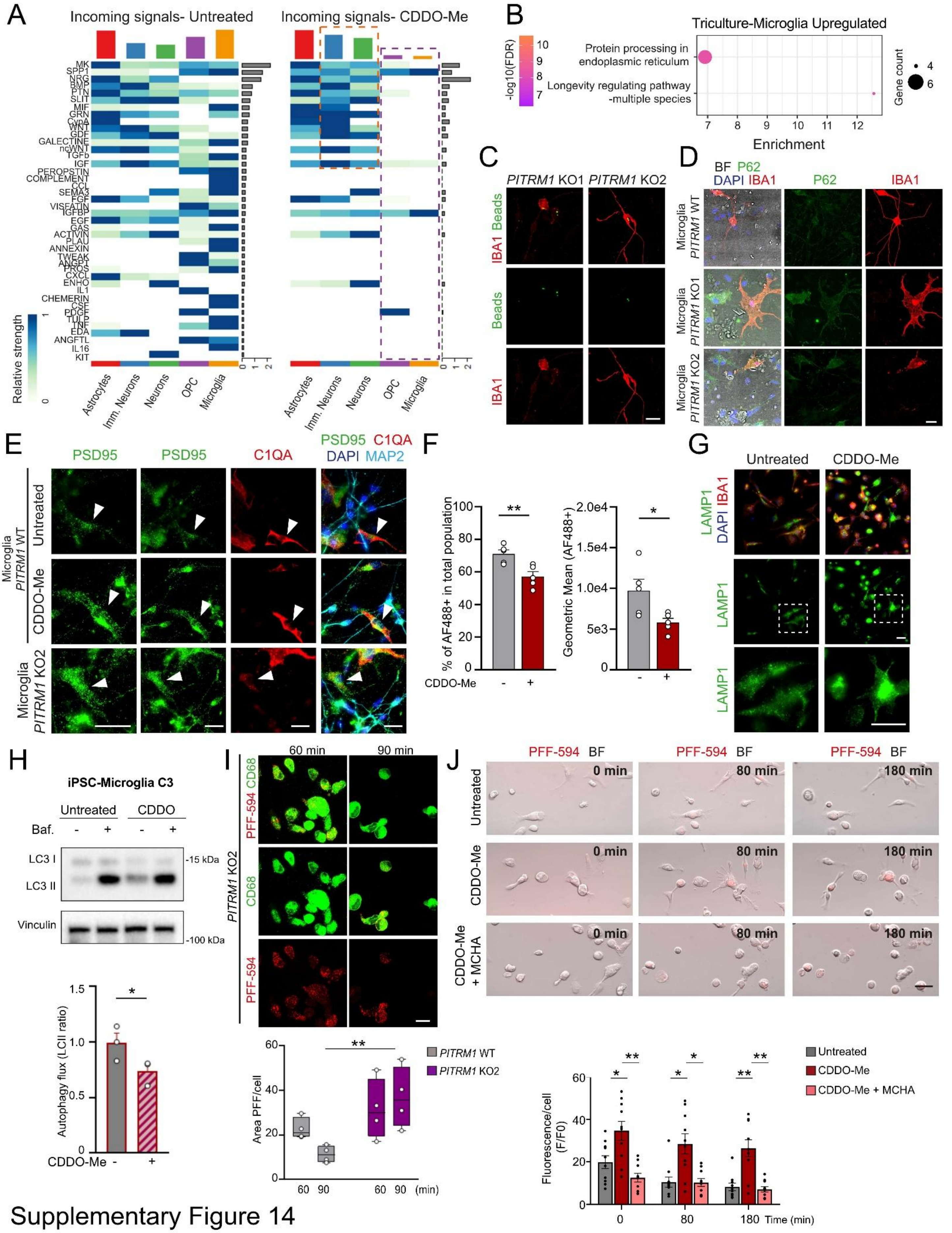


Supplementary Figure 14

**Supplementary Figure 14. Mitochondrial proteotoxic stress induces microglial senescence and alters intercellular communication in human iPSC-derived tricultures. A)** Heatmap showing incoming ligand-receptor signaling interactions across cell types within tricultures under control and CDDO-Me-treated conditions. The y axis denotes ligand-receptor pairs, and the top bar plot shows summed incoming signal strength per cell type. Color intensity reflects signaling strength. Dashed orange box highlights increased incoming signaling in neurons and astrocytes following treatment; dashed purple box highlights reduced incoming signaling in microglia and oligodendrocytes. **B)** Bubble plot of upregulated Reactome pathways in the microglial cluster within the triculture. Bubble size represents gene count; color scale indicates -log10(FDR). **C)** Representative fluorescence images of Alexa Fluor 488-labeled beads (green) and IBA1 (red) immunostaining in *PITRM1*-WT iPSC-derived tricultures containing *PITRM1*-KO1 or *PITRM1*-KO2 microglia. Scale bar, 10 μm. Images are representative of at least two independent experiments. **D)** Immunostaining for P62 (green) and IBA1 (red) in *PITRM1*-WT iPSC-derived tricultures containing *PITRM1*-WT or *PITRM1*-KO microglia. Nuclei were stained with DAPI (blue). BF, bright field. Scale bars, 10 μm. Images are representative of at least two independent experiments. **E)** Representative images of *PITRM1*-WT tricultures containing WT or *PITRM1*-KO2 microglia under untreated or CDDO-Me-treated conditions. Cultures were stained for PSD95 (green), C1QA (red), MAP2 (cyan), and DAPI (blue). Arrows indicate microglia selected for magnification. Scale bar, 10 μm. Images are representative of at least two independent experiments. **F)** Phagocytic activity assay in iPSC-derived microglia (C2) exposed to CDDO-Me or left untreated, followed by incubation with AF488-labeled beads and analysis by flow cytometry. Left, percentage of $AF488^+$ cell population from total population; right, geometric mean fluorescence intensity of $AF488^+$ population reflecting particle uptake per cell. Mean + SEM; unpaired two-tailed *t* test, **P* = 0.0303, ***P* = 0.0077; n = 5 independent experiments. **G)** Representative confocal images of IBA1 (red) and the lysosomal marker LAMP1 (green) in untreated and CDDO-Me-treated iPSC-derived microglia (C2). Nuclei were stained with DAPI (blue). Scale bar, 10 μm. Images are representative of at least two independent experiments. **H)** Western blot analysis of LC3-

II in untreated and CDDO-Me-treated iPSC-derived microglia, with or without bafilomycin A1 (Baf) for 4 hours. Autophagic flux was quantified as the ratio of LC3-II levels in Baf-treated versus untreated cells and then normalized to the untreated control. Mean + SEM; unpaired two-tailed *t* test, **P* = 0.0192, n = 3 independent experiments (C3). **I)** Representative confocal images of CD68 (green) and PFF-594 (red) in *PITRM1*-WT and *PITRM1*-KO2 human iPSC-derived microglia (C2), captured at 60- and 90-min post-treatment. Scale bar, 10 μm. Center line, median; box, 25th–75th percentiles; whiskers, min–max; two-way ANOVA with Bonferroni *post hoc* correction between genotypes at each time point, ***P* = 0.0077. n = 4 independent experiments. **J)** Live-cell imaging of PFF-594 (red) and bright-field (BF) images in human iPSC-derived microglia under untreated, CDDO-Me-treated, or CDDO-Me + MCHA conditions at different timepoints. Scale bar, 10 μm. Bottom, quantification of the PFF-594-positive area relative to total cell area over time. Mean ± SEM; repeated measures two-way ANOVA (time × treatment) with Geisser-Greenhouse correction and Bonferroni *post hoc* correction between treatments at each time point; **P* = 0.0436 (0 mins, untreated versus CDDO-Me), 0.0137 (80 mins, untreated versus CDDO-Me), 0.0114 (80 mins, CDDO-Me versus CDDO-Me + MCHA), ***P* = 0.0019 (0 mins, CDDO-Me versus CDDO-Me + MCHA), 0.0038 (180 mins, untreated versus CDDO-Me), 0.0024 (180 mins, CDDO-Me versus CDDO-Me + MCHA); n = 10 cells from 2 independent experiments (C2).

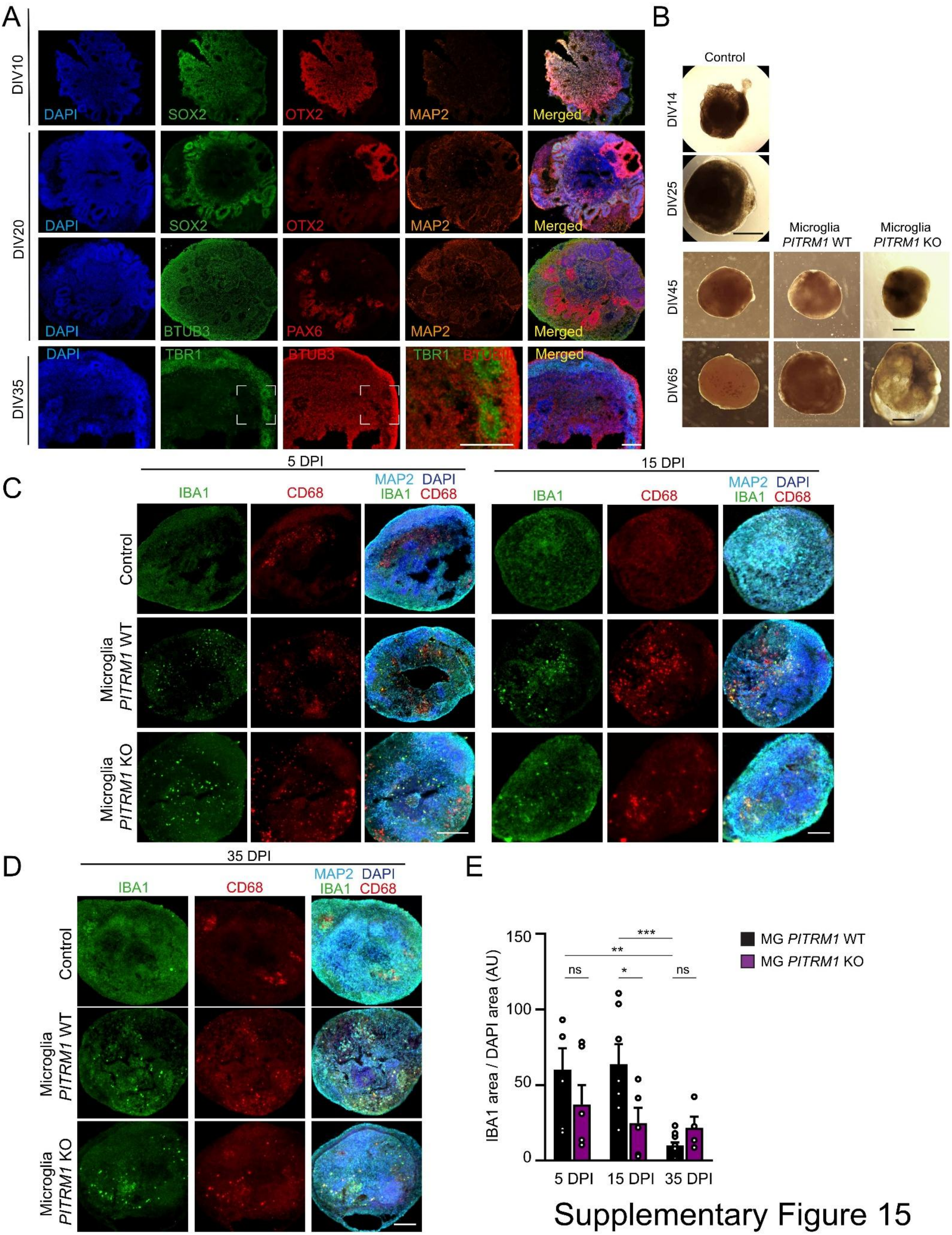


Supplementary Figure 15

**Supplementary Figure 15. Development and characterization of iPSC-derived cortical organoids and MgBr assembloids.** All experiments were performed using iPSC line C2 and KO2. **A)** Immunostaining for the indicated markers in cortical organoids at 10, 20, and 35 days in vitro (DIV). Nuclei were counterstained with DAPI (blue). Scale bars, 25 μm. **B)** Representative brightfield images of iPSC-derived cortical organoids and MgBr assembloids, generated from *PITRM1*-WT human iPSC-derived organoids with the integration of *PITRM1*-WT or *PITRM1*-KO human iPSC-derived microglia. Scale bars, 1 mm. Images are representative of at least two independent differentiations. **C, D)** Representative immunofluorescence images of IBA1 (green), CD68 (red), and MAP2 (cyan) staining in MgBr assembloids at 5 days post integration (DPI, C), 15 DPI (C), and 35 DPI (D). Cell nuclei were counterstained with DAPI (blue). Scale bar, 100 μm. **E)** Quantification of IBA1-positive area normalized to DAPI-positive area in MgBr assembloids at 5, 15, and 35 DPI. Mean ± SEM; two-way ANOVA with Bonferroni *post hoc* correction, **P* = 0.0149, ***P* = 0.0021, ****P* = 0.0006, ns = not significant; n = 6 (5 DPI WT), 7 (15 DPI WT), 10 (35 DPI WT) and 6 (5 DPI KO), 5 (15 DPI KO), 4 (35 DPI KO) individual organoids.

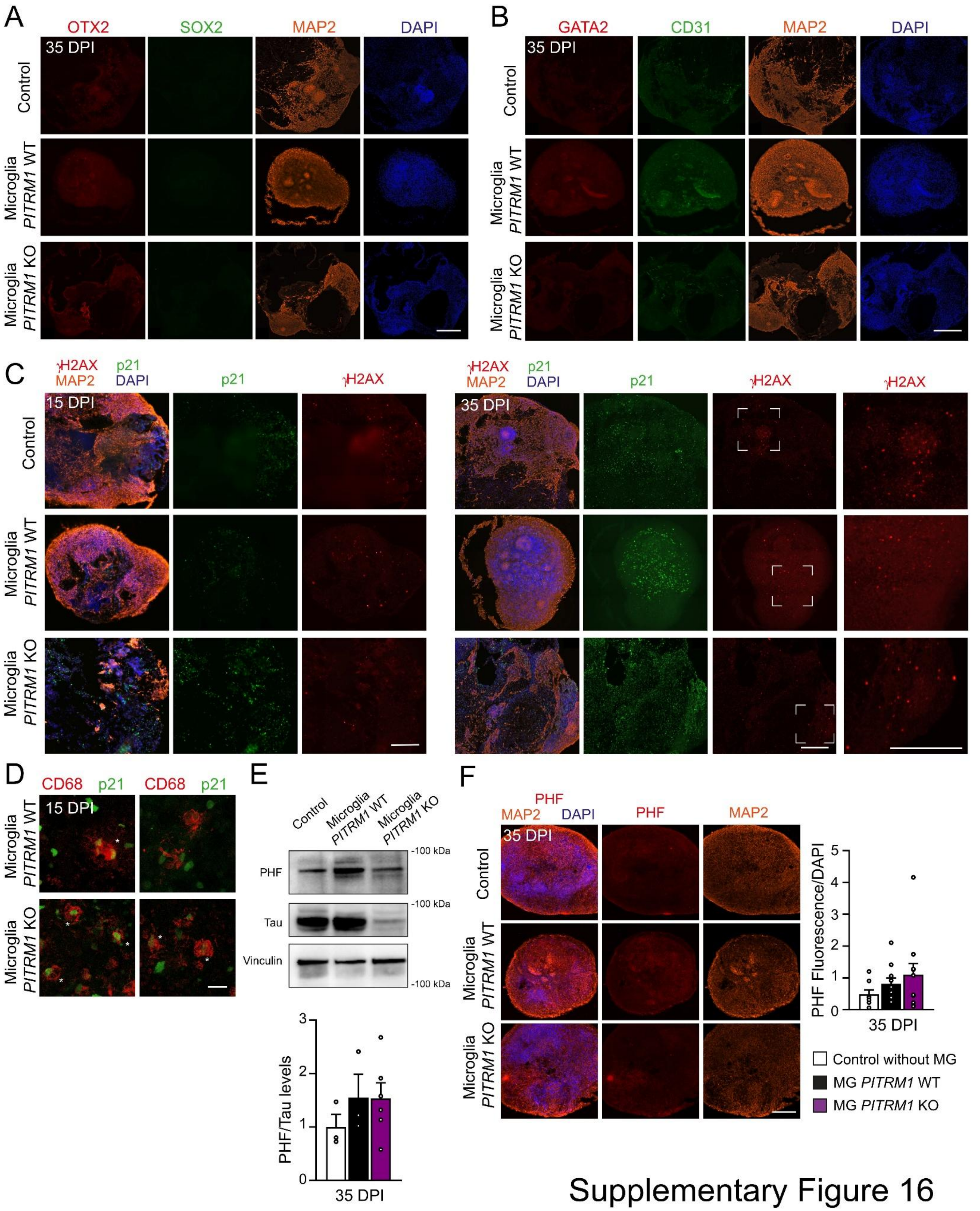


Supplementary Figure 16

**Supplementary Figure 16. Integration of *PITRM1*-KO microglia induces a senescence phenotype in iPSC-derived MgBr assembloids.** All experiments were performed using iPSC line C2 and KO2. **A)** Immunostaining for SOX2 (green), OTX2 (red), and MAP2 (orange) in MgBr assembloids at 35 days post integration (DPI). Cell nuclei were counterstained with DAPI (blue). Scale bars, 25 µm. **B)** Immunostaining for CD31 (green), GATA2 (red), and MAP2 (orange) in MgBr assembloids at 35 DPI. Nuclei were counterstained with DAPI (blue). Scale bars, 70 µm. Images are representative of at least four independent organoids. **C)** Immunostaining for p21 (green), γH2AX (red), and MAP2 (orange) in MgBr assembloids at 15 and 35 DPI. Cell nuclei were counterstained with DAPI (blue). Right, high-magnification images of the insets. Scale bars, 100 µm. **D)** Representative confocal images of p21 (green) and CD68 (red) staining in MgBr assembloids at 15 DPI. White asterisks indicate p21-positive microglia. Scale bar, 10 µm. **E)** Representative Western blot showing phosphorylated tau in the Thr181 residue (PHF), total tau and vinculin (loading control) protein levels in MgBr assembloids at 35 DPI. Quantification of PHF/total tau ratios normalized to vinculin and the control group is shown below. Mean ± SEM; one-way ANOVA with Bonferroni *post hoc* correction; n = 3 (control), 3 (microglia WT) and 6 (microglia KO) individual organoids. **F)** Representative immunofluorescence images of phosphorylated tau (PHF, red) and MAP2 (orange) in MgBr assembloids at 35 DPI. Cell nuclei were counterstained with DAPI (blue). Right, quantification of PHF fluorescence intensity normalized to DAPI-positive area. Mean ± SEM; one-way ANOVA with Bonferroni *post hoc* correction; n = 8 (control), 10 (microglia WT) and 11 (microglia KO) individual organoids.

**Supplemental Methods**

**Generation of human iPSC-derived cortical neurons, astrocytes, and microglia**

The control neuronal precursor cell (NPC) lines were previously generated and characterized [1, 2]. iPSC colonies were selected and cultured for four days in embryoid body (EB) media containing 20% KO serum replacement (Fisher Scientific, Cat. # 0-828-028) and 80% DMEM/F-12 (Thermo Scientific, Cat. # 11320074) supplemented with 1% nonessential amino acids (NEAAs) (Thermo Scientific, Cat. # 11140035), 1% penicillin-streptomycin (PS, Thermo Scientific, Cat. # 15140122), 10 µM SB431542 (SB, Selleck Chemicals, Cat. # S1067), and 2.5 µM dorsomorphin (Sigma-Aldrich, Cat. # P5499). Beginning on day five, the EBs were cultured for an additional 4 days in N2B27 medium containing 100% DMEM/F-12, 1% N2 (Fisher Scientific, Cat. # 17502048), 1% B27 without vitamin A (Thermo Scientific, Cat. # 12587010), 1% NEAAs, 1% PS, 10 µM SB, 2.5 µM dorsomorphin, and 20 ng/mL FGF2 (PeproTech, Cat. # 100-18B-250). Then, neural rosettes were isolated with Accutase (STEMCELL Technologies, Cat. # 07920), replated on a surface coated with Corning® Matrigel® Growth Factor Reduced Basement Membrane Matrix (Matrigel, Dutscher, Cat. # 354230) and maintained in N2B27 medium. Secondary or tertiary rosettes were manually dissected to purify NPCs, which were maintained in basal NPC medium containing 1:1 DMEM-Ham's F-12 and Neurobasal medium (Fisher Scientific, Cat. # 21103049), 0.5% N2, 1% B27 without vitamin A, 1% PS, and 1% GlutaMAX (Fisher Scientific, Cat. # 35050038), supplemented with 3 µM CHIR99021 (Sigma-Aldrich, Cat. # SML1046-5MG), 0.5 µM purmorphamine (Merck, Cat. # 540220), and 150 µM ascorbic acid (AA, Sigma-Aldrich, Cat. # A8960-5G). The NPCs were split into Matrigel-coated wells every 5–6 days at a ratio of 1:10. For cortical neuron differentiation, 80% confluent NPCs were seeded in neuronal differentiation medium and cultured for one week. This medium consisted of basal neuronal medium (1:1 DMEM-Ham's F-12 medium and Neurobasal medium), 0.5% N2, 1% B27 without vitamin A, 1% PS, and 1% GlutaMAX supplemented with 200 µM AA, 1 µM purmorphamine, and 100 ng/mL FGF8 (PeproTech, Cat. # 100-25-500). After 7 days, the cells were split and seeded in neuronal maturation medium consisting of basal neuronal medium supplemented with 20 ng/mL BDNF (PeproTech, Cat. # 450-02) and 1 mM dibutyryl-cyclic AMP (dbcAMP; Sigma-Aldrich, Cat. # A0455,1000-ITW). The medium was replaced every other day, with the final split performed after 14 days in vitro (DIV). Experiments were conducted after 21 DIV. For astrocyte differentiation, NPCs were seeded at a density of 15,000 cells/cm² in commercial astrocyte differentiation medium (ScienCell Research Laboratories, Cat. # 1801-SC)[3]. The cells were passaged (15,000 cells/cm² in differentiation medium) with Accutase every 5–6 days on Matrigel-coated plates. After 30 days, the medium was replaced with astrocyte maturation medium [3]. The cells were split at 90% confluence, maintained in maturation medium, and used for experiments between passages 3 and 10 after the medium was changed to maturation medium.

A total of $3\times10^6$ iPSCs were seeded into a well of an AggreWell 800 plate (STEMCELL Technologies, Cat. # 34811) to form EBs and generate microglia [4, 5]. EBs were cultured for 4 days in mTeSR™ Plus medium (STEMCELL Technologies, Cat. # 100-0276) supplemented daily with 50 ng/mL bone morphogenetic protein 4 (BMP4) (Proteintech, Cat. # HZ-1045), 50 ng/mL vascular endothelial growth factor (VEGF) (ImmunoTools, Cat. # 11343665), and 20 ng/mL stem cell factor (SCF) (Proteintech, Cat. # HZ-0024). After 4 days, the EBs were transferred to 6-well plates (15 EBs/well) and cultured in X-VIVO15 medium (Lonza, Cat. # 02-060Q) supplemented with 1% GlutaMAX, 1% PS, 100 ng/mL macrophage colony-stimulating factor (M-CSF, Proteintech, Cat. # HZ-1192), 25 ng/mL interleukin-3 (IL-3, Proteintech, Cat. # HZ-1074), and 0.055 mM β-mercaptoethanol (Sigma-Aldrich, Cat. # M3148). Fresh medium was added weekly. After 3–4 weeks, the macrophage precursors that appeared in the supernatant were harvested. These cells were then plated at a density of 100,000 cells/cm² and cultured for 10 days in Advanced DMEM/F12 medium (Thermo Fisher, Cat. # 12634010) supplemented with 1% N2, 1% GlutaMAX, 1% PS, 100 ng/mL M-CSF, 100 ng/mL IL-34 (Proteintech, Cat. # Hz-1316), 10 ng/mL granulocyte-macrophage colony-stimulating factor (GM-CSF, Proteintech, Cat. # HZ-1002), and 0.055 mM β-mercaptoethanol.

**Human iPSC-derived tricultures**

Prior to establishment of the triculture system, iPSC-derived cortical neurons were generated as described above until the final split at 14 DIV. For this purpose, neurons were dissociated with Accutase and plated on glass coverslips coated with Matrigel. Neurons were maintained in neuronal maturation medium composed of basal neuronal medium supplemented with 20 ng/mL BDNF and 1 mM dbcAMP. After four days, iPSC-derived astrocytes between passage 3 and passage 10 after maturation were seeded onto the neurons at a ratio of 3:1 (neurons/astrocytes) and maintained in the final neuronal maturation medium. Three days later, iPSC-derived microglial precursors were added to the cocultures at a ratio of 3:1:1 (neurons/astrocytes/microglia). The tricultures were maintained in the final neuronal maturation medium supplemented with 100 ng/mL IL-34, 100 ng/mL M-CSF, and 10 ng/mL GM-CSF for an additional 7 days before the experiments were performed.

**Human iPSC-derived MgBr assembloids**

We used a previously described protocol to generate and culture human cortical organoids, with modifications [6]. A total of 9,000 iPSCs were seeded into a well of a Nunclon™ Sphera™ 96-well microplate (Thermo Fisher, Cat. # 174925) to generate EBs. The EBs were cultured from days 0 to 5 in dual SMAD inhibition medium to promote neuroectoderm patterning. This medium consisted of mTeSR™ Plus supplemented daily with 1% NEAAs, 1% GlutaMAX, 100 μM β-mercaptoethanol, 200 nM LDN 193189 hydrochloride (LDN, Axon MedChem, Cat. # 1509), 10 μM SB, and 50 μM Y-27632 (ROCK inhibitor; Selleck Chemicals, Cat. # S1049). Beginning on day 6, the organoids were cultured in cortical induction medium consisting of 1:1 DMEM-Ham's F-12 and Neurobasal medium supplemented with 1% NEAAs, 1% GlutaMAX, 1% N2, 1% B27 without vitamin A, 100 μM β-mercaptoethanol, 20 ng/mL FGF2, and 20 ng/mL EGF (PeproTech, Cat. # HZ-1326). On day 7, the EBs were embedded in Matrigel at a 3:2 ratio with medium to form an embedding mixture. The EBs were washed with fresh medium and embedded in Matrigel in a 6-well ultralow-attachment plate with a mixture of Matrigel and medium. The Matrigel-EB mixture was incubated for 30 minutes at 37°C, and the organoids were maintained in cortical induction medium until day 14. From days 14 to 30, the organoids were maintained in differentiation medium consisting of 100% Neurobasal medium supplemented with 1% NEAAs, 1% GlutaMAX, 1% N2, 1% B27 without vitamin A, 1% PS, 100 µM β-mercaptoethanol, 2.5 µg/mL insulin (Thermo Scientific, Cat. # 12585014), 200 µM AA, 20 ng/mL BDNF, and 1 mM dbcAMP. On day 20, the Matrigel was dissociated from the organoids, and the plates were placed on an orbital shaker to increase oxygenation and improve medium distribution. Individual brain organoids on day 30 of differentiation were placed into a well of a Nunclon™ Sphera™ 96-well microplate to generate iPSC-derived MgBr assembloids. Subsequently, on day 10 of in vitro differentiation, 2x10^5 pre-differentiated iPSC-derived microglia were seeded on top of the organoids in a final volume of 200 µL of assembloid maturation medium. This medium consisted of 100% Neurobasal medium supplemented with 1% NEAAs, 1% GlutaMAX, 1% N2, 1% B27, 1% PS, 100 µM β-mercaptoethanol, 20 ng/mL BDNF, 1 mM dcAMP, 10 ng/mL GDNF (Fisher Scientific, Cat. # 450-10), 100 ng/mL IL-34, 100 ng/mL M-CSF, and 10 ng/mL GM-CSF. The medium was changed twice daily for 3 days without removing the microglia in suspension. The assembloids were then transferred to a 6-well ultralow-attachment plate. After 2 days, the MgBr assembloids were placed on an orbital shaker and maintained in maturation medium until day 45 (15 DPI) or day 65 (35 DPI) of differentiation.

**Immunofluorescence staining and image analysis**

For immunostaining, cells seeded on coverslips were fixed with 4% paraformaldehyde (PFA; Sigma-Aldrich, Cat. # 1004968350) for 10 minutes, rinsed with phosphate-buffered saline (PBS, Thermo Scientific, Cat. # 14190169) and blocked with 10% normal goat or donkey serum (NGS/NDS, Millipore, Cat. # 50095; Sigma-Aldrich, Cat. # S30-100ML) in PBS containing 0.2% Triton X-100 (PBST) at room temperature for 1 hour. The cells were then incubated with primary antibodies diluted in PBST containing 5% NGS/NDS overnight at 4°C. The following day, the cells were incubated for 2 hours at room temperature with Alexa 488/568/647-conjugated secondary antibodies (1:1000, Invitrogen). Nuclei were stained for 5 minutes with 1 μM 4’,6-diamidino-2-phenylindole dihydrochloride (DAPI; Sigma-Aldrich, Cat. # D9542-10MG), coverslips were mounted on glass slides with mounting medium (Dako, Agilent Technologies, Cat. # S302380-2), and images were acquired with a Leica TCS SP8 confocal microscope with a 60×/1.4 numerical aperture oil immersion objective (Leica). Brain organoids and MgBr assembloids were fixed with 4% (w/v) PFA for 1 hour, and individual organoids were equilibrated in 30% sucrose in PBS overnight at 4°C. The next day, the organoids were embedded in blocks in OCT compound (Tissue-Tek, Cat. # 0094-4583-01). Slices with a thickness of 20 μm were cryosectioned and mounted on Epredia™ SuperFrost™ Plus slides (Fisher Scientific, Cat. # 10149870). The tissues were blocked with 10% (v/v) PBS containing NGS in 0.5% Triton X-100, incubated with primary antibodies overnight, and then incubated with secondary antibodies for 3 hours. When needed, BODIPY staining was performed to visualize lipid droplets within cells. The cells or tissues were incubated with 1 μM BODIPY 493/503 dye (Thermo Fisher, Cat. # D3922) diluted in PBS for 30 minutes at room temperature. For the detection of protein aggregates, the sections were stained with 10 μM thioflavin-T (Sigma-Aldrich, Cat. # T3516) for 30 minutes at room temperature and washed three times with PBS. The sections were stained with 1 μM DAPI for 5 minutes, and after three washes with PBS, they were mounted with Dako mounting medium for image acquisition. For the mouse brain tissue, antigen retrieval was performed in free-floating brain slices for 20 minutes in citrate buffer at 80°C. The tissues were blocked with mouse-on-mouse blocking reagent (MOM; Vector Laboratories, Cat. # BMK-2202) according to the manufacturer’s instructions. Primary antibodies were incubated with the MOM protein concentrate overnight, followed by an incubation with secondary antibodies for 3 hours. Images were acquired with a Leica TCS SP8 confocal microscope with a 60×/1.4 numerical aperture oil immersion objective (Leica) or with an EVOS M7000 Imaging System microscope with a 20×/0.4 numerical aperture (Thermo Fisher). For the quantification of p21-, p16-, and γH2AX-positive nuclei, images were analyzed via the counting and scoring pipeline in CellProfiler image analysis software (version 4.2.6, RRID: SCR_007358) according to the suggested guidelines, with DAPI as a counterstain. For the quantification of the total area and fluorescence, the fluorescence intensity per area was calculated with Fiji (version 2.7.0, RRID:SCR_002285). For mouse brain tissue, colocalization between p21 and IBA1 was quantified using the Coloc2 plugin in Fiji and reported as Manders' tM1 and tM2 values, reflecting the fractional overlap of the p21 signal within IBA1+ cells. A list of the antibodies used is provided in Supplementary Table 1.

**Functional assays via live-cell imaging**

The cells were incubated with specific fluorescent dyes tailored for each desired measurement to assess cellular functions by live-cell imaging. All the incubations were performed for 30 minutes at 37°C. After the incubation, the cells were washed twice with PBS, and imaging was conducted with a Leica TCS SP8 confocal microscope equipped with a 63×/1.4 numerical aperture oil immersion objective (Leica). For the assessment of mitochondrial function, 100 nM tetramethylrhodamine methyl ester perchlorate (TMRM) combined with 100 nM MitoTracker Green (both from Invitrogen, Cat. # T668, and # M46750) were used. An excitation wavelength of 568 nm was used for TMRM, and an excitation wavelength of 488 nm was used for MitoTracker Green. Emitted fluorescence was measured at wavelengths shorter than 574 nm for TMRM and between 490 and 530 nm for MitoTracker Green. The TMRM intensity was quantified in mitochondria stained with MitoTracker Green via Fiji (version 2.7.0, RRID: SCR_002285). The basal TMRM fluorescence intensity was set at 100%, and the fluorescence intensity after complete mitochondrial depolarization by carbonyl cyanide-(trifluoromethoxy) phenylhydrazone (FCCP, Sigma-Aldrich, Cat. # C2910) was considered the background fluorescence intensity. The background value was then subtracted from the basal value to calculate a corrected intensity measurement. For calcium measurements, iPSC-derived neurons and astrocytes were incubated with 1 μM Fluo-4 AM (Invitrogen, Cat. # F14217). An excitation wavelength of 488 nm and an emission wavelength range of 490–530 nm were used for Fluo-4 AM. For characterization experiments, neurons were stimulated with 0.6 mM KCl (Sigma-Aldrich, Cat. # P3911-25G), and astrocytes were stimulated with 1 mM glutamate (Merck, Cat. # 6106-04-3) for 3 minutes following an equilibration period to establish the baseline fluorescence intensity. For neuronal function assays, 100 mM KCl was applied following 5 min equilibration, and fluorescence was recorded every 30 s for 15 min. Oxidative stress was evaluated using 1 μM DCF (Invitrogen, Cat. # C400). An excitation wavelength of 488 nm and an emission wavelength range of 490–530 nm was used for DCF. Images were analyzed by measuring the fluorescence intensity with Fiji (version 2.7.0, RRID:SCR_002285), and the values were normalized to the baseline values to ensure consistency.

**Bulk RNA-sequencing and data analysis**

Total RNA was isolated from three biological replicates per sample of iPSC-derived neurons (24 hours), astrocytes, and microglia (6 hours) treated with or without 1 μM CDDO-Me using an RNeasy Mini Kit (Qiagen, Cat. # 74106) according to the manufacturer's instructions. High-quality RNA samples were subjected to bulk RNA-sequencing at GENEWIZ GmbH (Leipzig, Germany). Briefly, mRNAs were enriched via poly(A) selection, and cDNA libraries were prepared via strand-specific RNA-seq with poly(A) selection (Illumina). Sequencing was performed on the Illumina NovaSeq platform (Illumina), generating 150 bp paired-end reads. Quality control and data analysis were conducted via a standard RNA-sequencing pipeline. The differential gene expression analysis was performed with DESeq2 (version 1.44.0, RRID: SCR_015687) to normalize counts and identify DEGs. The functional enrichment analysis of the DEGs (fold change > 2; Benjamini-Hochberg FDR < 0.05) was conducted with ShinyGO (version 0.80, http://bioinformatics.sdstate.edu/go/, RRID: SCR_019213), and the data were filtered to assess statistical significance (FDR < 0.05). Venn diagrams of the DEGs shared between iPSC-derived neurons, astrocytes, and microglia were generated via Draw Venn Diagram (accessed March 2023; https://bioinformatics.psb.ugent.be/webtools/Venn/). The enrichment of CDDO-Me-induced microglial DEGs within annotated human brain microglial clusters was assessed using Fisher's exact test against reference datasets from the Human Microglia Atlas [7].

**GSEA**

GSEA of senescence-associated pathways was performed with GSEA software (version v4.3.2, https://www.gsea-msigdb.org; RRID: SCR_003199) via the Molecular Signatures Database (MSigDB, version v6.2, RRID: SCR_016863). Briefly, RNA counts from control and CDDO-Me-treated samples were ranked based on the signal-to-noise ratio (SNR), which was calculated as follows: $(\mu\_treated - \mu\_control)/(\sigma\_treated + \sigma\_control)$. Genes were sorted by the SNR in descending order, and GSEA was used to calculate the enrichment score (ES), normalized ES (NES), and corresponding FDR. For this study, the default GSEA settings were used, with the permutation parameter set to "gene set" due to the small sample size (n<7) and the gene set size limited to 10–500 genes. The input data included RNA-sequencing read counts from all three cell types; a sample label mapping file; and gene sets related to senescence, SASP, and cell cycle regulation from MSigDB, along with the manually curated SenMayo SASP gene set (GSEA M45803) [8-13].

**Untargeted MS-based lipidomic analyses**

Lipidomic profiling of iPSC-derived microglia was performed by Lipotype GmbH (Dresden, Germany) using a high-resolution shotgun lipidomic platform. The cell pellets were flash-frozen, shipped on dry ice, and processed using a modified chloroform/methanol extraction protocol with internal standards specific to each lipid class [14]. Lipid extracts were analyzed on a Q Exactive Orbitrap mass spectrometer (Thermo Fisher) with a Triversa Nanomate automated nano-ESI source in both positive and negative ion modes. External mass calibration was performed weekly to ensure <5 ppm mass accuracy. Lipids were identified using LipotypeXplorer software based on MS1 and MS/MS data (https://www.science.org/content/product/lipotypexplorer). Quantification data were normalized to internal standards and reported in mol% formats. Lipid species with signal intensities ≥5-fold above the blank and noise thresholds were retained. High reproducibility (median CV ~4%) was confirmed in quality control samples (mammalian blood).

**Targeted LC-MS metabolomic analyses**

For the metabolomic analysis, 5 × $10^5$ iPSC-derived microglia were detached with Accutase, centrifuged, and immediately flash-frozen in a collection tube on dry ice. Six biological replicates per condition were analyzed, and the samples were stored at -80°C until analysis. For the metabolomic analysis, the extraction solution was composed of 50% methanol, 30% acetonitrile (ACN), and 20% water. The volume of the extraction solution was adjusted for the number of cells (1 mL per 1 × $10^7$ cells). After the addition of the extraction solution, the samples were vortexed for 5 minutes at 4°C and centrifuged at 16000 × g for 15 minutes at 4°C. The supernatants were collected and stored at −80°C until analysis. LC-MS analyses were conducted with a QExactive Plus Orbitrap mass spectrometer equipped with an Ion Max source and a HESI II probe coupled to a Dionex UltiMate 3000 UHPLC system (Thermo Fisher). External mass calibration was performed with a standard calibration mixture every seven days, as recommended by the manufacturer. The samples (5 µL) were injected onto a ZIC-pHILIC column (150 mm × 2.1 mm; i.d. 5 µm) with a guard column (20 mm × 2.1 mm; i.d. 5 µm) (Merck) for LC separation. Buffer A was 20 mM ammonium carbonate and 0.1% ammonium hydroxide (pH 9.2), and buffer B was ACN. The chromatographic gradient was run at a flow rate of 0.200 µL/minute as follows: 0–20 minutes, linear gradient from 80–20% of buffer B; 20–20.5 minutes, linear gradient from 20–80% of buffer B; 20.5–28 minutes, 80% buffer B. The mass spectrometer was operated in full scan, polarity switching mode with the spray voltage set to 2.5 kV and the heated capillary held at 320°C. The sheath gas flow was set to 20 units, the auxiliary gas flow was set to 5 units, and the sweep gas flow was set to 0 units. Metabolites were detected across a mass range of 75–1,000 m/z at a resolution of 35,000 (at 200 m/z), with the automatic gain control target set to $10^6$ and the maximum injection time to 250 ms. A lock mass technique was used to ensure mass accuracy below 5 ppm. The data were acquired with Xcalibur software v4.4 (RRID:SCR_014593, Thermo Fisher). The peak areas of the metabolites were determined with TraceFinder software v5.1 SP1(RRID:SCR_023045, Thermo Fisher) and identified by the exact mass of each singly charged ion and by the known retention time on the HPLC column. Data analysis was performed with MetaboAnalyst (version 6.0, https://www.metaboanalyst.ca/, RRID: SCR_016723). The statistical analysis module was utilized for PCA. The enrichment analysis module was employed to perform MSEA based on the Small Molecule Pathway Database (SMPDB; RRID: SCR_004844) with metabolite sets containing at least 2 entries. The joint pathway analysis module was used to integrate the metabolic pathway analysis results from the combined metabolomic and bulk RNA-sequencing data utilizing the parameters hypergeometric test for the enrichment analysis, degree centrality for the topology measure, and combined *P* values (unweighted) for the integration method.

**Single-cell transcriptome library preparation**

iPSC-derived cells in the triculture system were dissociated into single-cell suspensions by an incubation with Accutase for 15 minutes at 37°C. After dissociation, the cell suspensions were filtered through a 40 μm filter and resuspended in PBS containing 0.04% BSA at a final concentration of 1000 cells/μL, ensuring that more than 95% of the cells were viable. Single-cell RNA-sequencing libraries were generated with the 10X Chromium Next GEM Single Cell 3' Reagent Kit v3.1 (10x Genomics, Cat. # 1000128) according to the manufacturer's instructions. Libraries were pooled and subjected to paired-end sequencing on the Illumina NovaSeq 6000 platform (SP Flow Cell, Illumina) with a sequencing depth of 300 million reads per library at GENEWIZ GmbH (Leipzig, Germany).

**Single-cell sequencing data analysis**

The sequencing data were demultiplexed and filtered via the 10X Genomics Cell Ranger pipeline to generate filtered gene-barcode matrices, which were used as the input for the downstream analysis with the R package Seurat (Seurat version 4.1.0, RRID: SCR_007322; R version 4.3.3, RRID: SCR_001905) [15, 16]. A total of 7,160 cells were sequenced. Low-quality cells were filtered out (number of unique molecular identifiers (UMIs) < 2500, number of detected genes < 500, and mitochondrial DNA ratio < 0.2). Genes with a count of zero or expressed in fewer than ten cells were also removed. DoubletFinder was used to remove doublets from the dataset (DoubletFinder version 3, RRID: SCR_018771), and SCtransform (version 0.4.1, RRID: SCR_022146) was used to normalize the values, regressing out cell cycle reads and mitochondrial reads. The data were integrated via the Harmony package (version 1.2.0, RRID:SCR_022206). Principal components (PCs) were determined, and the first 40 PCs were used to cluster the cells via a K-nearest neighbor graph. By performing a UMAP analysis of the two integrated datasets followed by cluster analysis with a resolution of 0.8, 11 clusters across both conditions were identified. Nebulosa's kernel function (Nebulosa version 3.19, DOI: 10.18129/B9.bioc.Nebulosa) was used to visualize the density estimation for cell type-specific markers in the UMAP plot. Conserved markers were identified across all samples for each cluster with the FindConservedMarkers function in the Seurat package with the default settings (min.pct = 0.25, logfc.threshold = 0.25) and used to assign cell types to cell clusters.

The UCell package (version 2.7.6, RRID:SCR_027109 [17]), which is based on the Mann-Whitney U statistic, was used to evaluate UPR$^{mt}$ signatures (*HSPD1, HSPA9,* and *DDIT3*) in the single-cell datasets. Merging of the neuronal, astrocytic, and microglial clusters was performed after manual annotation to increase the statistical power of the differential expression (DE) analysis. DEGs between the conditions in each cluster were identified with the FindMarkers function in the Seurat package using the MAST test (padj < 0.05, logFC > 2), and *P* value adjustment was performed with Bonferroni correction based on the total number of genes in the dataset. Venn diagrams of the DEGs shared between iPSC-derived neurons, microglia, and astrocytes in the monoculture and triculture systems were generated via Draw Venn Diagram tool (https://bioinformatics.psb.ugent.be/webtools/Venn/). The functional enrichment analysis of the DEGs (fold change > 2; Benjamini-Hochberg FDR < 0.05) was conducted via ShinyGO (version 0.80, http://bioinformatics.sdstate.edu/go/, RRID: SCR_019213) by filtering the results for statistical significance (FDR<0.05) and using the KEGG PATHWAY Database (RRID: SCR_018145) and the Reactome database (RRID: SCR_003485) as the pathway databases. The CellChat package (version 2.1.0, RRID: SCR_021946) was used for the visualization and analysis of cell-cell communication events identified from the single-cell data, according to the developer's instructions. The corresponding annotation file was downloaded from https://raw.githubusercontent.com/hbctraining/scRNA-seq/master/data/annotation.csv.

**α-Synuclein pre-formed fibril treatment**

The production and purification of recombinant human α-synuclein followed the methodology described previously [2]. The α-synuclein cDNA was cloned and inserted into the pET-21d(+) DNA vector (Novagen, Merck Millipore, Cat. #69743; GenBank accession U13874), and this plasmid was expressed in *E. coli* BL21(DE3) cells (Novagen, Merck, Cat. # 69450, NCBI Taxonomy ID: 469008). After protein production, the bacteria were collected, and unwanted proteins were removed via centrifugation. Ion exchange chromatography on Q-Sepharose Hi-Trap columns equilibrated with Solution A (50 mM Tris, pH 7.4; Amersham Biosciences) was used to eliminate contaminating nucleic acids and proteins. The purified α-synuclein protein was eluted by applying a gradient of Solution B (50 mM Tris, pH 7.4, and 1 M KCl; Amersham Biosciences) to the soluble fraction. The α-synuclein-containing fractions were combined and further purified via Superose 12 column chromatography (Amersham Biosciences) in 20 mM HEPES (pH 7.4) with 100 mM KCl. For pre-formed fibril (PFF) production, α-synuclein monomers were incubated at 37°C with shaking at 1000 rpm for 7 days. α-Synuclein PFFs were validated via electron microscopy, a sedimentation assay (100,000 × g for 60 minutes), and a thioflavin-T assay (Sigma-Aldrich, Cat. # T3516). A PE/Atto 594 Conjugation Kit - Lightning-Link (Abcam, Cat. # ab269900) was used to label α-synuclein PFFs according to the manufacturer's instructions. For the treatment of iPSC-derived microglia, α-synuclein PFFs were diluted in PBS to 100 µg/mL and then sonicated (10-second pulses at a 30% amplitude, six times at 2-minute intervals) using a Q700-220 sonicator (Qsonica). α-Synuclein PFFs were then diluted in microglial medium and added to cultures to a final concentration of 1 µg/mL. The cells were treated with CDDO-Me, etoposide, or MCHA, as indicated. For immunostaining, the cells were fixed with 4% PFA for 10 minutes after 30, 60, and 90 minutes of treatment. Immunostaining was performed as described above. Images were acquired with a Leica TCS SP8 confocal microscope with a 60×/1.4 numerical aperture oil immersion objective (Leica). For live cell imaging, images were acquired with an Evident APX100-HCU box-type microscope with a 20×/0.8 numerical aperture objective (Evident). Images were acquired every 20 minutes for 3 hours. The particle analysis was performed via the Speckle Counting and Object Tracking pipelines in CellProfiler Image Analysis Software (version 4.2.6, RRID:SCR_007358) according to the suggested guidelines, and CD68 staining or brightfield imaging was used to detect the cell body.

#### Supplemental Tables

#### Supplemental Table 1: List of antibodies

##### Primary antibodies

| Target | IF/IHQ | WB | Host | Company | Cat.Number | RRID |
|---|---|---|---|---|---|---|
| Anti-ß-Amyloid 1-16 | 1/250 | | Mouse | Biolegend | 803004 | RRID:AB_2715854 |
| ß-Actin | | 1/5000 | Mouse | Santa Cruz | sc-47778 | RRID:AB_626632 |
| ATF5 | | 1/2000 | Rabbit | Abcam | ab184923 | RRID: AB 2800462 |
| PECAM-1 (CD31) | 1/50-1/500 | | Mouse | Santa Cruz | sc-376764 | RRID:AB_2801330 |
| CD68 Monoclonal Antibody (KP1) anti-human | 1/500 | | Mouse | Thermo Fisher Scientific | MA5-13324 | RRID:AB_10987212 |
| C1qA | 1/1000 | | Rabbit | Proteintech | 11602-1-AP | RRID: AB_2067153 |
| CTIP2 | 1/500 | | Rat | Abcam | 18465 | RRID:AB_2064130 |
| GATA-2 (H-116) | 1/100 | | Rabbit | Santa Cruz | sc-9008 | RRID:AB_2294456 |
| GFAP | 1/500 | 1/1000 | Rabbit | Agilent | Z0334 | RRID:AB_10013382 |
| Histone H2A.X | 1/500 | | Mouse | Santa Cruz | sc-517348 | RRID:AB_2783871 |
| HSP60 (B-9) | 1/200 | 1/2000 | Mouse | Santa Cruz | sc-271215 | RRID:AB_10607973 |
| IBA1 (IHC) | 1/500 | | Rabbit | Wako | 019-19741 | RRID:AB_839504 |
| IBA1 | 1/500-1/1000 | | Rabbit | Wako | 016-20001 | RRID:AB_839506 |
| LAMP1 | 1/100 | | Mouse | DSHB | H4A3 | RRID: AB_2296838 |
| LONP1 | | 1/5000 | Rabbit | Proteintech | 15440-1-AP | RRID: AB_2137152 |
| LC3B | | 1/500 | Rabbit | Proteintech | 18725-1-AP | RRID: AB_2137745 |
| MAP2 | 1/1000-1/5000 | | Chicken | Millipore | AB 5622 | RRID:AB_91939 |
| OTX2 | 1/400 | | Goat | Neuromics | GT15095 | RRID:AB_2157174 |
| Human CDKN2A | 1/500 | | Mouse | Santa Cruz | sc-56330 | RRID:AB_785018 |
| p21 WAF1 / CIP1 | 1/500 | | Rabbit | Cell Signaling Technology | 2947S | RRID:AB_823586 |
| P62 | 1/1000 | 1/1000 | Rabbit | Sigma Aldrich | P0067 | RRID: AB_1841064 |
| PAR | | 1/500 | Mouse | AdipoGen Life Sciences | AG-20T-0001-C050 | RRID:AB_2490281 |
| PAX6 | 1/500 | | Rabbit | Biolegend | 901301 | RRID:AB_2565003 |
| PHF Tau Thr181 | 1/1000 | 1/1000 | Mouse | Thermo Fisher Scientific | MN1050 | RRID:AB_223651 |
| PITRM1 | | 1/1000 | Rabbit | Genetex | GTX119739 | RRID:AB_10617717 |
| pS129 Synuclein | | 1/1000 | Rabbit | Genetex | GTX50222 | RRID:AB_11179523 |
| PSD95-specific, DLG4 | 1/250 | | Rabbit | Proteintech | 20665-1-AP | RRID: AB_2687961 |

| | | | | | | |
|---|---|---|---|---|---|---|
| Anti-SPI1 Pu.1 (7C6B05) | 1/500 | | Mouse | Biolegend | 658002 | RRID:AB_2562720 |
| S100ß | 1/100 | | Mouse | Sigma Aldrich | S2532 | RRID:AB_477499 |
| SOX2 | 1/500 | | Mouse | Abcam | ab75485 | RRID:AB_1278243 |
| α-synuclein | | 1/1000 | Mouse | BD Bioscience | 610787 | RRID:AB_398108 |
| TBR1 | 1/500 | | Rabbit | Abcam | 31940 | RRID:AB_2200219 |
| Human TOMM20 | 1/200 | 1/1000 | Mouse | Santa Cruz | sc-17764 | RRID:AB_628381 |
| Total Tau | | 1/1000 | Mouse | Thermo Fisher Scientific | MN1000 | RRID: AB_2314654 |
| ß-Tubulin | 1/1000 | | Mouse | Biolegend | 802001 | RRID:AB_2564645 |
| Vinculin | | 1/5000 | Mouse | Santa Cruz | sc-73614 | RRID:AB_1131294 |

#### Secondary antibodies

| **Name** | **IF/IHQ** | **WB** | **Host** | **Company** | **Cat.Number** | **RRID** |
|---|---|---|---|---|---|---|
| Anti-Rabbit IgG (H+L) Alexa Fluor™ 488 | 1/1000 | | Goat | Invitrogen | A-11008 | RRID:AB_143165 |
| Anti-Rabbit IgG (H+L) Alexa Fluor™ 568 | 1/1000 | | Goat | Invitrogen | A-11011 | RRID:AB_143157 |
| Anti-Mouse IgG (H+L) Alexa Fluor™ 488 | 1/1000 | | Goat | Invitrogen | A-11001 | RRID:AB_2534069 |
| Anti-Mouse IgG (H+L) Alexa Fluor™ 568 | 1/1000 | | Goat | Invitrogen | A-11004 | RRID:AB_2534072 |
| Anti-Chicken IgG (H+L) Alexa Fluor™ 647 | 1/1000 | | Goat | Invitrogen | A-21449 | RRID:AB_2535866 |
| Anti -Rat IgG (H+L) Alexa Fluor™ 568 | 1/1000 | | Goat | Abcam | Ab175476 | RRID:AB_2813739 |
| Anti -mouse IgG, HRP - linked Antibody | | 1/5000 | Horse | Cell Signaling | Cat# 7076 | RRID:AB_330924 |
| Anti -rabbit IgG, HRP - linked Antibody | | 1/5000 | Goat | Cell Signaling | Cat# 7074 | RRID:AB_2099233 |

**Supplemental Table 2: Primer sequences.**

| Primer | Sequence |
|---|---|
| ATF4 FW | 5'-GTC CCT CCA ACA ACA GCA AG-3' |
| ATF4 RV | 5'- CTA TAC CCA ACA GGG CAT CC-3' |
| LONP1 FW | 5'- CCC GCG CTT TAT CAA GAT T-3' |
| LONP1 RV | 5'- AGA AAG ACG CCG ACA TAA GG-3' |
| HSP60 RNA FW | 5'- TGA CCC AAC AAA GGT TGT GA-3' |
| HSP60 RNA RV | 5'- CAT ACC ACC TCC CAT TCC AC-3' |
| RPLP0 FW | 5'-CCT CAT ATC CGG GGG AAT GTG-3' |
| RPLP0 RV | 5'- GCA GCA GCT GGC ACC TTA TTG-3' |
| HSPA9 FW | 5'- GGA AGC TGC TGA AAA GGC TA-3' |
| HSPA9 RV | 5'- CTT GGG TCC AGA AGA ATC CA-3' |
| DDIT3/chop FW | 5'- AGC CAA AAT CAG AGC TGG AA-3' |
| DDIT3/chop RV | 5'- TGG ATC AGT CTG GAA AAG CA-3' |
| CLPP RNA FW | 5'- CTC TTC CTG CAA TCC GAG AG-3' |
| CLPP RNA RV | 5'- GGA TGT ACT GCA TCG TGT CG-3' |
| CDKN2A FW | 5'-GAAGGTCCCTCAGACATCCCC-3' |
| CDKN2A RV | 5'-CCCTGTAGGACCTTCGGTGAC-3' |
| CXCL8 FW | 5'-GCTCTGTGTGAAGGTGCAGT-3' |
| CXCL8 RV | 5'-TGCACCCAGTTTTCCTTGGG-3' |
| mtND1 FW | 5'-CCACCTCTAGCCTAGCCGTTTA-3' |
| mtND1 RV | 5'-GGGTCATGATGGCAGGAGTAAT-3' |
| mtCYB FW | 5'-ATCACTCGAGACGTAAATTATGGCT-3' |
| mtCYB RV | 5'-TGAACTAGGTCTGTCCCAATGTATG-3' |
| ATF5 FW | 5'-TGGCTCGTAGACTATGGGAAA-3' |
| ATF5 RV | 5'-ATCAACTCGCTCAGTCATCCA-3' |
| BCL2 FW | 5'-GGTGGGGTCATGTGTGTGG-3' |
| BCL2 RV | 5'-CGGTTCAGGTACTCAGTCATCC-3' |
| BAX FW | 5'-CCCGAGAGGTCTTTTTCCGAG-3' |
| BAX RV | 5'-CCAGCCCATGATGGTTCTGAT-3' |
| BAK FW | 5'-GTTTTCCGCAGCTACGTTTTT-3' |
| BAK RV | 5'-GCAGAGGTAAGGTGACCATCTC-3' |
| CASP3 FW | 5'-CATGGAAGCGAATCAATGGACT-3' |
| CASP3 RV | 5'-CTGTACCAGACCGAGATGTCA-3' |
| CASP8 FW | 5'-GTTGTGTGGGGTAATGACAATCT-3' |
| CASP8 RV | 5'-TCAAAGGTCGTGGTCAAAGCC-3' |
| CASP9 FW | 5'-CTCAGACCAGAGATTCGCAAAC-3' |
| CASP9 RV | 5'-GCATTTCCCCTCAAACTCTCAA-3' |
| TOMM20 FW | 5'-GGTACTGCATCTACTTCGACCG-3' |
| TOMM20 RV | 5'-TGGTCTACGCCCTTCTCATATTC-3' |
| TOMM70 FW | 5'-TGTTTTGCATTGTACCGCCAG-3' |
| TOMM70 RV | 5'-TAGTGCATAGCCTTCGGCAC-3' |
| TIMM21 FW | 5'-ACTTTTCTACGAGCCGTACAGT-3' |
| TIMM21 RV | 5'-TGTTAAGCACGATGTATGGCAAA-3' |
| TIMM23 FW | 5'-GTCCCGCTAACTGGTATGAAC-3' |

| | |
|---|---|
| TIMM23 RV | 5'-CGTAAAGAAGGCCAGCTCAAATC-3' |
| TNFa FW | 5'- CAC TT GGA GTG ATC GGC C-3' |
| TNFa RV | 5'- CTC AGC TTG AGC GTT TGC TAC |
| IL1B FW | 5'- TCC CCA GCC CTT TTG TTG A-3' |
| IL1B RV | 5'- TTA GAA CCA AAT GTG GCC GTG-3' |
| IL6 FW | 5'- GGT ACA TCC TCG ACG GCA TCT-3' |
| IL6 RV | 5'- GTG CCT CTT TGC TGC TTT CAC-3' |
| CDKN1A RNA FW | 5'- GGC AGA CCA GCA TGA CAG ATT-3' |
| CDKN1A RNA RV | 5'- GCG GAT TAG GGC TTC CTC T-3' |
| NDUFV1 FW | 5'- GGG TAT CTG TGC GTT TCA GC-3' |
| NDUFV1 RV | 5'- GGT TGG TGA AAA TCC GGT CT-3' |
| NDUFS2 FW | 5'- AGC AAA GAA ACA GCC CAC TG-3' |
| NDUFS2 RV | 5'- GGC CCA AAG TTC AGG GTA AT-3' |
| Actin FW | 5'- TGA CCC AGA TCA TGT TTG AGA-3' |
| Actin RV | 5'- AGT CCA TCA CGA TGC CAG T-3' |

**Supplemental Unprocessed Blots**

**Supplementary Figure 4D**

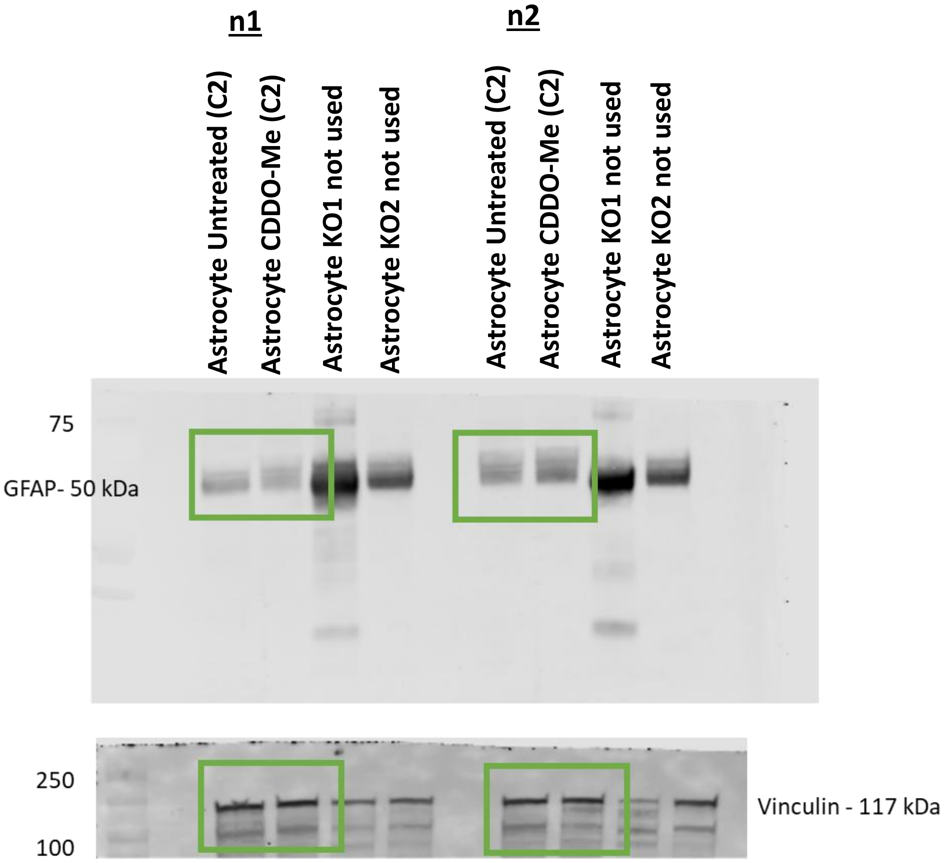

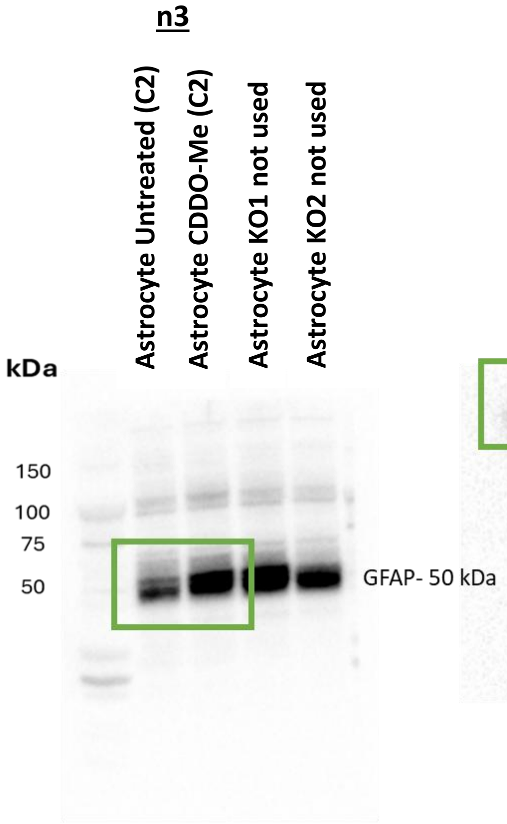
n3
Astrocyte Untreated (C2)
Astrocyte CDDO-Me (C2)
Astrocyte KO1 not used
Astrocyte KO2 not used
kDa
150
100
75
50
GFAP- 50 kDa

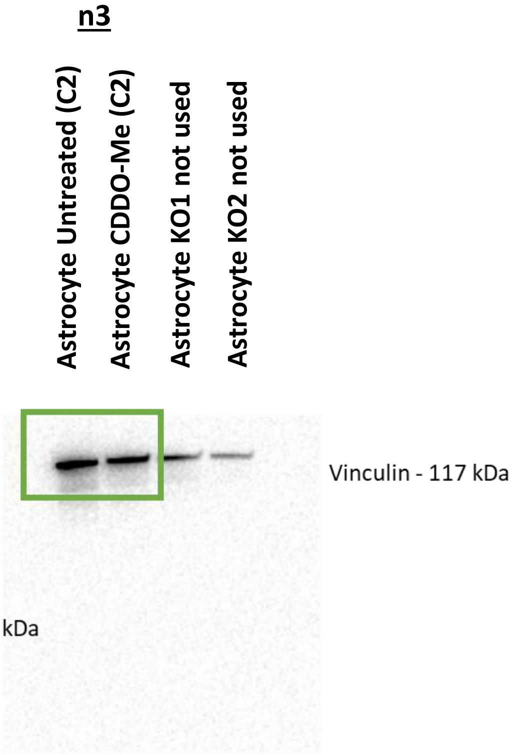
n3
Astrocyte Untreated (C2)
Astrocyte CDDO-Me (C2)
Astrocyte KO1 not used
Astrocyte KO2 not used
Vinculin - 117 kDa

**Supplementary Figure 4F**

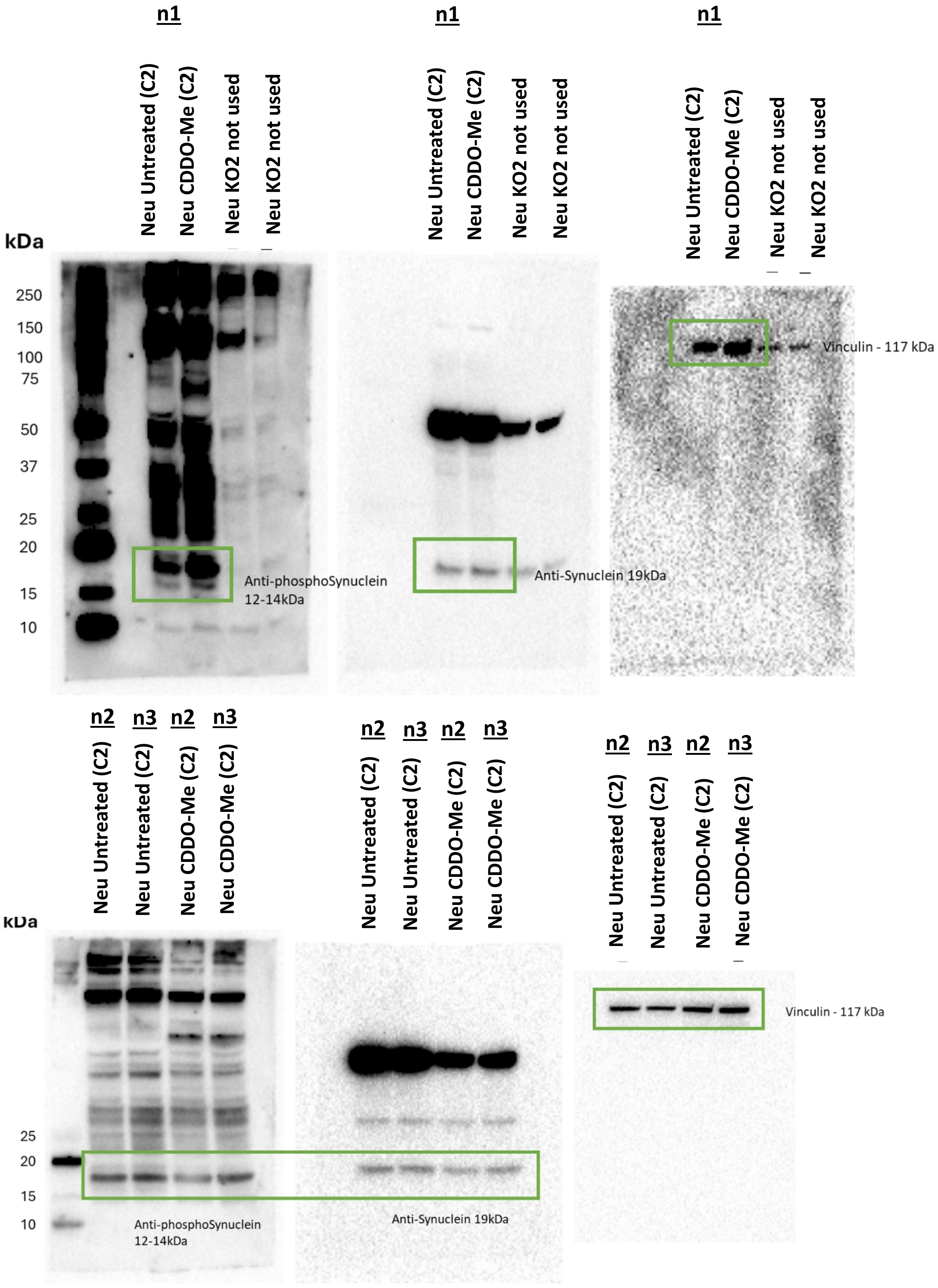

#### Supplementary Figure 4F

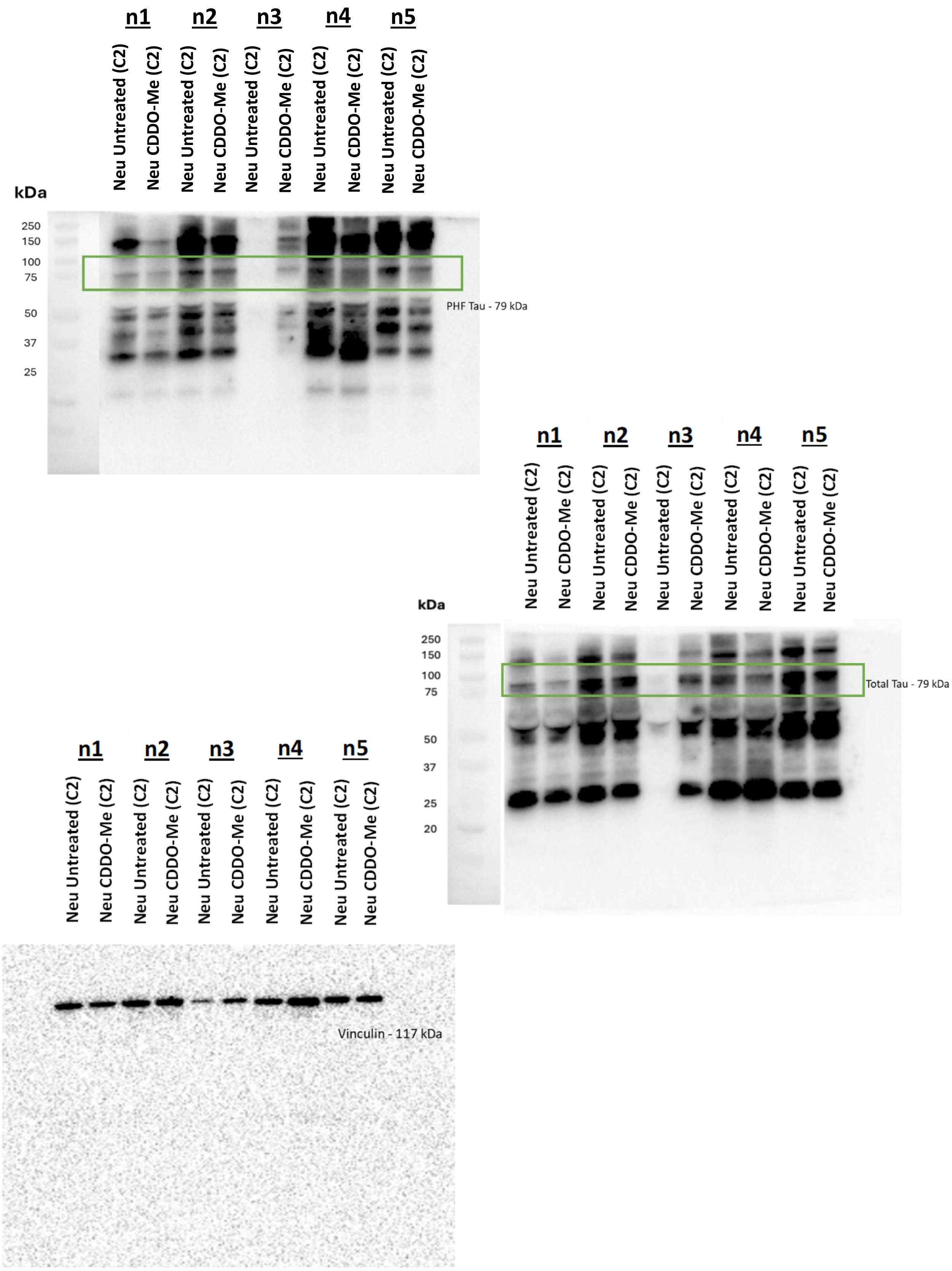

#### Supplementary Figure 6H

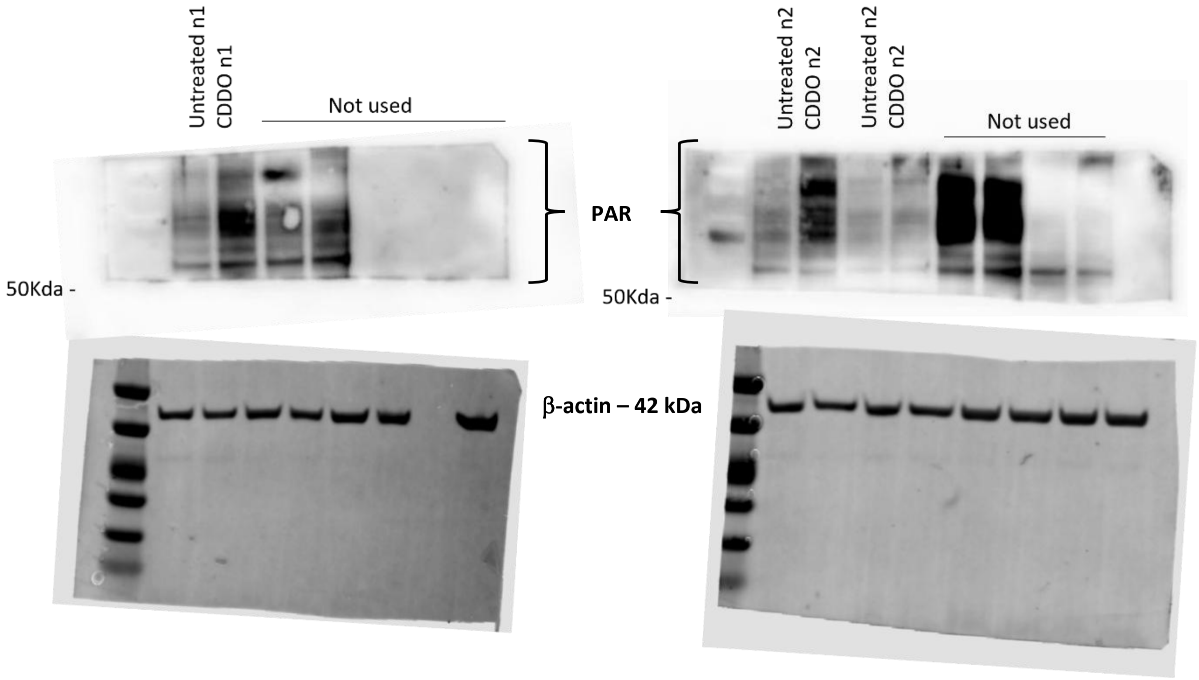

**Supplementary Figure 7A**

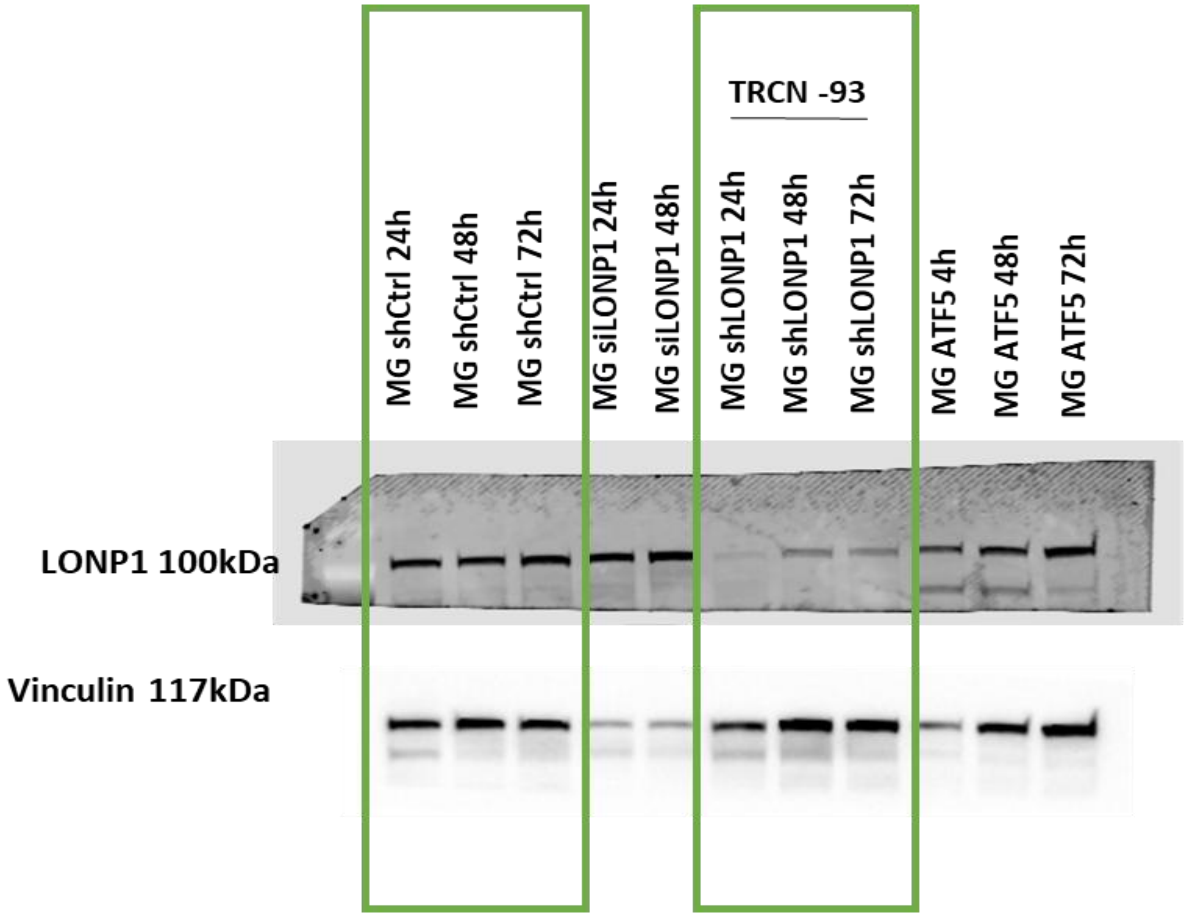

**Supplementary Figure 7F**

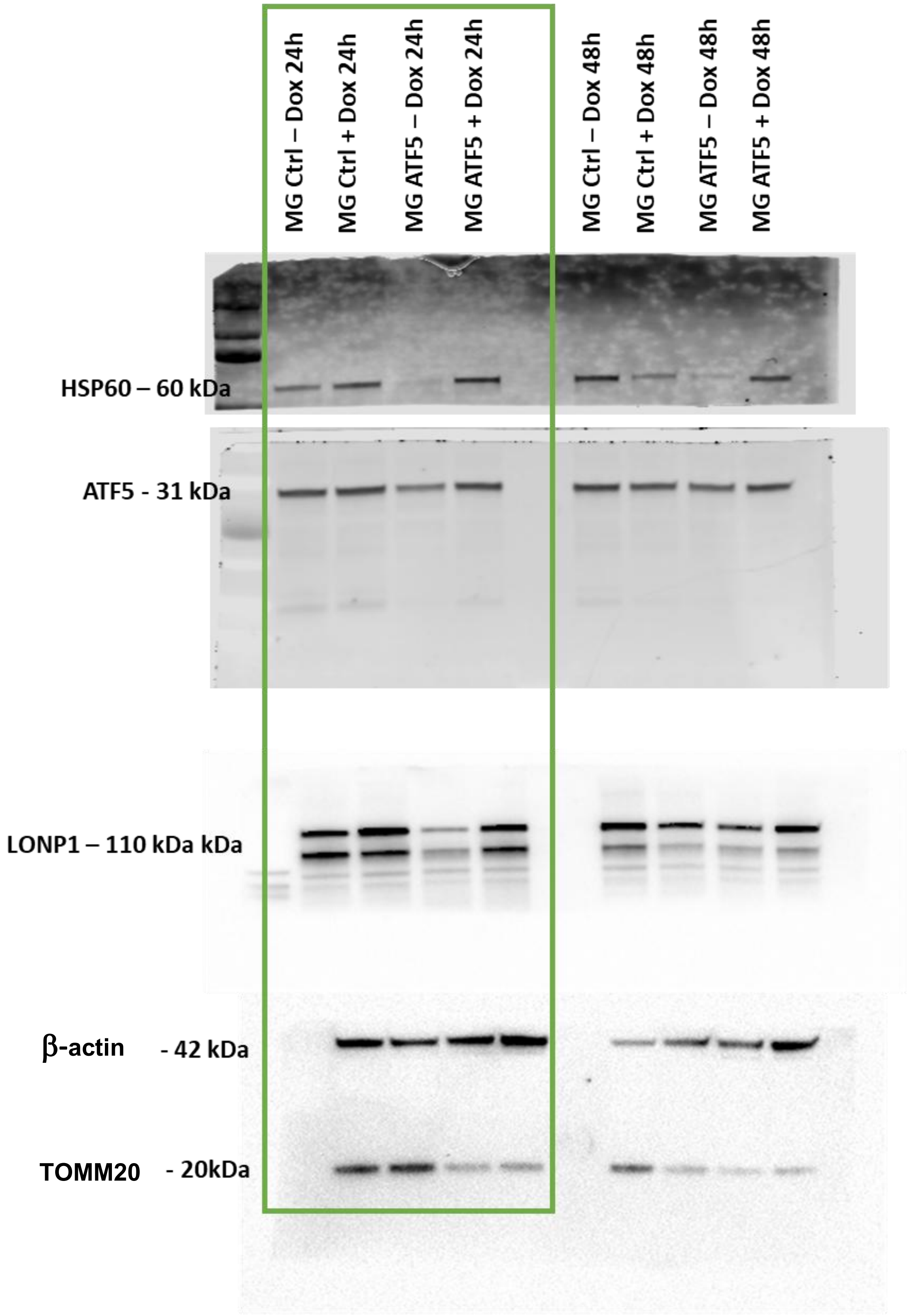

**Supplementary Figure 9A**

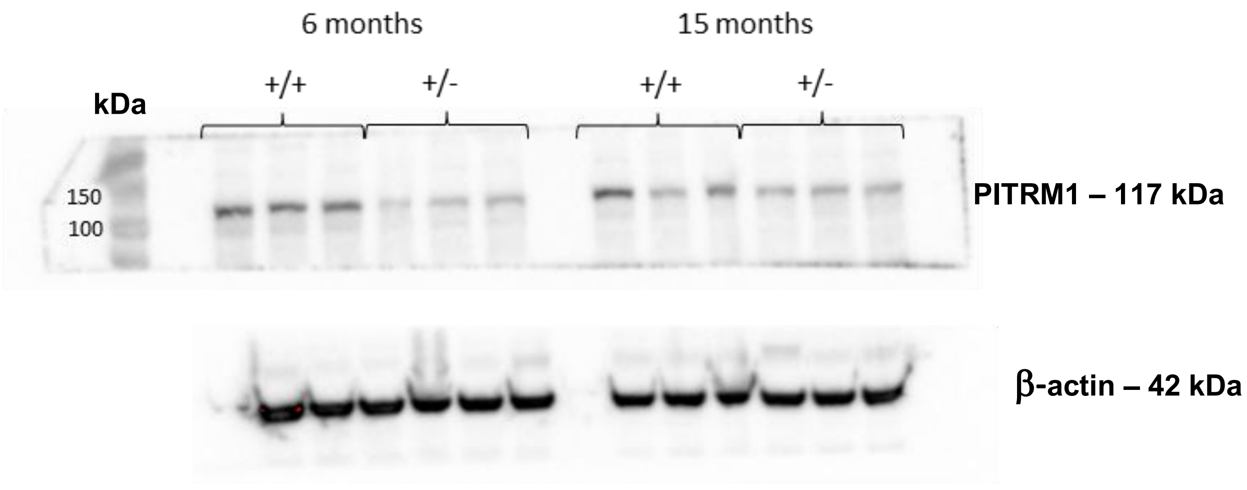

Supplementary Figure 14H

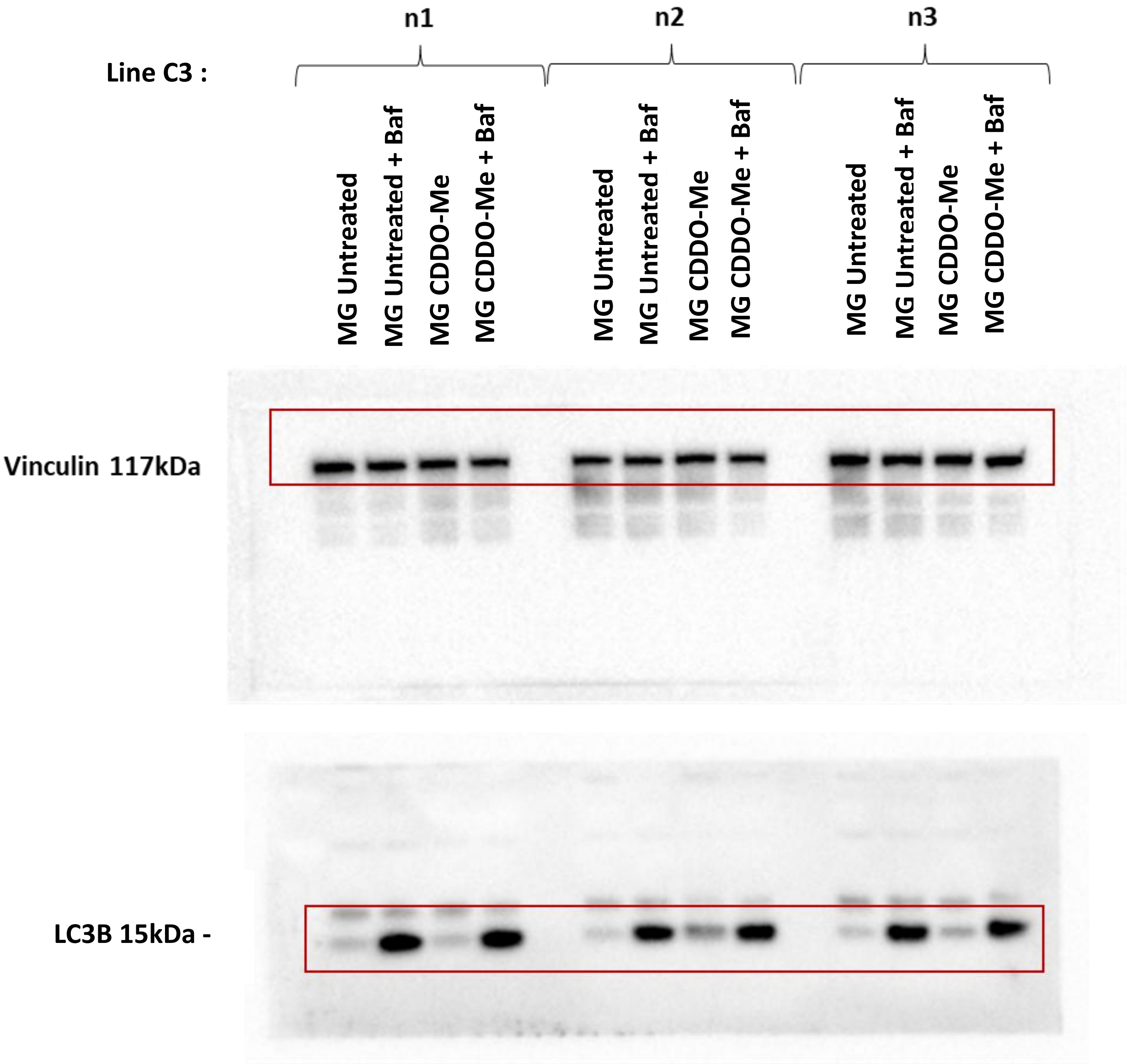

#### Supplementary Figure 16E

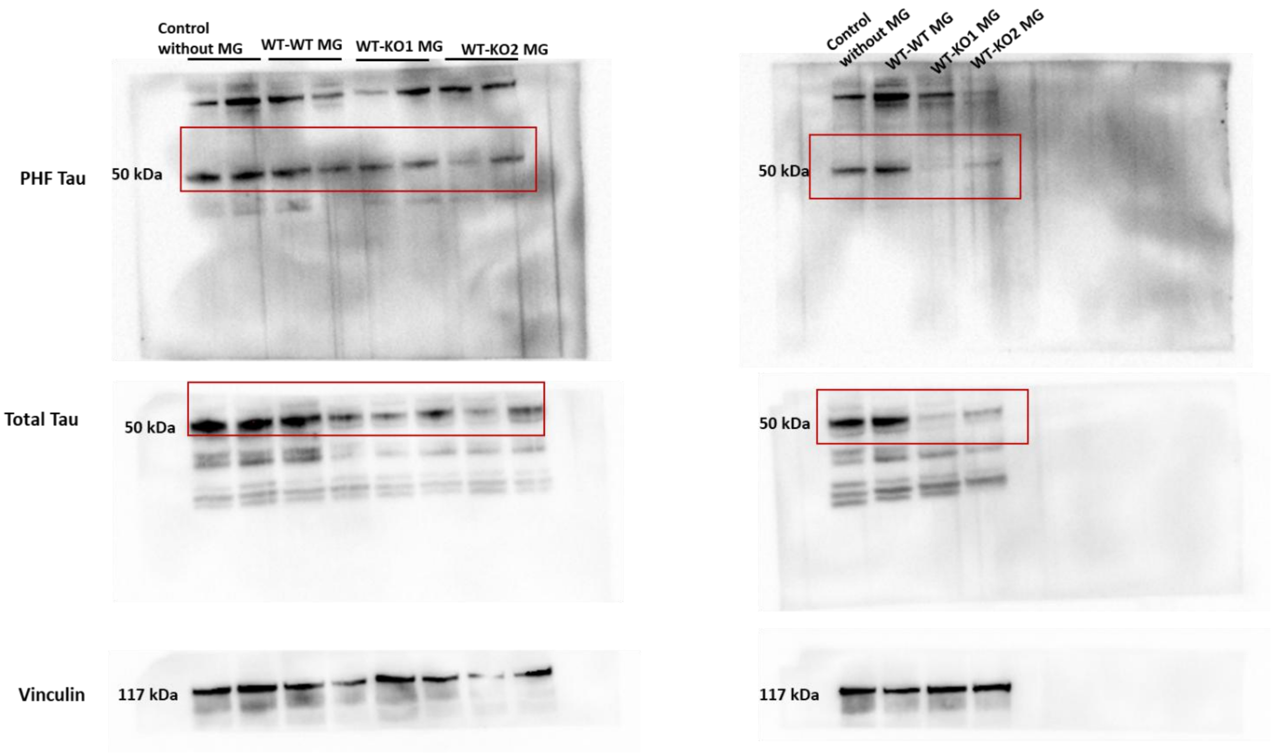
